# Genome-wide analyses point to differences in genetic architecture of BMI between tall and short people

**DOI:** 10.1101/2020.09.24.312181

**Authors:** Bochao D. Lin, Benjamin H. Mullin, Scott G. Wilson, John P. Walsh, Yue Li, Roger Adan, Jurjen J. Luykx

## Abstract

To examine differences in the genetic architecture of BMI between tall and short people, we conducted genome-wide and follow-up analyses using UK Biobank data. We identify 57 loci as height-specific, detect differences in SNP-based heritability between tall and short people and show how genetic correlations between the two rises during the lifespan. Using phenome-wide analyses (PHEWAS), a significant association between a short people-specific locus on *MC4R* and energy portion size was detected. We identify one locus (*GPC5*-*GPC6)* with different effect directions on BMI in short and tall people. PHEWAS indicates this locus is associated with bone mineral density. Transcriptome-wide analyses hint that genes differentially associated with BMI in short vs tall people are enriched in brain tissue. Our findings highlight the role of height in the genetic underpinnings of BMI, provide biological insight into mechanisms underlying height-dependent differences in BMI and show that in short and tall people obesity is a risk factor that differentially increases susceptibility for disease.

## Introduction

Body mass index (BMI) is defined as weight in kilograms divided by the square of height in meters (kg/m^2^), which has been a universal measure to classify obesity (BMI> 30) in epidemiology and clinical practice since the 19th century [1]. Meta-analysis of BMI in a twin study shows significant BMI heritability differences in different age groups: BMI heritability increases with age before adulthood and then decreases [2]. This decrease is likely due to the larger influence of environmental factors over time. Recently, genome-wide association studies (GWAS) have demonstrated sex-differences in the genetic architecture of BMI. For example, the GIANT consortium has identified novel loci for BMI in GWAS of sex-specific cohorts in European populations: genetic variants nearby *ASB4* and *LOC284260* for women, and genetic variants nearby *USP37* and *ZEYB10* for men [3]. Another large GWAS of BMI in a sex-/age-specified cohort identified 15 loci with significantly different effects on BMI in younger versus older individuals (split by median age: 50)[4]. Thus, BMI heritability differs across age groups, ethnicity and sex.

A GWAS of BMI adjusting for height in young children yielded marked differences relative to conventional BMI GWASs not parsed by height, and identified a genetic locus on *ADCY3* for height-adjusted BMI [5]. Importantly, a GWAS of BMI accounting for height in adults is currently lacking. Recent research shows a negative association between BMI and height: BMI tends to be higher among shorter adults, especially in women [6]. Disturbed control in regulation of energy balance and environmental factors may differentially affect tall and short people in the risk of becoming obese. With fixed food portion sizes served to short and tall individuals, shorter adults are at increased risk for obesity, simply because their lean mass is lower and they require less energy for metabolism. In other words, short people may be more susceptible to becoming obese through environmental factors, such as exposure to food. Therefore, height may be an important factor to consider when analyzing the genetic architecture of BMI.

Here, we hypothesized that the genetic architecture of short people differs from that of taller people. We expected that GWAS of BMI stratified on adults’ height would help identify novel obesity-associated genetic variants and deepen the understanding of which genes impact obesity in humans. We therefore conducted a GWAS of BMI using the UK Biobank (UKB) cohort to identify BMI-associated genetic variants specific for height. In addition, we conducted several post-GWAS analyses to interpret the results, including SNP-based heritability estimates, conditional analysis using Multi-trait-based conditional and joint analysis (mtCOJO), transcriptome-wide association analysis (TWAS), phenome-wide association studies (PHEWAS) of the loci, Mendelian Randomization (MR) and Summary-data-based Mendelian Randomization analysis (SMR). Using these methods, we provide consistent and converging evidence for differences in genetic architecture between short and tall people.

## Results

### Phenotype-level analyses of energy intake in height deciles

We analyzed data from 502,538 participants of the UKBB with a mean age of 57.0 (standard deviation (SD) 8.1), of whom 54.6% were female. We observed that mean BMI decreases with increasing height in both men and women **(Supplementary Figure 1)**. Although fat percentages were stable for most female height groups, the fat percentage decreased with increasing height in men (**Supplementary Figure 2**). Regarding energy intake (in KJ) and portion energy intake (energy intake divided by individuals’ weight-(KJ/Kg)), mean energy intake went up with increasing height in both men and women **(Figure 1A)**, which is expected as lean mass increases with increased height. However, confirming our hypothesis, we found that portion energy intake (corrected for kg body weight) decreased with increasing height **(Figure 1B)**, which shows that, when corrected for body weight, shorter individuals eat relatively more calories. In addition, energy intake increased with increasing body fat percentage (**Supplementary Figure 3A**), and portion energy intake decreased with increasing body fat percentage (**Supplementary Figure 3B**). We also calculated Pearson correlations between BMI, height, fat percentage, and height percentage in height decile groups **(Supplementary Figure 4)**. The positive correlations of BMI, fat percentage and waist hip ratio (WHR) were estimated in height decile groups. A negative correlation between height and BMI (r= −0.012, p <2.2 × 10^−16^ in total population), and height with fat percentage (r= −0.495, p<2.2 × 10^−16^ in total population) was observed **(Supplementary Figure 4)**. Importantly, all phenotypic correlations in height decile groups were different relative to the entire UKB study population. To summarize, we thus provide consistent phenotype-level evidence across height deciles in both men and women for our hypothesis that relative energy intake (energy intake divided by weight) is indeed less in tall people relative to short people.

**Figure 1.**
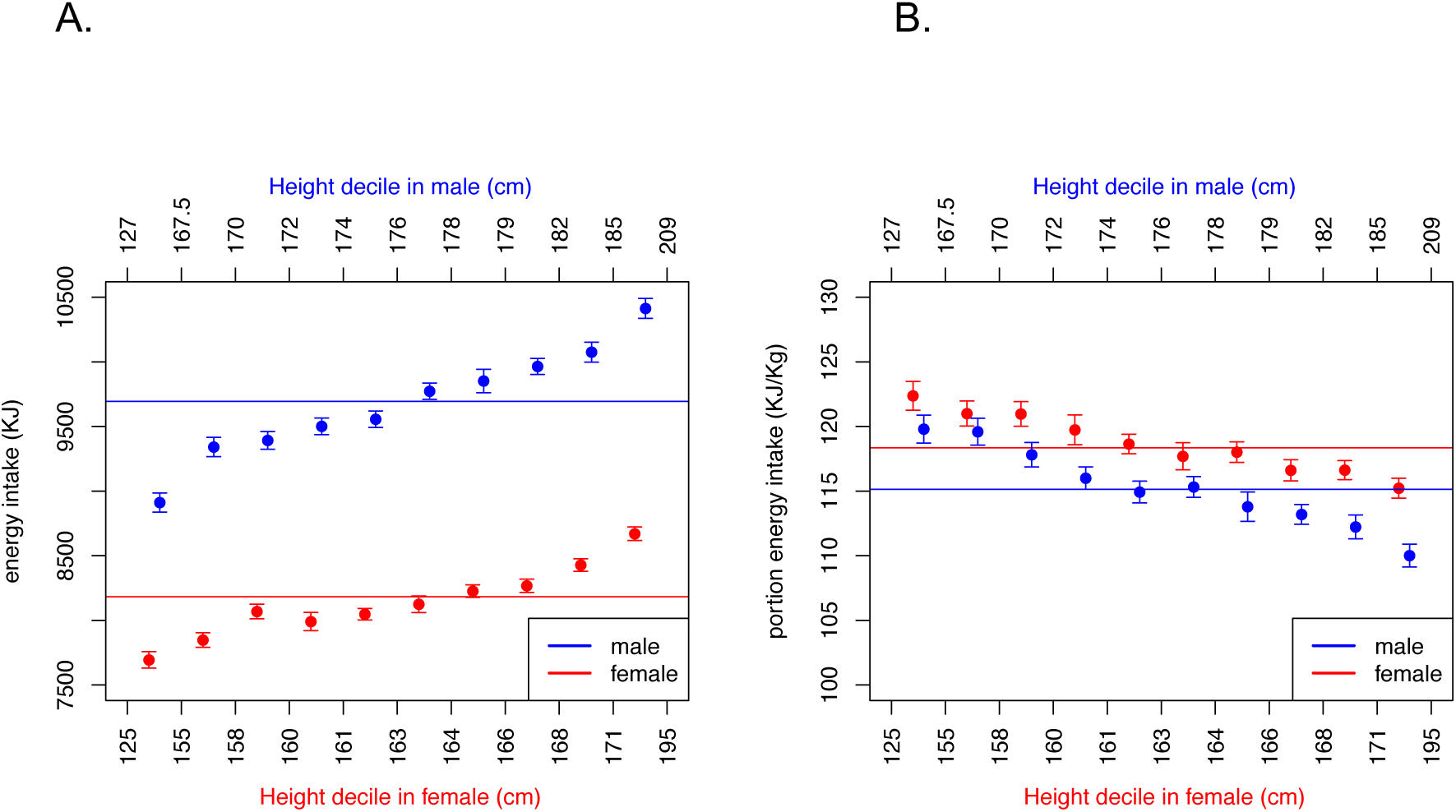
Energy intake and portion energy intake distributions in height decile groups. The red and blue lines represent the mean energy intake (in KJ; left panel, A) and portion energy intake (energy intake divided by individuals’ weight, in KJ/Kg; right panel, B). The dots and error bars represent the means and standard errors, respectively. Note that energy intake increases with increasing height in both men and women, while portion energy intake decreases with increasing height.

### Genome-wide association analyses (GWAS)

We analyzed whole-genome common variant data from 502,538 participants of the UKBB. We randomly selected 2/3 of participants (n=237,802) for our discovery dataset, and 1/3 of samples (n=121,598) for our replication dataset. We defined short individuals as those at the < 33% quantile of height (173cm for men, 160cm for women; **Supplementary Figures 5 and 6**, n = 71,144 in the discovery dataset, n=36,761 in the replication dataset) and persons at the > 67% quantile of height (179cm for men, 165cm for women; **Supplementary Figure 6**; n =75,199 in the discovery dataset, n=38,089 in the replication dataset) as the tall group. We ran GWASs of BMI on 7,904,644 SNPs that passed quality control checks (MAF≥0.01, imputation quality R^2^> 0.8) using PLINK2 ^[7]^ in linear regression models. As a proof of concept, we first performed GWAS of BMI in the entire study population and corroborated previous findings (**Supplementary Figure 7**) [8]. Then we performed GWAS meta-analysis using discovery and replication cohorts, of BMI in short people and in the tall group separately. We found 45 loci to be significantly associated with BMI in short people and 51 loci to be significantly associated with BMI in tall people (**Supplementary Figure 8)**. 25 genome-wide significant loci overlapped between short and tall people group. Thus, 46 unique loci were identified as potentially height-dependent.

### SNPs with different effects in the two height groups

To discover SNPs with different BMI effects in the two height groups (**Supplementary Table 1**), we selected BMI-associated SNPs based on two criteria: 1) *SNPs with P-value differences*: SNPs that are associated at P<5×10^−8^ with BMI in short or tall subjects and in addition to that with P values that are ≥ 10^5^ fold different (absolute values of log10 (P-short)-log10(P-tall) > 5) between GWAS-short and GWAS-tall (**Supplementary Figure 8 A**); 2) *SNPs with Z scores (effect size/ standard error) differences*, defined as the SNPs with absolute values of log2(Z - short) – log2(Z-tall) or vice versa > 1 (**Supplementary Figure 8 B**). The independent SNP list (n=57; 27 loci for short people and 31 loci for tall people) resulting from both criteria is shown in **Supplementary Table 2**. For category 1), we identified 23 loci associated with BMI in short people and 27 loci associated with BMI in tall people. For category 2), we identified 21 loci (6 loci for short people and 15 loci for tall people). The LocusZoom plots for these SNPs are shown in **Supplementary data file (PDF file)**. There were 15 independent SNPs meeting both criteria (3 loci were specific for short people and 12 loci were specific for tall people; **Supplementary Figure 8 C)**. More specifically, the loci nearby *USP33, KNTC1, SAE1, UPK3B* had both larger effects sizes and significantly smaller GWAS p-values for BMI in short people compared to tall people. The loci at 2p25.3, 3q25.2, 7q31.1 (*MON1A, XKR6, ADCY9, ETV5, BDNF, RPTOR, PACSIN1* and *ADARB1*) had both larger effects sizes and significantly smaller GWAS P values for BMI in tall compared to short people.

**Table 1.** SNP-based heritability and genetic correlations between BMI-short with BMI-tall in different age groups. Genetic correlation differences between short and tall individuals decrease with advancing age.

| Age / height | Short (M: < 173 cm; F: < 160 cm) | Tall (M: > 179 cm; F: > 165 cm) | Genetic correlation estimates |
| --- | --- | --- | --- |
| Young (age <54 years) | H<sup>2</sup>=0.27<br>SE=0.018<br>N=18592 | H<sup>2</sup>=0.30<br>SE=0.015<br>N=22383 | R<sub>g</sub>=0.9350<br>SE=0.0430 |
| Middle (54 ≤ age ≤ 61) | H<sup>2</sup>=0.24<br>SE=0.016<br>N=22898 | H<sup>2</sup>=0.28<br>SE=0.015<br>N= 23348 | R<sub>g</sub>=0.9799<br>SE=0.0505 |
| Old (age > 61) | H<sup>2</sup>=0.21<br>SE=0.013<br>N=29950 | H<sup>2</sup>=0.24<br>SE=0.013<br>N=29468 | R<sub>g</sub>=1.0000<br>SE=0.0597 |

### Multi-trait-based conditional and joint analysis (mtCOJO)

Using mtCOJO, we conditioned short people’s BMI GWAS results on tall people’s BMI GWAS results (**Figure 2 A**). There were two SNPs in the same LD locus on 13q31.3 showing significant associations (rs80285134 and rs117075592; **Supplementary Figure 9**). The top SNP was rs80285134 (MAF = 0.015, imputation r^2^ = 0.95), an intergenic variant (between *GPC5* and *GPC6*), which is located in an open chromatin region (so likely a regulatory variant). *GPC6* is associated with bone mineral density and plays a key role in osteoporosis [9]. We found no association of this SNP with height in public GWASs. The second SNP in this locus was rs117075592 (MAF=0.014, imputation r^2^=0.94), an intronic variant located 22kb from rs80285134, with which it exhibits a high degree of LD (r2= 0.854, D’ >0.9).

**Figure 2A.**
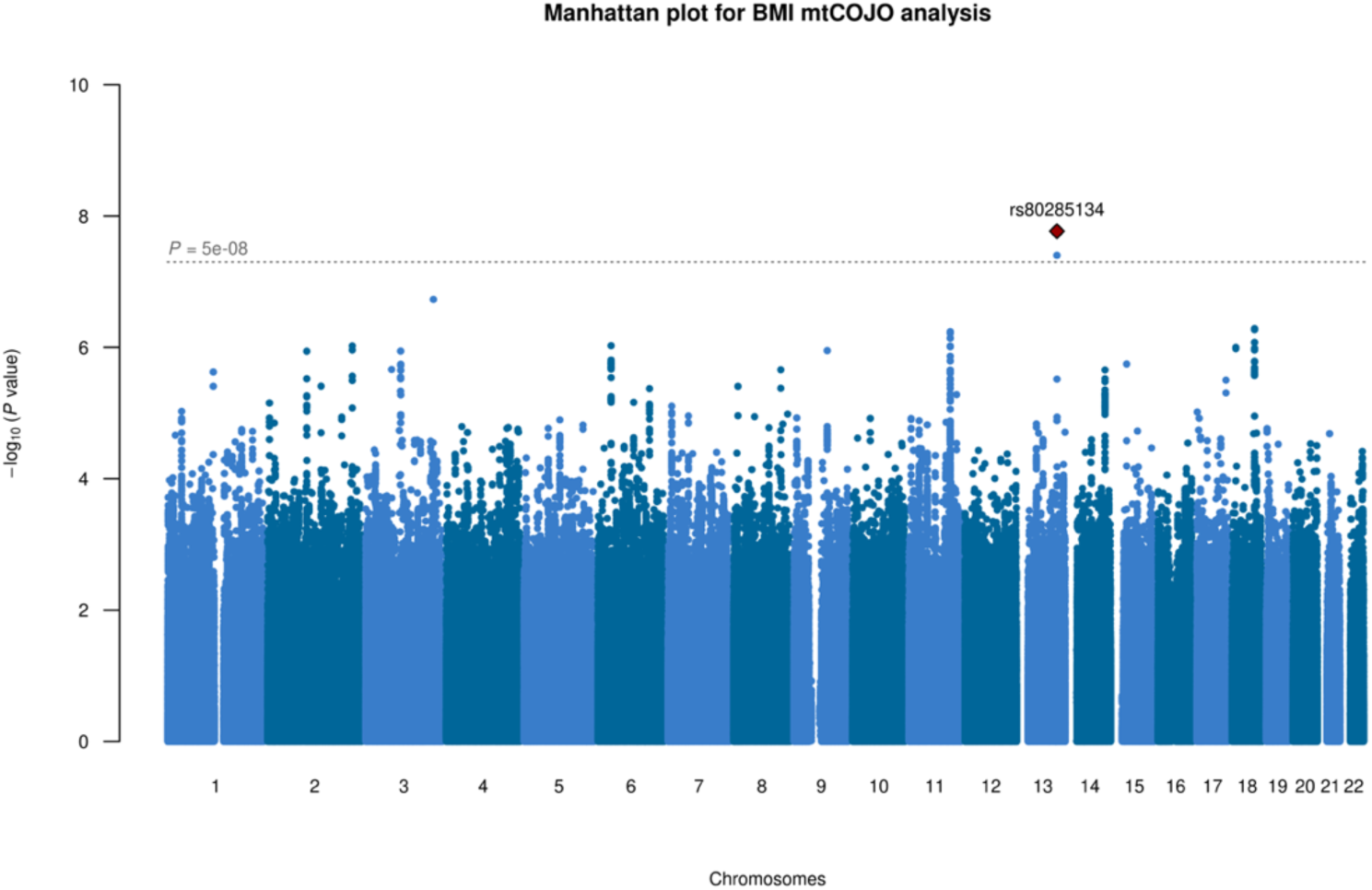
mtCOJO results of short people’s BMI GWAS results conditioned on tall people’s BMI GWAS results.

In checking the association of rs80285134 in our GWAS of BMI in short and tall individuals we had conducted (**Supplementary Table 3**), the minor allele A of rs80285134 had no effect on BMI in the total study population but increased BMI in short people (beta= 0.30, P = 5.9 × 10^−4^), while it decreased BMI in tall people (beta= −0.37, P= 1.7 × 10^−6^). The distribution of BMI for rs80285134 is shown in **Supplementary Figure 10**.

Using mtCOJO we then conditioned tall people’s BMI GWAS results on short people’s BMI GWAS results (**Figure 2 B**). The same 2 SNPs in the same locus on 13q31.3 were identified.

**Figure 2B.**
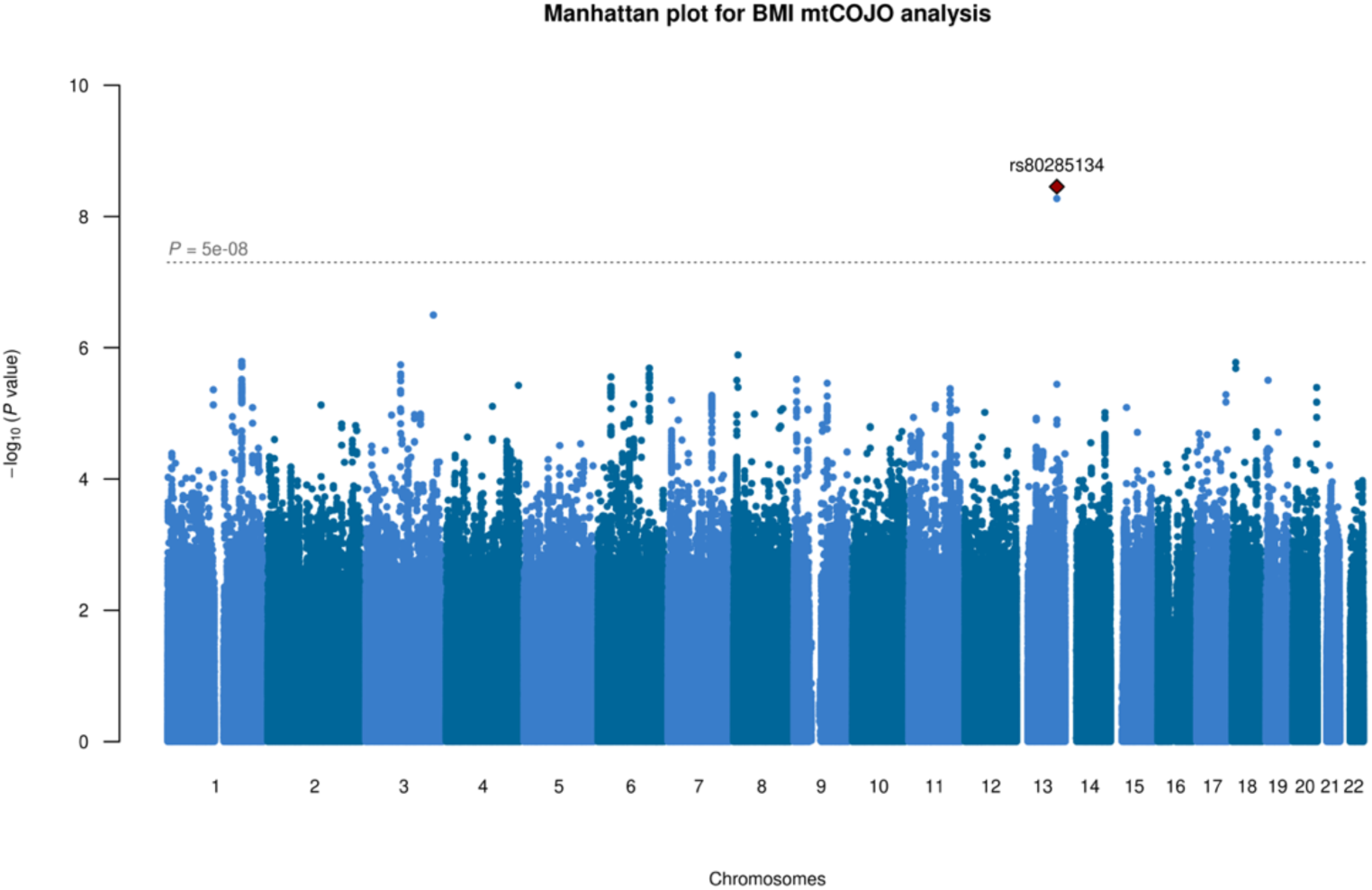
mtCOJO results of tall people’s BMI GWAS results conditioned on short people’s BMI GWAS results. The top SNPs in both analyses are located in an open chromatin region in the vicinity of *GPC6*, which is associated with bone mineral density and plays a key role in osteoporosis.

### Gene-set and cell-type enrichment analyses

We used FUMA (Functional Mapping and Annotation of Genome-Wide Association Studies) [10] to obtain gene-level summary statistics from our GWAS and mtCOJO results to perform gene-set and cell type enrichment analysis [11] for brain cells from the Linnarsson lab (http://mousebrain.org/) and bone marrow tissue (**Supplementary Figure 11)**. One gene, *SLC5A*, passed the genome-wide significance threshold in our gene-based test. Gene-set analysis was then conducted using summary statistics of GWAS BMI-total, BMI-short and BMI-tall, and our mtCOJO results. 16 gene-sets were significantly enriched for BMI in the total population. One gene set was enriched for BMI-short, reactome ntrk2 activates rac1, while one gene set was enriched for BMI -tall, neurogenesis. No gene-sets were found to be enriched for the mtCOJO results (**Supplementary Table 4**). In tissue enrichment analysis using GTEx datasets, 12 tissues were enriched for BMI short (**Supplementary Figure 12 B; see Supplementary Figure 12 A for results in all height groups**), namely: cerebellum, cerebellar hemisphere, frontal cortex BA9, cortex, anterior cingulate cortex BA24, pituitary, nucleus accumbens, hippocampus, amygdala, caudate nucleus, and putamen. For BMI-tall (**Supplementary Figure 12 C**), 4 brain regions were enriched: cerebellum, cerebellar hemisphere, cortex, and frontal cortex BA9. For the mtCOJO results, no tissues were significantly enriched (**Supplementary Figure 12 D & E**).

In our cell enrichment analysis using brain single-cell RNA data, 10 cell types were significantly enriched for BMI-short after multiple-testing correction **(Supplementary Figure 13 A)**, namely L23_Ddn cell in mouse somatosensory cortex, Vglut2_11 cell in mouse hypothalamus neurons, OPC cell in mouse oligodendrocytes, Gaba cells from both mouse and human midbrain, and HBCHO1, Neurons, CNS_neurons, and Hind cells in mouse brain atlas. Similarly, 8 cell types were significantly enriched for BMI-tall after multiple testing correction (**Supplementary Figure 13 B**), namely CA1PyrInt cell in mouse hippocampus, Vglut2 17 A930013F10RikPou2f2 cell in mouse hypothalamus neurons, NbGaba and Gaba cells in human midbrain, NbL2 cell I mouse midbrain, and HBINH3, HBGLU3 and TEGLU1 in mouse brain atlas. In contrast, there were no brain cells that were significantly enriched for mtCOJO results. In our cell enrichment analysis using bone marrow single-cell RNA data, no cells were significantly enriched for BMI-short, BMI-tall or mtCOJO results after multiple testing correction (**Supplementary Figure 14**). In summary, we provide consistent evidence that brain tissue is involved in differences in genetic architecture between short and tall individuals.

### Phenome-wide association analyses (PHEWAS)

We followed up on the previously detected locus with differential associations for BMI-short vs BMI-tall detected through mtCOJO (results section ‘Multi-trait-based conditional and joint analysis, mtCOJO’) and on loci passing both SNP criteria (results section ‘SNPs with different effects in the two height groups’). Regarding mtCOJO, we successfully extracted the associations of this locus (rs80285134) in 109 previously published GWASs and then ran PHEWAS. Three associations passed the FDR correction threshold, which were bone mineral density [12] (PMID=29304378; p=1.1 × 10^−4^, **Supplementary Figure 15**), number of days/week of vigorous physical activity of > 10 minutes (PMID = 31427789, p = 3.0E x 10^−4^) and moderate to vigorous physical activity levels (PMID = 2989952, p =3.1 × 10^−4^). Regarding the loci passing both SNP criteria, there were 1194 associations passing the genome-wide significance threshold (p<5 × 10^−8^), including metabolic traits and cognitive ability (**Supplementary Table 5)**.

We then conducted PHEWAS within the UKBB samples for 7 pre-selected phenotypes (see methods) on rs80285134 and 57 candidate height-dependent BMI variants (from **Supplementary Table 2**). We thus identified 23 short-specific BMI variants and 25 tall-specific variants as significantly associated with fat percentage (**Supplementary Table 6**). In addition, we identified 13 short-specific variants and 19 variants significantly associated with waist-hip ratio (WHR). The only significant association between height-specific variants and non-anthropometric traits was rs9947450 (a variant in *MC4R*) which had a positive effect on portion energy intake (beta= 2.74, p-value=8.99 × 10^−5^).

### Estimates of SNP-based heritability in short vs tall people

Using Genome-Complex Trait Analysis (GCTA), we estimated SNP-based heritability of BMI in short people and tall people in young (age <54 years), middle (54 <= age <= 61), and old (age > 61) groups from the discovery dataset (**Table 1**) and replication datasets (**Supplementary table 7**). The SNP-based heritability of BMI-short and BMI-tall in the young age groups were 0.27 (SE=0.018) and 0.30 (SE=0.015), with 93.50% (SE=0.043) genetic correlation. The SNP-based heritability of BMI-short and BMI-tall in the middle age group were 0.24 (SE=0.016) and 0.28 (SE=0.016), with 97.99% (SE=0.0505) genetic correlation. The SNP-based heritability of BMI-short and BMI-tall in the old age group were 0.212 (SE=0.013) and 0.24 (SE=0.019), with 100% (SE=0.0597) genetic correlation. The genetic correlation differences between BMI-short and BMI-tall thus decreased with age.

### Genetic correlations of BMI-short and BMI-tall with complex traits by LD hub

We used summary statistics from the BMI-short group and BMI-tall groups (see section ‘Genome-wide association analyses (GWAS) above)’ to estimate the genetic overlap using LD hub with 275 traits with GWAS results using a European cohort excluding the UKB cohort (e.g. anthropometric, bone, brain volume, cardiometabolic, glycemic, hematological traits, hormone, kidney, metabolites, and psychiatric disorders). There were 88 traits associated with BMI-short or BMI-tall (**Supplementary Figure 16**). BMI-tall was specifically (i.e. not in BMI-short) genetically correlated with 31 traits such as cigarettes smoked per day, free cholesterol in very large HDL, celiac disease, ulcerative colitis, and bipolar disorder. In contrast, BMI-short was specifically genetically correlated with 11 traits such as adiponectin, Parkinson’s disease, childhood IQ, anorexia nervosa, and insomnia.

### Transcriptome-Wide Association Studies (TWAS)

We used the summary statistics of our GWASs for BMI in short and tall individuals to perform TWAS across gene expression datasets in 53 tissues. We performed 256,074 association tests between SNPs and gene expression. Among 53 tissues, brain, blood and skeletal muscle tissues were enriched. We found 456 associations (**Supplementary Table 8**) for GWAS-short and 538 associations for GWAS-tall (**Supplementary Table 9**) that were significant after Bonferroni correction (P <0.05/ 256,074). Among the significant associations for BMI-short and BMI-tall, 342 associations were both significant for BMI-short and BMI-tall. There were 122 associations specific for GWAS-short and 203 associations specific for GWAS-tall. We identified a significant negative genetic correlation between *YPEL3* expression and BMI-short in 12 tissues but only one negative genetic correlation between *YPEL3* expression and BMI-tall in one tissue (brain spinal cord cervical) (**Supplementary Table 8 and Supplementary Table 9**).

Then we used our mtCOJO results for TWAS analysis. Among 256,074 cis-effects of gene expression in specific tissues with mtCOJO results (GWAS of BMI-short conditioned on GWAS of BMI-tall and vice versa), there were 9 significant associations in GWAS of BMI-short conditioned on BMI-tall (P <0.05/ 256,074; **Supplementary Table 10**) and 6 significant associations in GWAS of BMI-tall conditioned on BMI-short after Bonferroni correction (P <0.05/ 256,074; **Supplementary Table 11**).

### Height-dependent BMI variants in Mendelian Randomization (MR) analysis

To detect possible causal effects of BMI in short and BMI in tall people on diseases, we conducted MR analysis using height-specific BMI instruments[13]. There were 21 and 24 independent variants from **supplementary table 2** which were indicated as “short” and “tall” instruments and that were genome-wide significant in GWAS-short and GWAS-tall. Those were selected as short-specific and tall-specific BMI instruments, respectively. From 55 candidate outcomes we found that both short-specific and tall-specific BMI alleles increase the risks of breast cancer and obsessive-compulsive disorder (OCD). Furthermore, we found evidence for causal effects of short-specific BMI alleles on the risks of bipolar disorder, eczema, and hip osteoarthritis. We found evidence for causal effects of tall-specific BMI alleles to increase the risk of major depressive disorder and to decrease the risks of chronic kidney disease, inflammatory bowel disease and ulcerative colitis **(Supplementary Figure 17)**.

### Summary-data-based Mendelian Randomization analysis (SMR)

As the top SNP from the mtCOJO results, rs80285134, was associated with bone mineral density, we aimed to identify genes whose expression levels in osteoclasts were associated with height-specific BMI traits and height-dependent BMI traits because of pleiotropy. To further characterize functionally relevant genes and to identify potential pleiotropic effects on gene expression and BMI, we integrated the complete sets of BMI-short and BMI-tall GWAS summary results and BMI mtCOJO results with an osteoclast eQTL dataset [14] using the SMR software [15]. A total of 1050 genes with an eQTL association significant at P < 5 × 10^−8^ were identified and included in the analysis. Using a Bonferroni multiple-testing corrected significance threshold for SMR of < 4.8 × 10^−5^, we identified significant associations for 3 genes (*UHPR1BP1, HSD17B12* and *NUP160*) with BMI-short and 2 genes (*UHRF1BP1* and *AF131215*.*2*) with BMI-tall (**Supplementary table 12**).

Heterogeneity (HEIDI < 0.05) was detected for one gene (*NUP160*) for BMI-short and one gene (*AF131215*.*2*) for BMI-tall. Therefore, *HSD17B12* appears a candidate for harboring genetic effects on osteoclast gene expression in BMI-short, while *UHRF1BP1* may be relevant to both BMI-short and BMI-tall. Expression of *UHPR1BP1* was found to be negatively associated with BMI-short and BMI-tall, and *HSD17B12* was found to be positively associated with BMI-short. For *UHRF1BP1*, associated diseases include hypogonadotropic hypogonadism with anosmia and systemic lupus erythematosus. Gene Ontology (GO) annotations related to this gene included identical protein binding and histone deacetylase binding. For *HSD17B12*, the associated diseases include qualitative platelet defect and central hypoventilation syndrome. Recent studies show that deletion of *Hsd17b12* in adult mice results in lean body weight due to a decrease in food intake [16]. Among its related pathways are fatty Acyl-CoA biosynthesis and metabolism of steroid hormones.

## Discussion

To shed light on possible differences in the genetic architecture of BMI for short vs tall people, we have performed a range of analyses. We provide converging evidence for such differences. First, we identified several loci with height-specific effects on BMI (including *MC4R, SAE1, PAK5, CBLN4, ADARB1, TNRC6B*). Second, we found a significant association between a short people-specific locus in *MC4R* and energy portion size. *MC4R* encodes the melanocortin-4 receptor, which has been implicated in regulating meal size in rodent studies [17]. Third, using mtCOJO, we identified one SNP with significant BMI-effect differences between short and tall people: rs80285134. Using PHEWAS within the UKBB, the strongest associated phenotype with this SNP was bone mineral density. A leptin-independent homeostatic method of regulating body weight (“gravitostat”) was recently identified, depending on body weight sensing by bone cells in rodents [18]. The eQTL analyses identified *HSD17B12* to be associated with BMI-short. A deletion in this gene in adult mice results in a lean phenotype [16]. In addition, a translational human randomized clinical trial demonstrated that increased weight loading reduces body weight and body fat in obese subjects[19]. Fourth, we found that SNP-based heritability differences between BMI-tall and BMI-short decreased from young age to old age, with 93.5% (SE=0.043) genetic correlation at young age. Perhaps environmental factors as exposure to easily accessible palatable food, have a stronger impact on shorter people, a notion supported by our finding that short people eat more calories when corrected for body weight. Fifth, transcriptome-wide analyses (TWAS) indicated that the genes associated with different effects in tall vs short have significant cis-eQTL effects on brain tissue: the genes involved in cell proliferation and apoptosis were more enriched in BMI-short (such as *YPEL3*); in contrast, genes associated with metabolic disease, Osteogenesis Imperfecta, or birth weight were more enriched in BMI-tall (such as *PDCD2, TMEM91* and *GUF1*). Sixth, with TWAS results of mtCOJO we identified 39 genes associated with height-specific BMI in brain tissue, e.g. *CPDD2, GUF1*, and *MLH3*. And finally, BMI-short and BMI-tall were genetically correlated with different diseases.

We identified two well-known loci, 16p11.2 (nearby *FTO*) and 18q21.32 (nearby *MC4R*), to be more strongly associated with BMI in short people than tall people. The top SNP from 16p11.2 is also genome-wide significantly associated with mean corpuscular hemoglobin and whole bone mineral density. *MC4R* is a well-known gene associated with both height and BMI. Based on both criteria for BMI-short and tall-specific SNPs, one locus in *ADCY3* seems a particularly plausible candidate as it was previously identified as a height-specific genetic variant for BMI in young children [5]. In addition, the locus of *MC4R* (specific for short) is also associated with portion size of energy intake in our dataset, which is in line with MC4R activity affecting meal size [17]. This supports our hypothesis that genetic variants underlying BMI in short people are more related to the behavioral response to food than in tall people.

We have identified one locus (rs80285134) containing height-specific BMI variants using mtCOJO analysis. This variant is a chromatin regulatory variant for *GPC6* that was not only known to be associated with BMI but also it is also a relatively novel gene with a role in skeletal biology [9, 20]. The most strongly associated trait for this variant is bone mineral density, hinting at the possible contibution of height-specific BMI genes to the pathogenesis of osteoporosis. FUMA analysis of mtCOJO results prioritized one gene, *SLC5A5*, that is associated with dyshormonogenesis. mtCOJO analysis of BMI conditioned on height had been conducted previously to estimate the height-unconfounding BMI variants [8]. However, we identified BMI variants specific for height for the first time, to our knowledge.

Future work should be done to provide further biological insight into the biological determinants of BMI in short and BMI in tall, such as animal studies investigating effects of candidate height-specific BMI genes on BMI and feeding behavior. Both TWAS and tissue enrichment analyses using the GTEx dataset on our mtCOJO results showed enrichment in brain and muscle skeletal tissues, which may help conduct such translational studies. Based on our findings, future human genetics studies should take into account differences between BMI-short and BMI-tall, including per age group, when they perform cross-disorder analyses. Our MR findings of different diseases being causally linked to BMI-short vs BMI-tall may be taken into account when designing preventative intervention studies in risk groups.

In conclusion, our findings highlight the role of height in the genetic underpinnings of BMI, provide biological insight into mechanisms underlying height-dependent differences in BMI and set the stage for future cross-disorder genetic analyses as well as translational studies.

## Materials and Methods

### UK Biobank participants and sample quality control

The UK Biobank (UKB) is a large cohort of over 500,000 United Kingdom residents. It allows exploration of the genetic, environmental and lifestyle factors associated with a wide range of complex diseases [21]. The assessments cover a wide range of social, cognitive, lifestyle, and physical health measures. For the current analyses, we requested whole-genome common variant data and BMI-relevant phenotypic data, such as weight, height, impedance measurements, physical activities, and dietary habits. Informed consent was obtained from all participants, and this study was conducted under generic approval from the NHS National Research Ethics Service (approval letter dated 13 May 2016, Ref. 16/NW/0274) and under UK Biobank approval for application #44344 ‘Genome-wide association study of Body mass index in short and tall individuals’ (PI Bochao Lin).

We analyzed data from 502,538 participants of the UKBB. 378 individuals were excluded as their self-reported sex did not match their genetically determined sex. In addition, we excluded individuals with missing information for BMI, height and age (n=1,975). Individuals with outliers (defined as < or > 5 times the Standard Deviation, SD) of BMI (n = 473), height (n=5), weight (n=125) and age (n=0) were further excluded, resulting in 485,334 individuals. Furthermore, we restricted all analyses to 413,396 participants of European ancestry using genetic principle components (PCs) (more detail in **Supplementary Figure 18**). After removing related individuals (n = 53,223) based on relatedness using kinship data (estimated genetic relationship > 0.044), there were 359,400 unrelated genotyped participants left for the analyses described below.

### Phenotype-level analyses

We visualized the distributions of BMI, fat percentage, energy intake and portion energy intake in deciles of height groups (**Supplementary Figures 1-3**). Furthermore, we calculated Pearson correlations between BMI, height, fat percentage, and height percentage in height decile groups and age (**Supplementary Figure 4**).

### Genotyping, imputation and SNP quality control

In March 2018, UK Biobank released genetic data for 487,409 individuals, genotyped using the Affymetrix UK BiLEVE Axiom or the Affymetrix UK Biobank Axiom arrays (Santa Clara, CA, USA) which contain over 95% common SNP content [22]. Pre-imputation quality control, imputation and post-imputation cleaning were conducted centrally by UK Biobank (described in the UK Biobank release documentation). All genomic positions are in reference to hg19/build 37. Post-imputation, we selected SNPs based on minor allele frequency (MAF, > 0.01), genotyping call rate (> 0.95), Hardy–Weinberg disequilibrium test p-value > 10^−6^, and imputation quality (info > 0.8), resulting in 7,904,644 SNPs for the current analyses.

### Genome-wide analyses

#### Datasets

Before genome-wide analyses, we randomly selected 2/3 of participants (n=237,802) for our discovery dataset, and 1/3 of samples (n=121,598) for our replication dataset. Furthermore, we defined short individuals as those at the < 33% quantile of height (173 cm for men, 160 cm for women; see **Supplementary Figure 4**, n = 71,144 in the discovery dataset, n=36,761 in the replication dataset) and samples at the > 67% quantile of height (179cm for men, 165 cm for women; see **Supplementary Figure 4 and 5**) as the tall group (n =75,199 in the discovery dataset, n=38,089 in the replication dataset).

#### Genetic association testing

We ran a GWAS of BMI on 7,904,644 HapMap3 SNPs (MAF=0.01, imputation quality > 0.8) in UKB samples using PLINK2 [7] in linear regression models in both short (< 33% height quantile) and tall individuals (> 67% height quantile). Genetic-ancestry principal components (PCs) were constructed by EIGENSTRAT [23]. To improve computational efficiency, we firstly conducted mixed linear models of BMI on sex, age, square of age, recruitment center, genotyping chips and 10 PCs calculated from genotyped SNPs for UKB QCed samples, and then the residuals of that model were used as input data for GWASs (see **Supplementary Table 1**). We first conducted GWASs in the abovementioned discovery and replication datasets separately, then the summary statistics of GWASs from discovery dataset and replication datasets were meta-analyzed using the inverse variance-weighted fixed effect model implemented in METAL [24]. GWAS results reported are those of the meta-analysis, unless otherwise reported.

### Multi-trait-based conditional and joint analysis (mtCOJO)

To identify SNPs with different effects on BMI between short people and tall people, we performed BMI-short conditioning on BMI-tall using summary-level data by mtCOJO [25]. This method allows for the separation of marginal effects from conditional effects (BMI effects not confounded by height). In addition, we performed BMI-tall conditioning on BMI-short by mtCOJO analysis as a sensitivity analysis to test whether the same loci would be obtained.

### Functional Mapping and Annotation of Genome-Wide Association Studies (FUMA)

We used FUMA [10] to obtain gene-level summary statistics, gene-set analysis and cell type enrichment for bone marrow tissue.

### PHEWAS

To test the association of height-specific BMI SNPs with other traits, we queried and visualized the PHEWAS results for our top SNP (rs80285134) in the mtCOJO results of the candidate SNPs ATLAS resource (https://atlas.ctglab.nl)[26].

In addition, the variants resulting from our GWASs in short and tall (**Supplementary Table 3**) were used to conduct PHEWAS analysis in UKB samples. We had pre-selected 7 phenotypes for which we requested access to data, namely waist hip ratio (WHR), energy intake yesterday, energy portion intake yesterday (energy/ weight), trunk fat percentage, and time spent on vigorous sport, time spent on moderate sport, and time spent on light sport. The linear regression models were conducted of abovementioned phenotypes on selected SNPs with the covariates sex, age, age squared, 10 PCs, measurement center, and genetic chips. We used Bonferroni correction for multiple testing to determine the significance threshold.

### GCTA to estimate SNP heritability

The proportion of variance of BMI that can be explained by the measured and imputed SNPs (SNP-based heritability) was estimated in GCTA (Genome-wide Complex Trait Analysis) using the Restricted maximum likelihood (REML) analysis procedure, where we report the proportion of genetic variance explained on the variance of phenotype of interest ^[27]^. We selected SNPs with a minimum imputation R^2^ quality metric of 0.80 and MAF > 0.01 to calculate genetic relationship matrix (GRM). We further spilt the UK biobank short and tall individuals into young age group (age < 54, first age tertile), middle age group (54 <= age <= 61, middle age tertile), and old group (age > 61, last age tertile). Therefore, the SNP heritability of BMI was estimated in 6 different groups, namely: the short young-age group, short middle-age group, short old-age group, tall young-age group, tall middle-age group, and tall old-age group. In the end, we replicated GCTA analyses using our previously defined replication dataset.

### Genetic correlations of BMI-short and BMI-tall with complex traits by LD hub

We used summary statistics from the BMI-short group and BMI-tall group to check the genetic overlap using LD hub [28] with 275 traits with GWAS results using a European cohort excluding the UKB cohort (e.g. anthropometric, bone, brain volume, cardiometabolic, glycemic, hematological traits, hormone, kidney, metabolites, and psychiatric disorders).

### Transcriptome-Wide Association Studies (TWAS)

To investigate whether height-specific BMI-associated genetic variants influence BMI by modulating gene expression, we performed TWAS analysis that integrates both gene expression measurements and summary association statistics from GWAS to identify genes whose cis-regulated expression is associated with complex traits. To that end, we used the FUSION software [29] along with its prepackaged weights for gene expression data (see supplemental methods).

### Mendelian Randomization (MR) and Summary-based MR (SMR) analyses

To detect possible causal effects of short-specific BMI and tall-specific-BMI loci on diseases, we conducted MR analysis using height-specific BMI instruments in the TwoSampleMR R package [13]. More specifically, for short-specific BMI MR analysis, from 57 height-dependent BMI variants (**in supplementary table 2**), we selected 24 independent variants that were both significant in GWAS BMI-short and had larger effect sizes in GWAS of BMI-short than in GWAS of BMI-tall. Similarly, for tall-specific BMI, we selected 28 independent variants that were both significantly associated in GWAS of BMI-tall and had larger effect sizes in GWAS of BMI-tall than in GWAS of BMI-short. The beta values were extracted from GWAS of BMI-short and BMI-tall for short-specific BMI instruments and tall-specific BMI instruments, respectively. We performed MR analysis for 55 diseases using published summary statistics that overlapped with UKbiobank outcomes, ranging from psychiatric, autoimmune, bone, cancers, cardiovascular diseases, to metabolic diseases. More details regarding the settings were described in our previous MR study [30]. Summary-data-based Mendelian Randomization analysis (SMR) methods are reported in supplemental methods.

## Data availability

The datasets and scripts for analyses are available from the corresponding author on reasonable request.

