## Supplementary Figure for "Genome-wide analyses point to differences in genetic architecture of BMI between tall and short people"

**Lin et al.**

### Index

**Supplementary Table 1.** Distribution of BMI in the UK Biobank.

**Supplementary Table 2**. Independent SNPs list using 5 criteria extract the genetic variants with different BMI effects in short group and tall group.

**Supplementary Table 3**. The association test of significant mtCOJO result rs80285134 in BMI GWASs.

**Supplementary Table 4.** Significant gene-set results using for BMI-total, BMI-short, BMI-tall and mtCOJO results.

**Supplementary Table 5.** Significant PHWAS results using atlas GWASs for the height specific genetic variants.

**Supplementary Table 6.** Significant PHWAS results using UKBB samples for the height specific genetic variants.

**Supplementary Table 7.** SNP heritability and genetic correlation between BMI-short with BMI-tall in different age groups (replicated dataset).

**Supplementary Table 8.** Significant TWAS results of gene expression level across 53 tissues for BMI-tall.

**Supplementary Table 9.** Significant TWAS results of gene expression level across 53 tissues for BMI-short.

**Supplementary Table 10.** Significant TWAS results of gene expression level across 53 tissues for mtCOJO results (BMI-short conditional on BMI-tall).

**Supplementary Table 11.** Significant TWAS results of gene expression level across 53 tissues for mtCOJO results (BMI-tall conditional on BMI-short)

**Supplementary Table 12.** Significant SMR results of gene expression level in osteoclasts for BMI-related traits.

**Supplementary Figure 1.** BMI distribution in height decile groups.

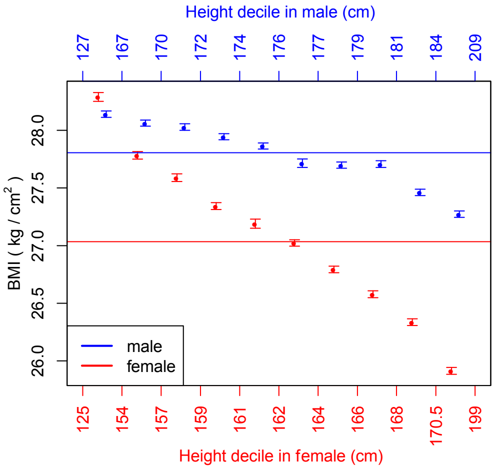


Note: The red and blue lines represent the mean BMI for females and males respectively. The dots and error bars represent the means and standard errors of BMI in height decile groups. The mean BMI decreases with increasing height in both men and women.

**Supplementary Figure 2.** Fat percentage distribution in height decile groups.

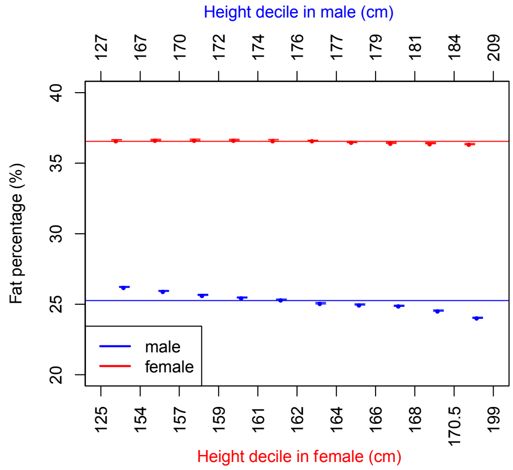


Note: The red and blue lines represent the mean fat percentage for females and males, respectively. The dots and error bars represent the means and standard errors of fat percentage in height decile groups. The fat percentages were stable in different height groups in females. However, fat percentage decreased with increasing height in males.

**Supplementary Figure 3.** Energy intake (A) and portion energy intake distribution (B) in fat percentage decile groups.

A. B.

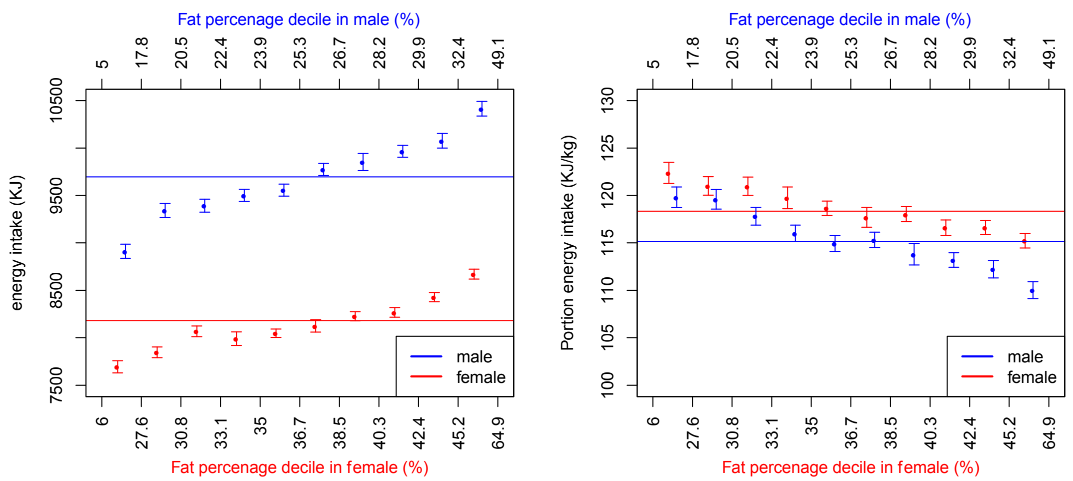


Note: The red and blue lines are the mean of energy intake (KJ) or portion energy intake - energy intake divided by individuals' weight (KJ/Kg) - for female and male. The dots and error bars show the means and standard errors of (portion) energy intake in fat percentage decile groups. The means of energy intake increase with increasing fat percentage in both men and women. However, the portion energy intake decreases with increasing fat percentage.

**Supplementary Figure 4. Phenotypic Correlations.**

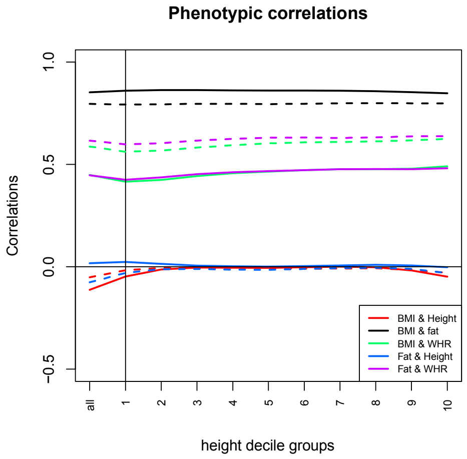


The figure shows the correlations of BMI, height, fat percentage, and waist hip ratio (WHR) in 10 decile groups for both females (dashed smooth lines) and males (smooth lines). As expected, BMI and fat percentage correlated in both males and females.

**Supplementary Figure 5. Scatt plot of distribution of BMI across height.**

**Fig 5 A** for male. **Fig 5 B** for female.

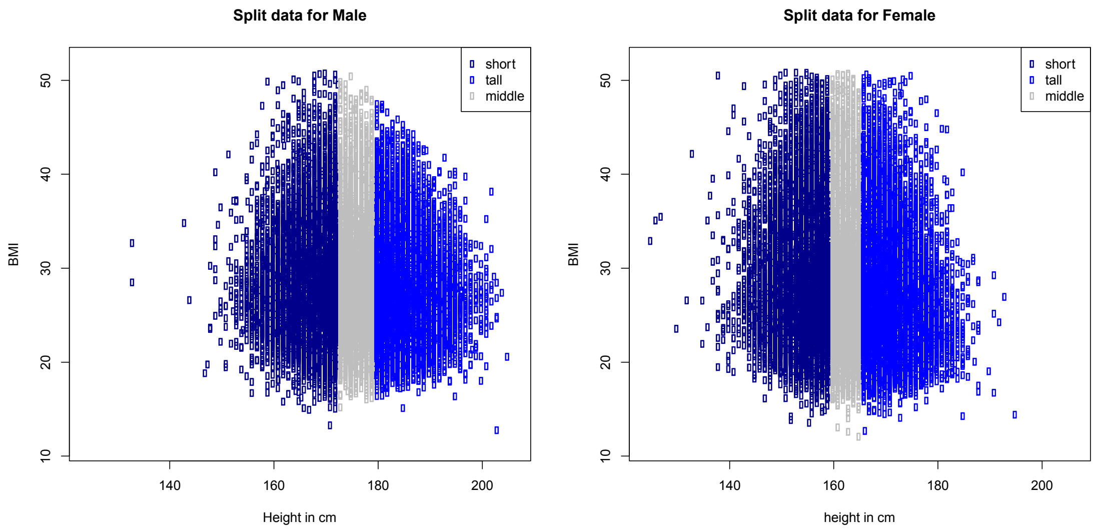

**Supplementary Figure 6. Distribution of height in men and women.**

**Fig 6 A** for males. **Fig 6 B** for females.

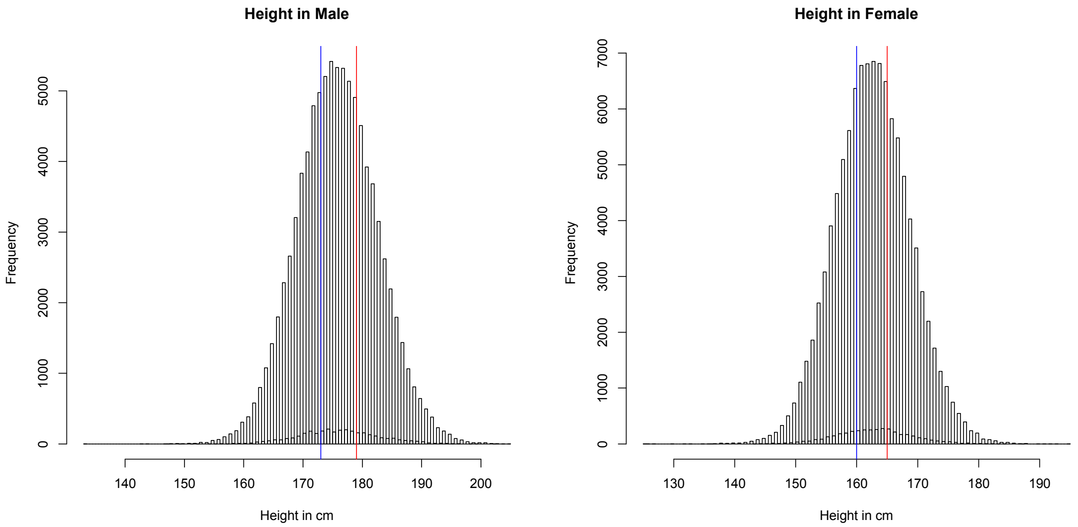


Note: for males, 33% quantile= 173cm (blue line), and 67% quantile=179 cm (red line).

For females, 33% quantile= 160cm (blue line), and 67% quantile=165 cm (red line).

**Supplementary Figure 7.** Manhattan plot of BMI in total population with the highlighted association (Meta-analysis results).

$GWAS1: Y \sim \alpha + \beta_1 * SNP + \beta_2 * sex + \beta_3 *\text{age} + \beta_4 * PC1 + \cdots + \beta_{14} * PC10 + \beta_{15} * genetic\ chips + \beta_{16} *\text{measurement center}$

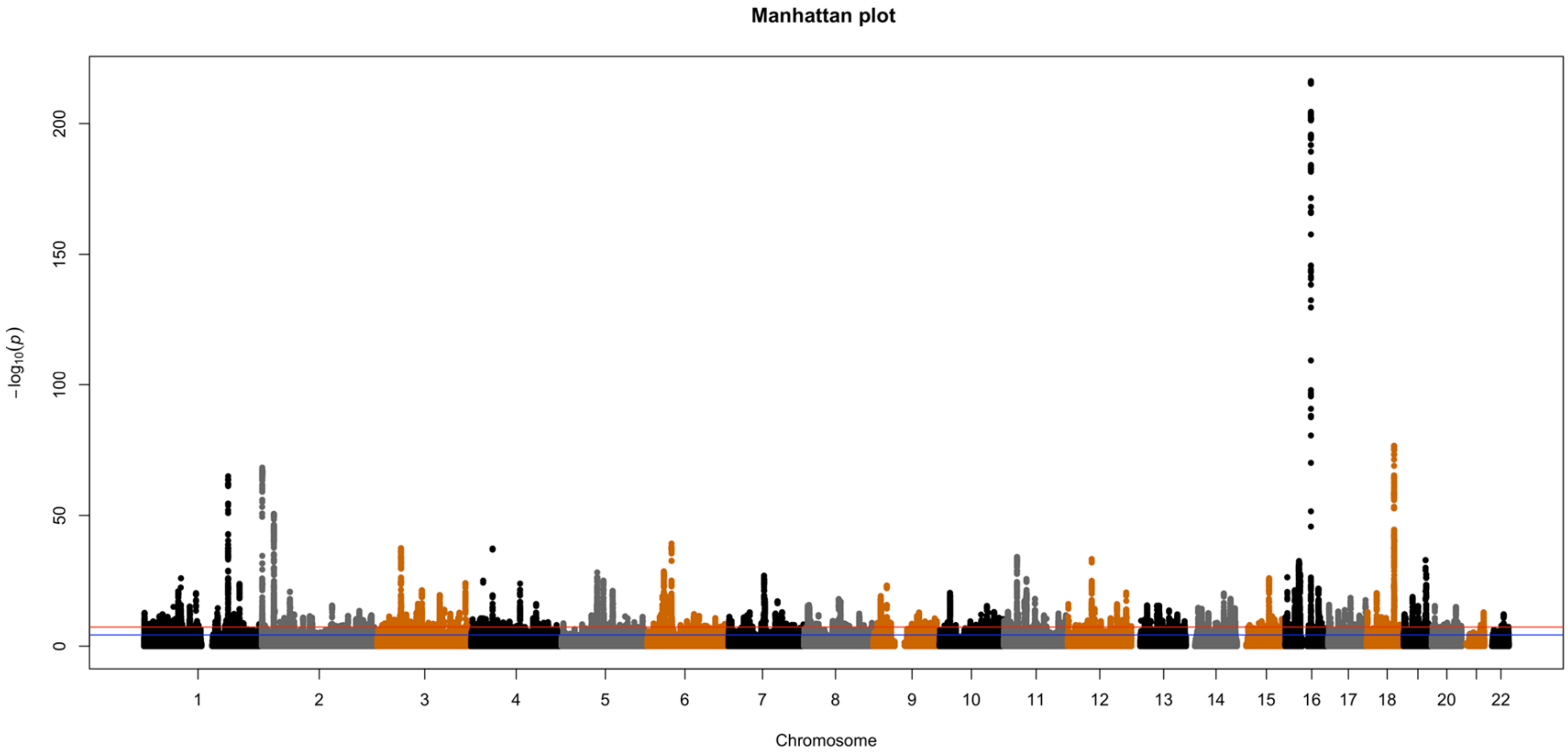

**Supplementary Figure 8 A.** Miami plot for GWAS short and GWAS tall (meta-analysis of discovery and replication cohorts), showing height-specific BMI-associated SNPs (SNPs with P-short < $5X10^{-8}$and P-tall > $5X10^{-8}$, or SNP with P -tall < $5X10^{-8}$and P - short > $5X10^{-8}$) with significant P values differences: absolute values of log10 (P-short)-log10(P-tall) > 5. The SNPs colored in blue are significant BMI-associated SNPs specific to short people. In red are significant BMI-associated SNPs specific to tall people.

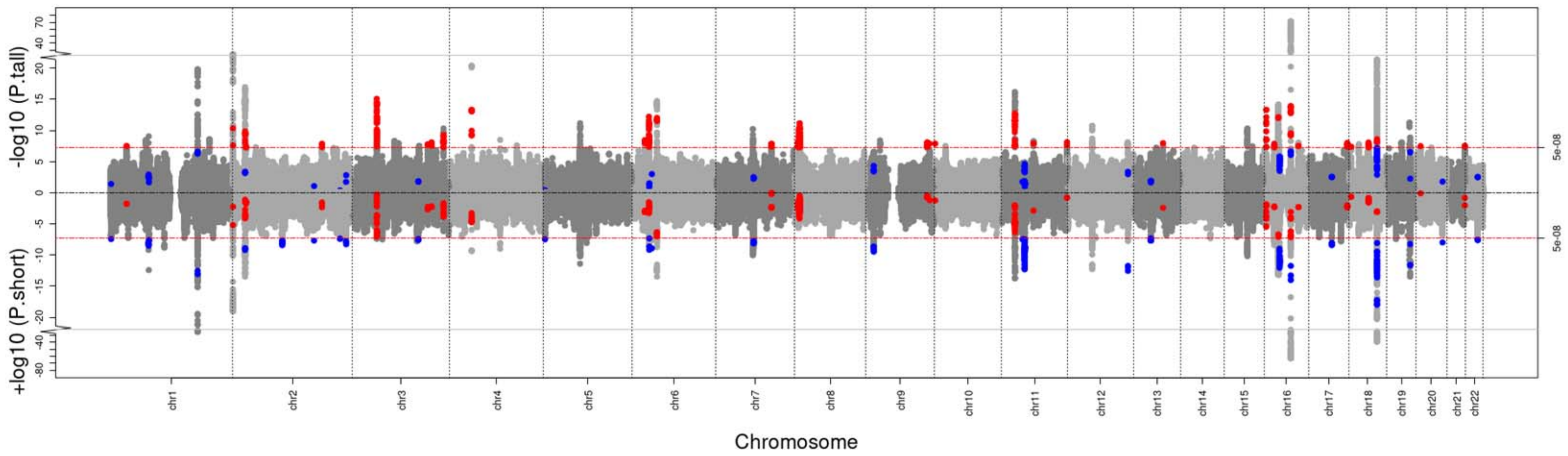

**Supplementary Figure 8 B.** Miami plot for GWAS short and GWAS tall, showing SNPs with Z scores (effect size/ standard error) differences, defined as the SNPs with absolute values of log2(Z - short) – log2(Z- tall) or vice versa > 1.

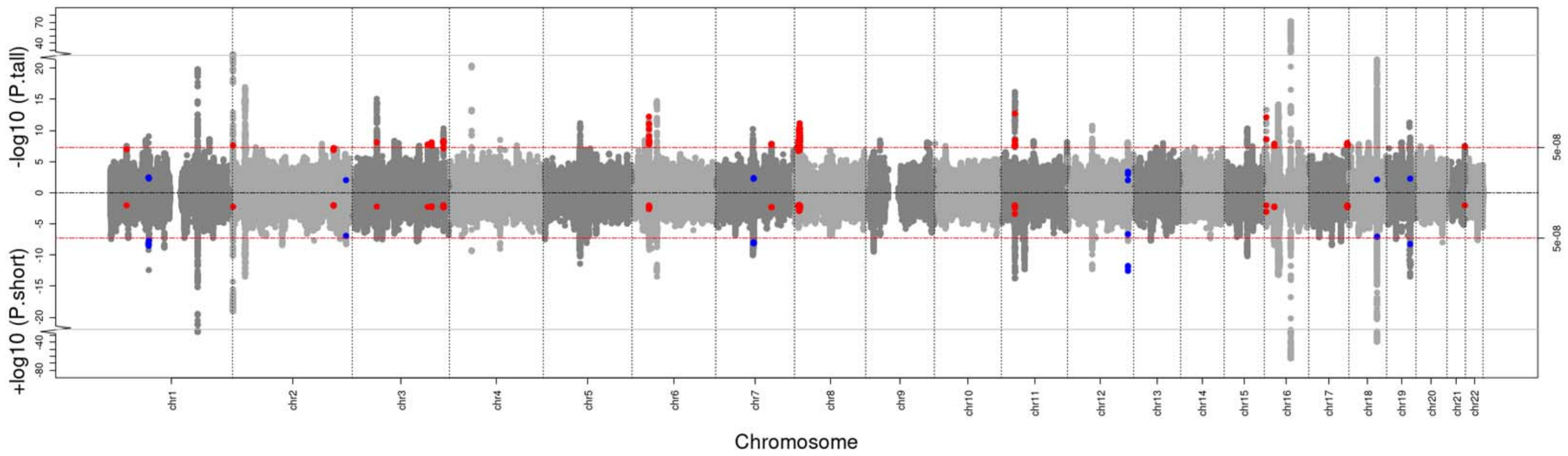

**Supplementary Figure 8 C.** Miami plot for GWAS short and GWAS tall, showing the SNPs meeting both criteria (as listed in Suppl fig 8A and 8B).

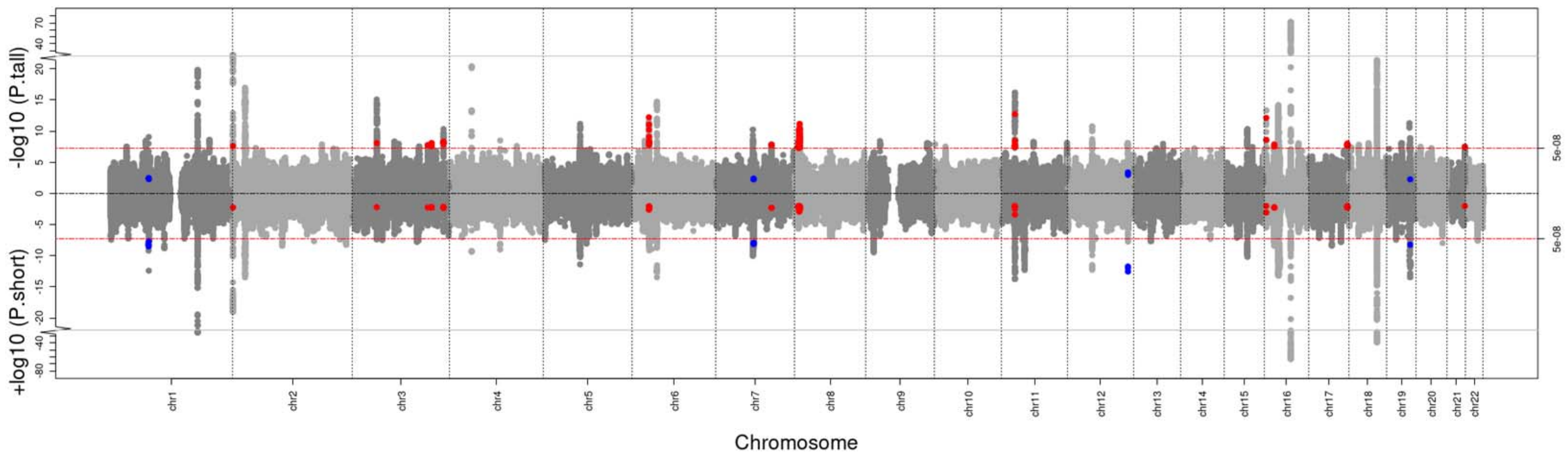

**Supplementary Figure 9.** Locus room plots for rs80285134 and rs117075592 in short people and tall people GWAS.

Figure 9 A. rs80285134 in short people.

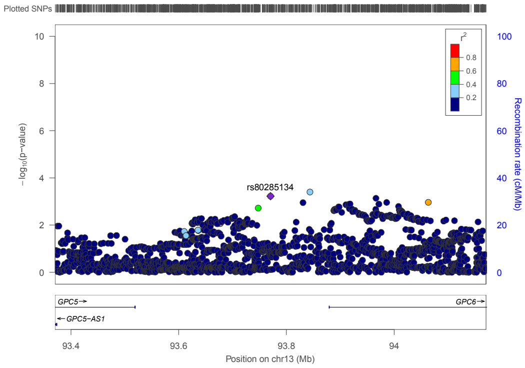


Figure 9 B. rs80285134 in tall people.

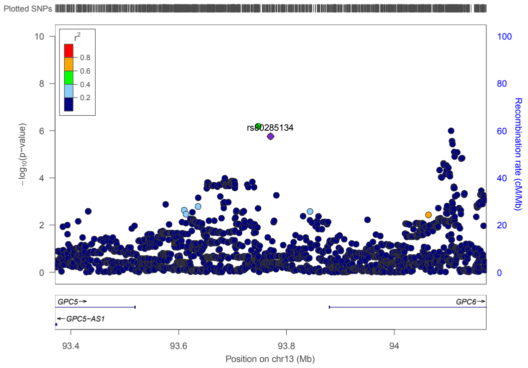


Figure 9 C. rs117075592 in short people.

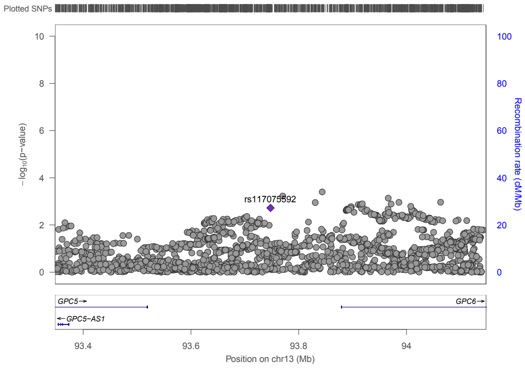


Figure 9 D. rs117075592 in tall people.

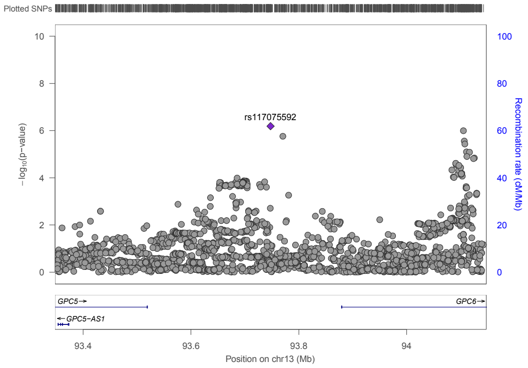


**Supplementary Figure 10.** BMI distribution in rs80285134 genotype group.

**Figure 10 A.** BMI distribution in total group.

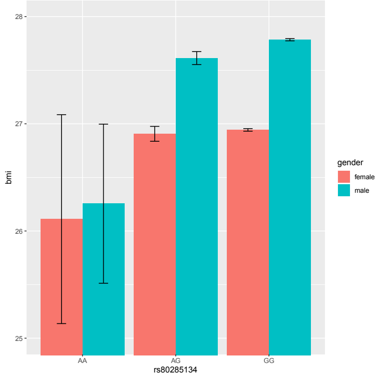

**Figure 10 B.** BMI distribution in 3 height groups (left panel for female, and right for male).

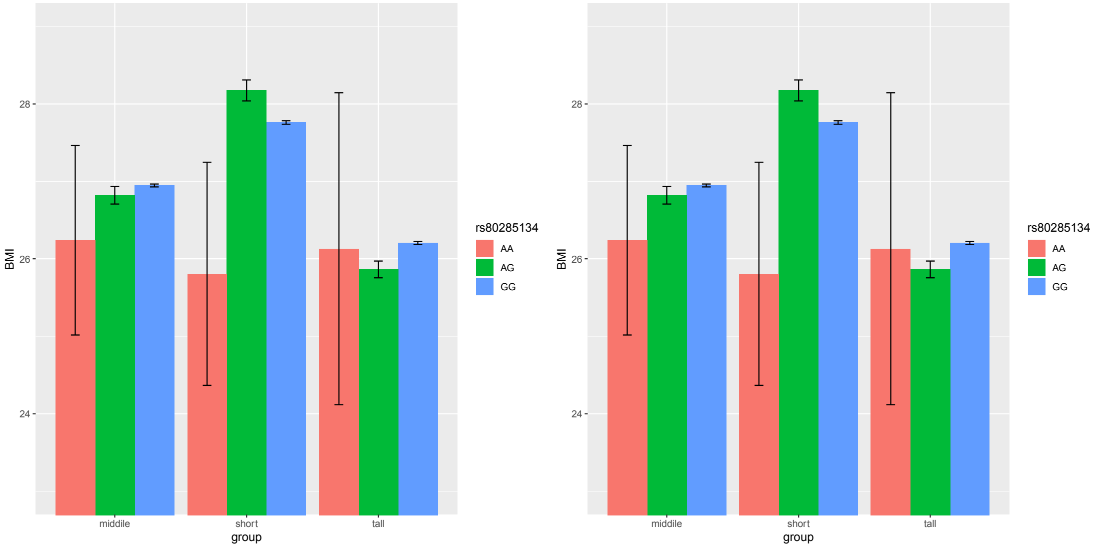


The minor allele A has no effect on BMI in total population (Suppl. Figure 10 A), but is associated with increased BMI in short people and decreased BMI in tall people (Suppl. Figure 10 B and C).

**Supplementary Figure 11 A.** Gene-level Manhattan plot for the mtCOJO results of BMI-short conditioned on BMI-tall. Genome wide significance (red dashed line in the plot) was defined at P = 0.05/18595 = 2.689e-6.

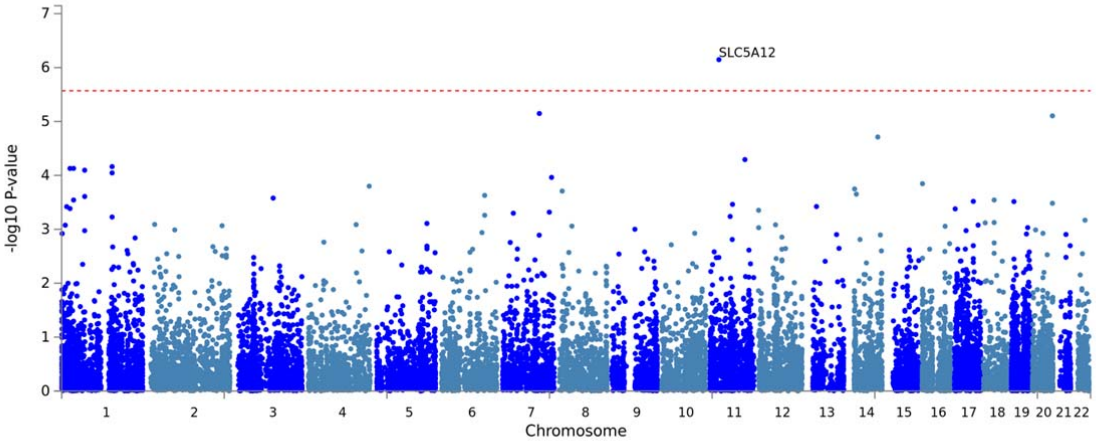


**Supplementary Figure 11 B.** Gene-level Manhattan plot for the mtCOJO results of BMI-tall conditional on BMI-short. Genome wide significance (red dashed line in the plot) was defined at P = 0.05/18734 = 2.669e-6.

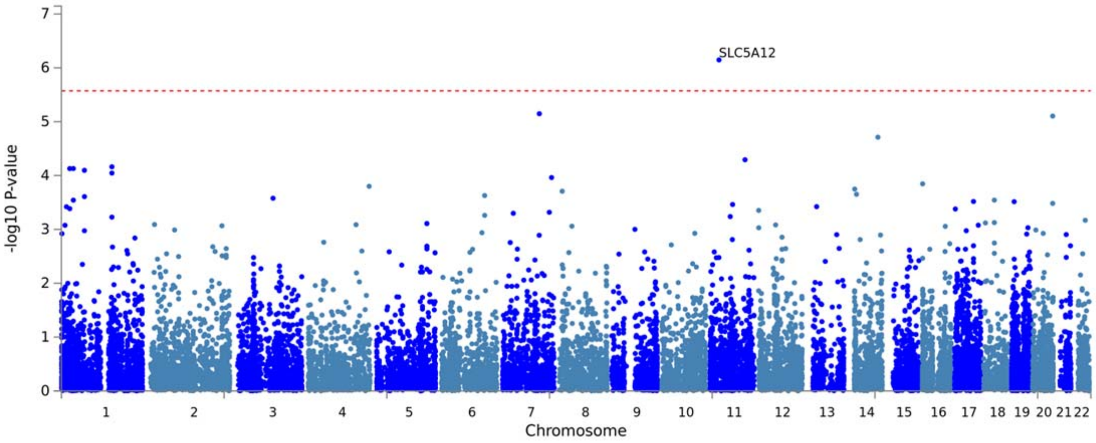

**Supplementary Figure 12. Tissue enrichment analysis using FUMA.**

**Supplementary Figure 12 A.** Tissue enrichment analysis using BMI GWAS in all height groups summary statistics.

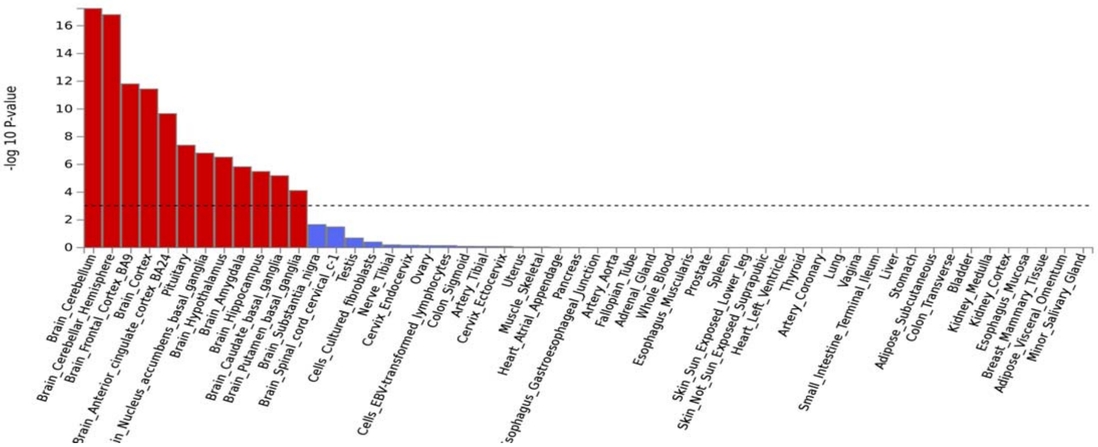


**Supplementary Figure 12 B.** Tissue enrichment analysis using BMI-short GWAS summary statistics.

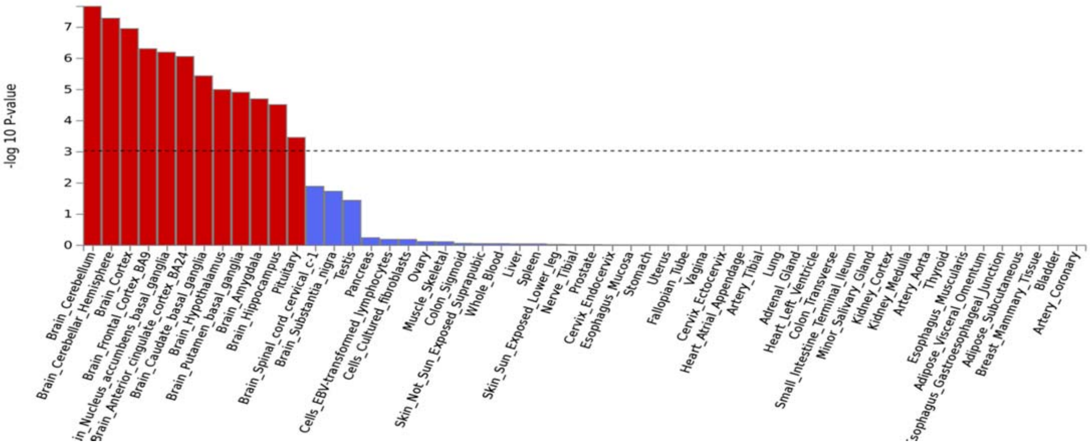


**Supplementary Figure 12 C.** Tissue enrichment analysis using BMI-tall GWAS summary statistics.

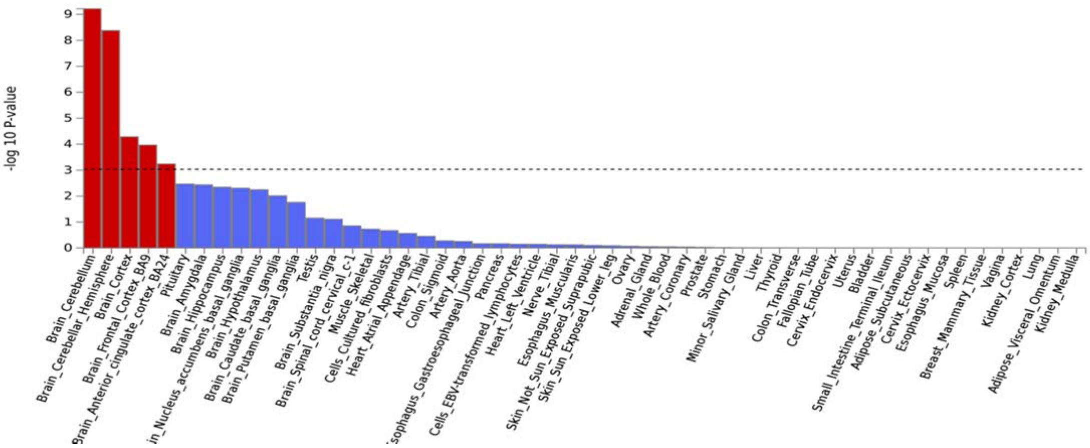

**Supplementary Figure 12 D.** Tissue enrichment analysis using mtCOJO results (BMI-short conditioned on BMI-tall).

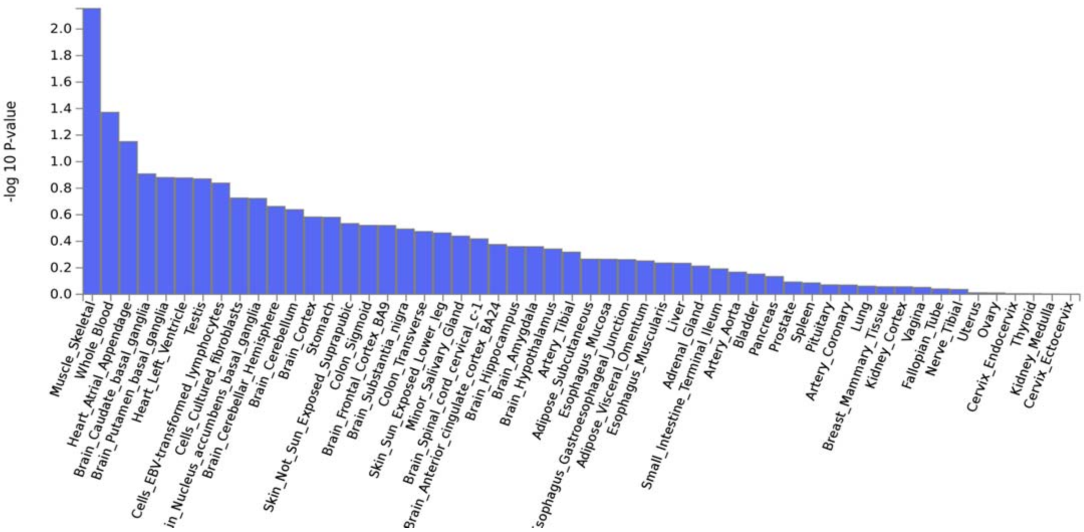


**Supplementary Figure 12 E.** Tissue enrichment analysis using mtCOJO results (BMI-tall conditioned on BMI-short).

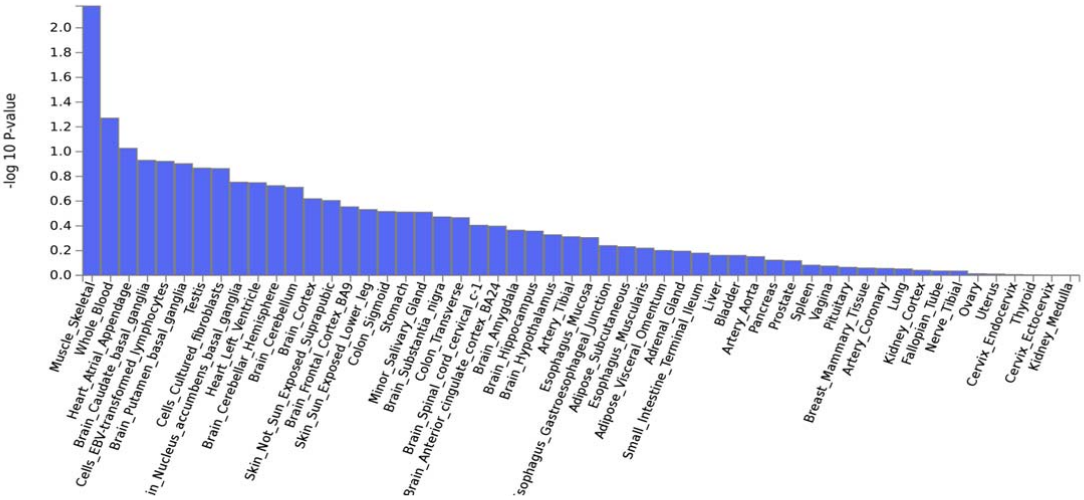

**Supplementary Figure 13. Brain cell type enrichment analysis for GWASs and mtCOJO results.**

**Supplementary Figure 13 A.** Significant brain cell type enriched for BMI-short.

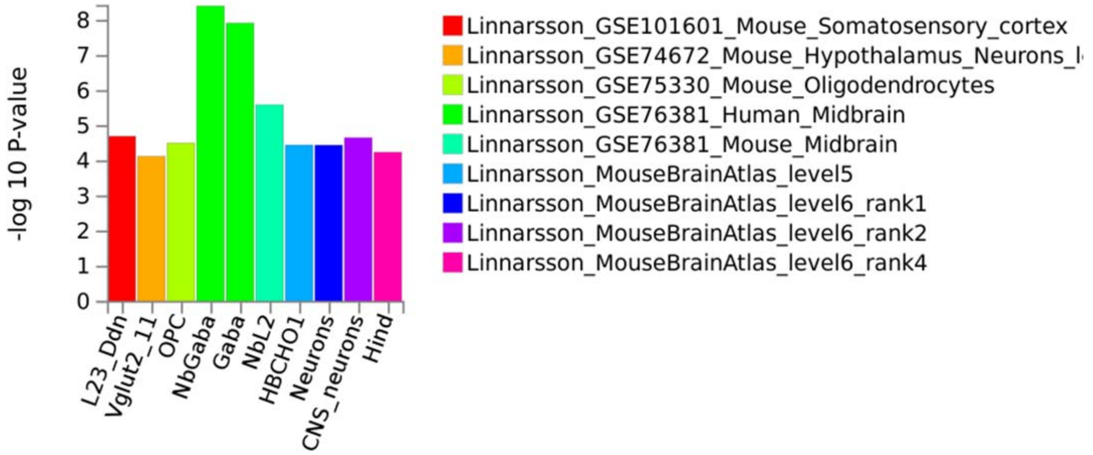


**Supplementary Figure 13 B.** Significant brain cell type enriched for BMI-tall.

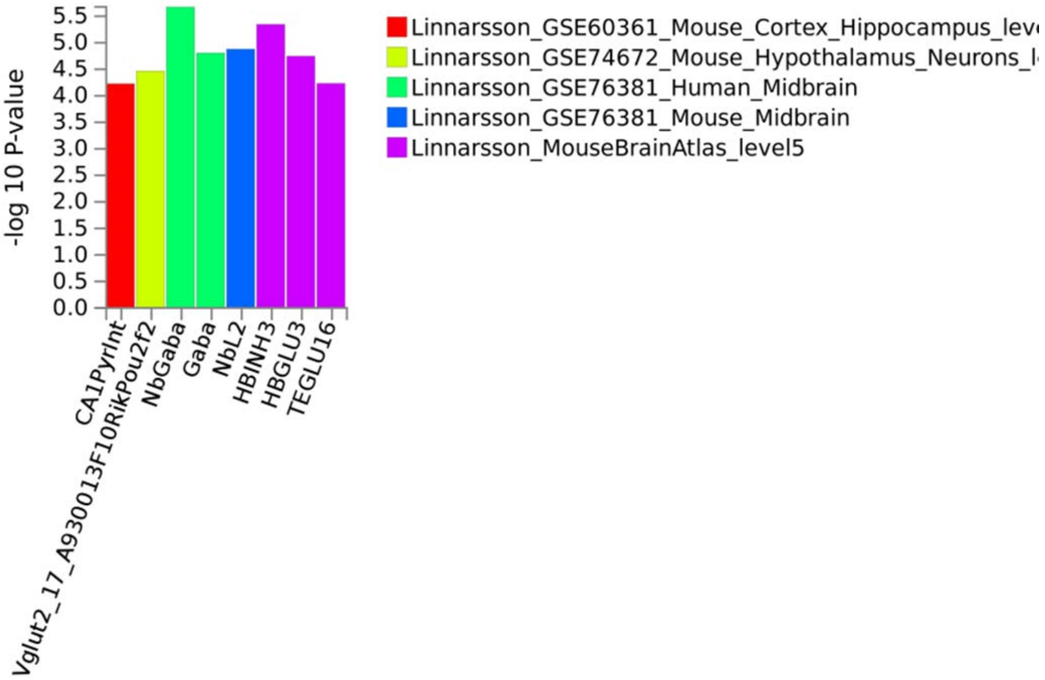

**Supplementary Figure 14. Bone marrow cell type enrichment analysis for GWASs and mtCOJO results.** The top enriched cell passed nominal p-value threshold, but did not pass Bonferroni correction.

**Supplementary Figure 14 A.** BMI-short

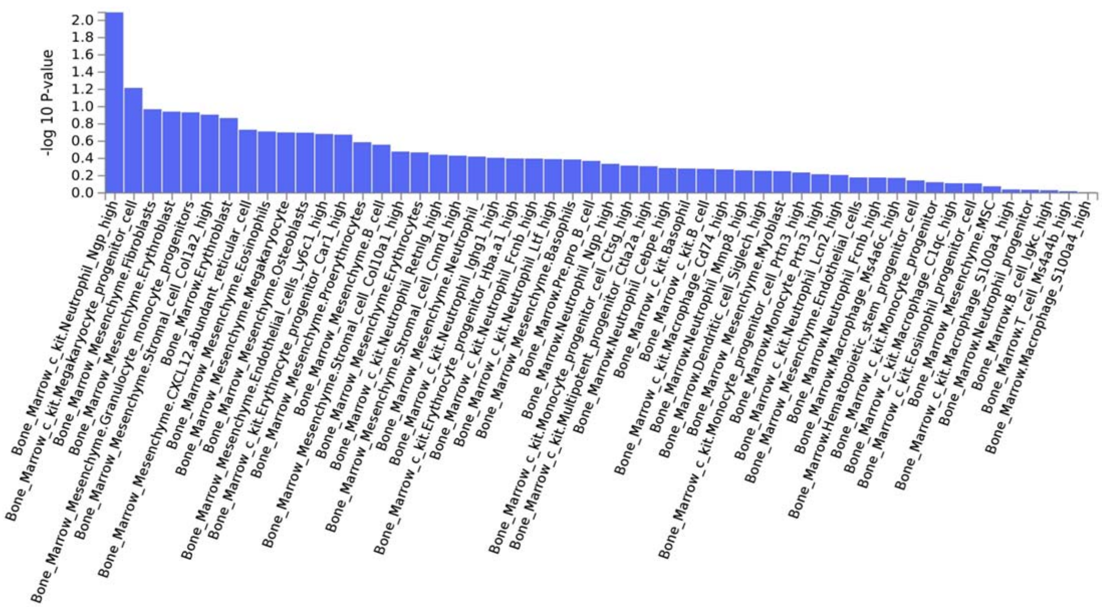


For BMI -short, the top cell was Neutrophil Ngp high cell (beta= 0.283, p=0.008).

**Supplementary Figure 14 B.** BMI-tall.

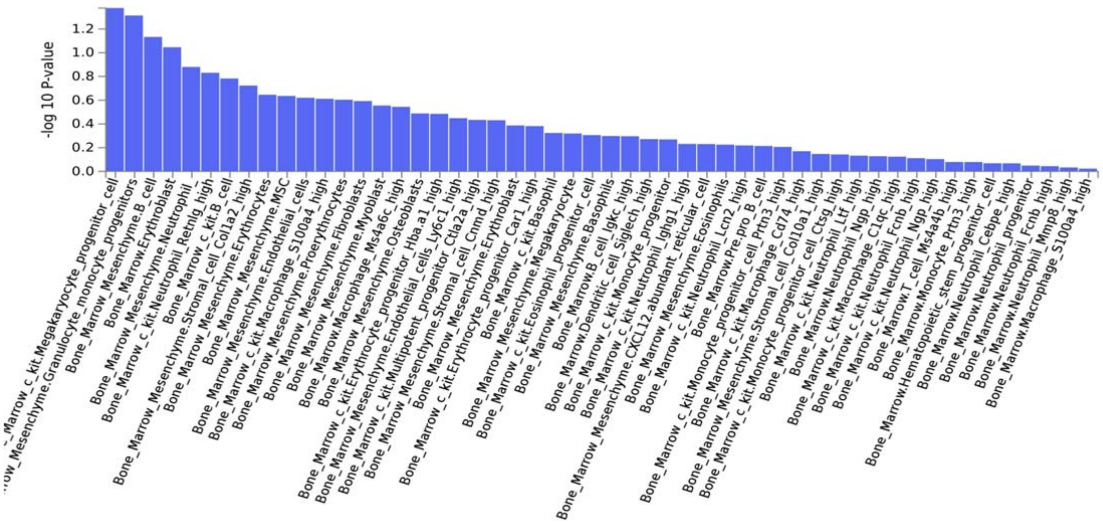


For BMI-tall, the top cell was megakaryocyte progenitor cell (beta=0.205, p= 0.042).

**Supplementary Figure 14 C.** BMI-short conditioned on BMI-tall.

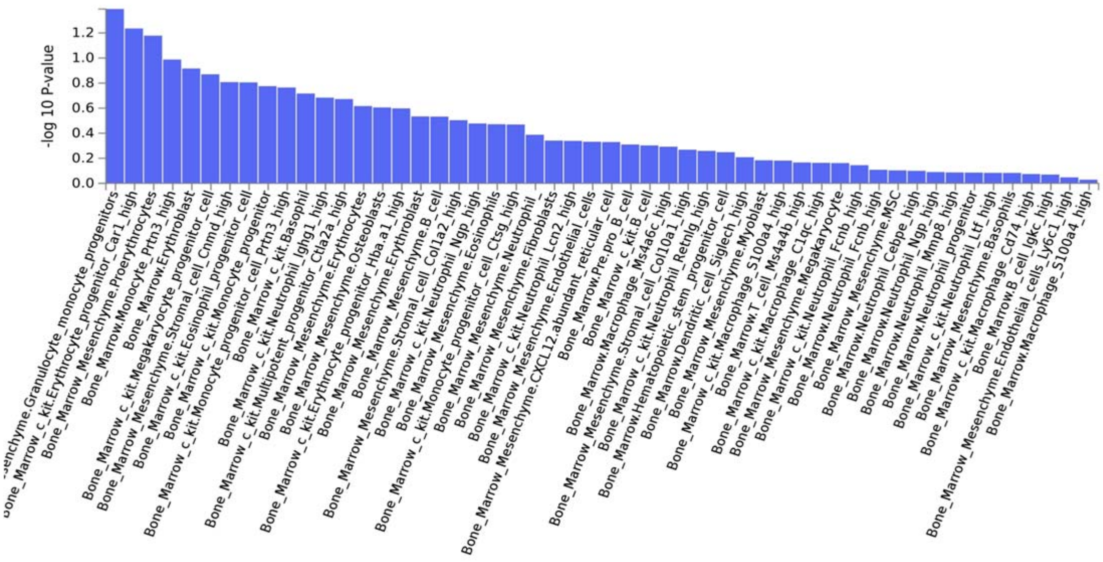


Using the mtCOJO results of  BMI- short conditioned on BMI- tall, the top cell is Bone marrow mesenchyme granulocyte monocyte progenitors were enriched (beta= 0.131, p = 0.036).

**Supplementary Figure 14 D.** BMI-tall conditional on BMI-short.

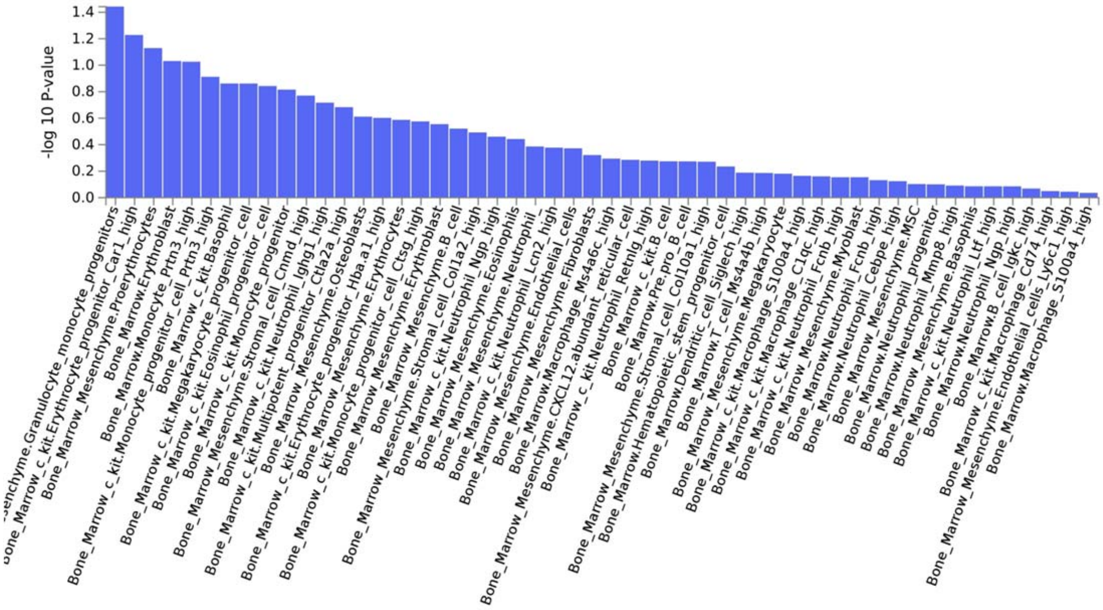


Using the mtCOJO results of BMI- tall condition on BMI- short, the top cell is Bone marrow mesenchyme granulocyte monocyte progenitors were enriched (beta=0.128, p =0.041).

**Supplementary Figure 15**. PHEWAS results for rs80285134.

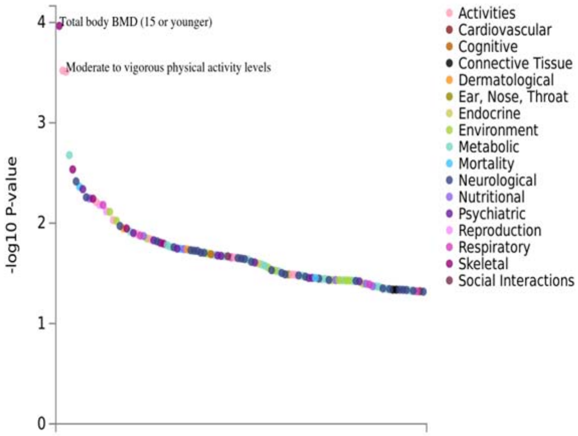


**Supplementary Figure 16**. Genetic correlations between BMI-short and BMI-tall with 88 traits using LD hub.

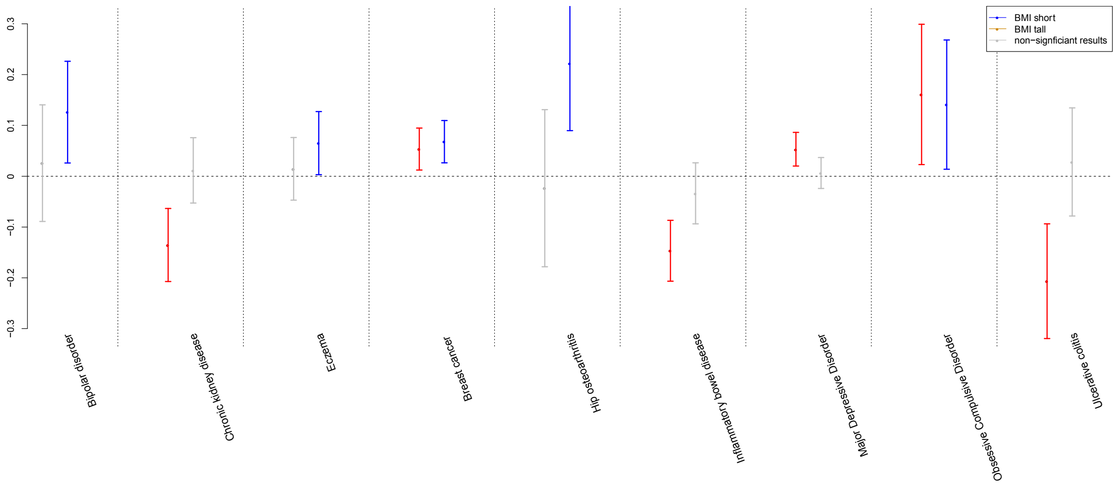


Note: The figures showed genetic correlations (rg with standard errors) of BMI-short (in blue) and BMI-tall (in red) with 88 traits. The traits were sorted by the differences we found in rg-short vs. rg-tall.

**Supplementary Figure 17**. The significant MR results (p<0.05) using short specific BMI and tall specific BMI variants as instruments in MR analysis (fix effect inverse variance weighted model).

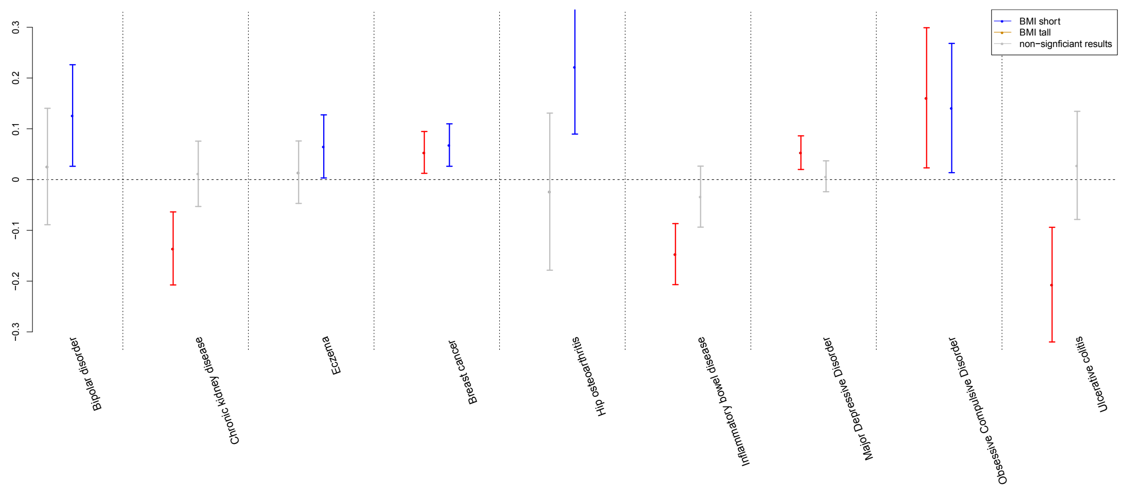


The dots are beta values from fixed-effect inverse variance weighted model. The error bars represent 95% CI. The blue color represents significant MR results of short-specific BMI, the red color represents significant MR results of tall-specific BMI, and the grey color represents non-significant MR results.

**Principal components analyses (PCA)**

The principal components (PCs) analyses within UKbiobank samples, and UKbiobank samples along with Hapmap3[1] populations were conducted by EIGENSTRAT [2]. A strict selection for SNPs with overlap with Hapmap3 SNPs was conducted: 1. MAF>0.05, HWE>0.001; 2. Removal of 20 long LD regions [3]; 3. LD pruned with an $R^2$ of 0.5; which resulted in 112,732 best quality genotyped SNPs used to calculate genetic PCs.
PCs were firstly calculated with hapmap3 population to exclude European ethnic outliers: exceeding 10 times the standard deviation of Utah residents with Northern and Western European ancestry from the CEPH collection (CEU) and Toscani in Italia (TSI) populations for the first 4 PCs. In total, 44,370 individuals were excluded from this dataset. Secondly, another PCA was conducted using the same SNPs but calculated only in UKBB samples. Another 26968 individuals were considered as ethnic outliers by the first 4 PCs exceeding 3 times the standard deviation in the UKBB samples. In the end, 413,996 individuals remained (**Supplementary Figure 1)**. Compared with the self-report ethic information, 94.8% of the ethnic-QCed samples are self-reported as “white British descent”. PCA was conducted using the same SNPs within UKBB QC-ed samples, that can be used as covariates in analyses to correct for population stratification.

**Supplementary Figure 18.** First 2PCs of UKBB quality controlled samples with Hapmap 3 dataset.

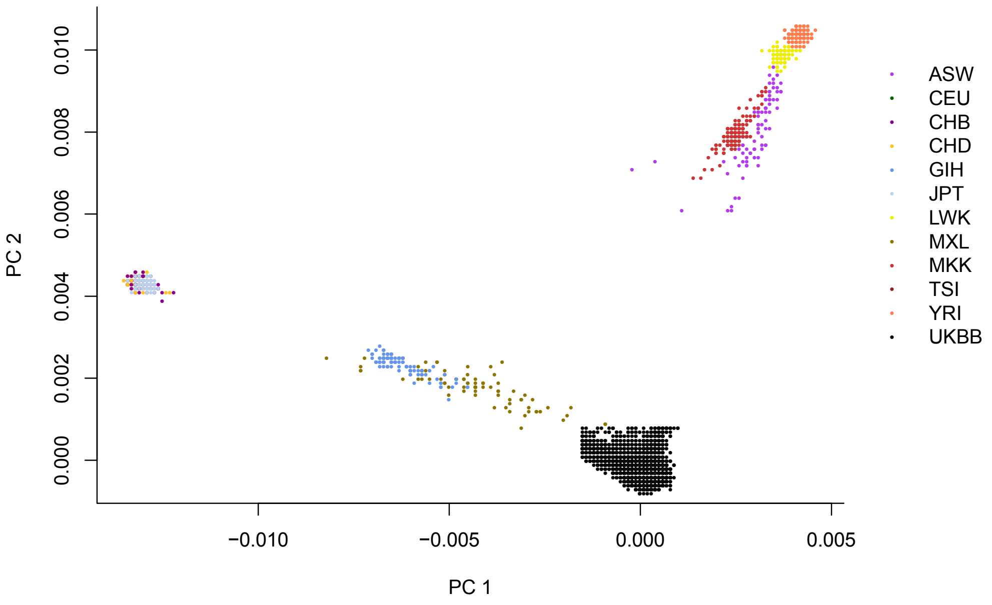

**Transcriptome-Wide Association Studies (TWAS)**

We performed summary-based TWAS approach using the FUSION software. Weights for gene expression measured using RNA sequencing data were obtained from the CommonMind Consortium30 (dorsolateral prefrontal cortex, n = 452), the Genotype-Tissue Expression Project [4] (GTEx; 48 tissues; n = 449), and the Metabolic Syndrome in Men study [5,6] (adipose, n = 563). Expression microarray data were obtained from the Netherlands Twins Registry [7] (NTR; blood, n = 1247), and the Young Finns Study [8,9] (YFS; blood, n = 1264). Briefly, this approach uses reference linkage disequilibrium (LD) and reference gene expression panels with GWAS summary statistics to estimate the association between cis-genetic components of gene expression, or alternative splicing events. First, for each panel, FUSION estimated the heritability of steady-state gene and alternative splicing expression levels explained by SNPs local to each gene (i.e., 1 Mb flanking window) using a mixed-linear model. Genes with nominally significant (P < 0.05) estimates of SNP-heritability (cis-$h2g$), are then put forward for training predictive models. Genes with non-significant estimates of heritability are pruned, as they are unlikely to be accurately predicted. Next, FUSION fits predictive linear models (e.g., Elastic Net, LASSO, GBLUP37, and BSLMM38) for every gene using local SNPs. The model with the best cross-validation prediction accuracy (significant out-of-sample R2; nominal P < 0.05) was used for prediction into our GWAS cohort. This was repeated for all expression datasets. We refrain from reporting genes from the HLA region due to complicated LD patterns.

**Summary-data-based Mendelian Randomisation analysis (SMR)**

To identify genes whose expression levels in osteoclasts are associated with BMI traits, we performed an integrative analysis of an osteoclast eQTL dataset [10] (± 1 Mb) using the Summary-data-based Mendelian Randomisation (SMR) software [11] with our GWAS and mtCOJO results. SMR analysis detects dual association signals in the GWAS and eQTL datasets by testing for associations between gene expression level and traits of interest at the top associated eQTL for each gene. SMR detects pleiotropic effects by performing a heterogeneity in dependent instruments (HEIDI) test, which compares the association signals for nearby co-inherited markers in the GWAS and eQTL datasets. A significant HEIDI test indicates heterogeneity in the association profiles of the two datasets, thereby suggesting that the association signals seen in each dataset are less likely to be driven by the same causal variant. A Bonferroni multiple-testing corrected significance threshold of $P < 4.7 \times 10^{-5}$ was used for the SMR test, while a conservative significance threshold of $P < 0.05$ was set for the HEIDI test as an indicator of heterogeneity.

1 <https://www.sanger.ac.uk/resources/downloads/human/hapmap3.html> (

2 Price, A. L. *et al.* Principal components analysis corrects for stratification in genome-wide association studies. *Nat Genet* **38**, 904-909, doi:10.1038/ng1847 (2006).

3 Price, A. L. *et al.* Long-range LD can confound genome scans in admixed populations. *Am J Hum Genet* **83**, 132-135, doi:10.1016/j.ajhg.2008.06.005 (2008).

4 Carithers, L. J. & Moore, H. M. The Genotype-Tissue Expression (GTEx) Project. *Biopreserv Biobank* **13**, 307-308, doi:10.1089/bio.2015.29031.hmm (2015).

5 Stancakova, A. *et al.* Hyperglycemia and a common variant of GCKR are associated with the levels of eight amino acids in 9,369 Finnish men. *Diabetes* **61**, 1895-1902, doi:10.2337/db11-1378 (2012).

6 Stancakova, A. *et al.* Changes in insulin sensitivity and insulin release in relation to glycemia and glucose tolerance in 6,414 Finnish men. *Diabetes* **58**, 1212-1221, doi:10.2337/db08-1607 (2009).

7 Wright, F. A. *et al.* Heritability and genomics of gene expression in peripheral blood. *Nat Genet* **46**, 430-437, doi:10.1038/ng.2951 (2014).

8 Nuotio, J. *et al.* Cardiovascular risk factors in 2011 and secular trends since 2007: the Cardiovascular Risk in Young Finns Study. *Scand J Public Health* **42**, 563-571, doi:10.1177/1403494814541597 (2014).

9 Raitakari, O. T. *et al.* Cohort profile: the cardiovascular risk in Young Finns Study. *Int J Epidemiol* **37**, 1220-1226, doi:10.1093/ije/dym225 (2008).

10 Mullin, B. H. *et al.* Expression Quantitative Trait Locus Study of Bone Mineral Density GWAS Variants in Human Osteoclasts. *J Bone Miner Res* **33**, 1044-1051, doi:10.1002/jbmr.3412 (2018).

11 Zhu, Z. *et al.* Integration of summary data from GWAS and eQTL studies predicts complex trait gene targets. *Nat Genet* **48**, 481-487, doi:10.1038/ng.3538 (2016).
