## Supplementary material for "Genome-wide analyses point to differences in genetic architecture of BMI between tall and short people"

### Locus zoom plots for candidate SNPs (in Supplementary table 2)

Supplementary material

Bochao Lin

### 1p36.33 (top SNP rs11260621, an intron variant in *GNB1*).

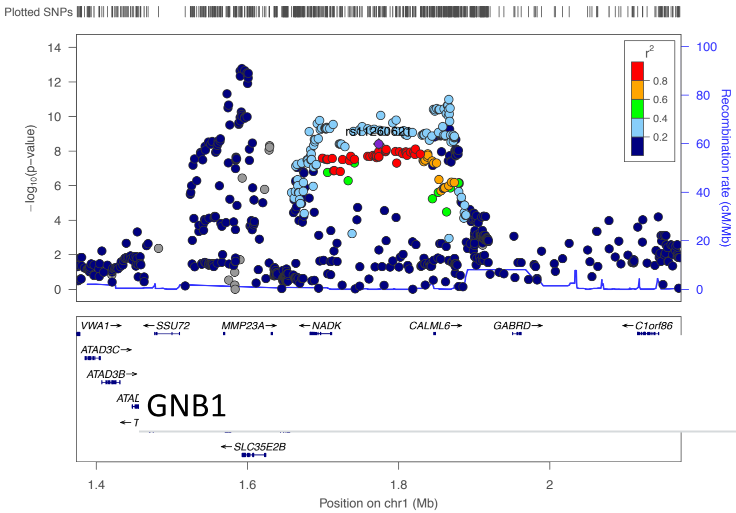


Total population

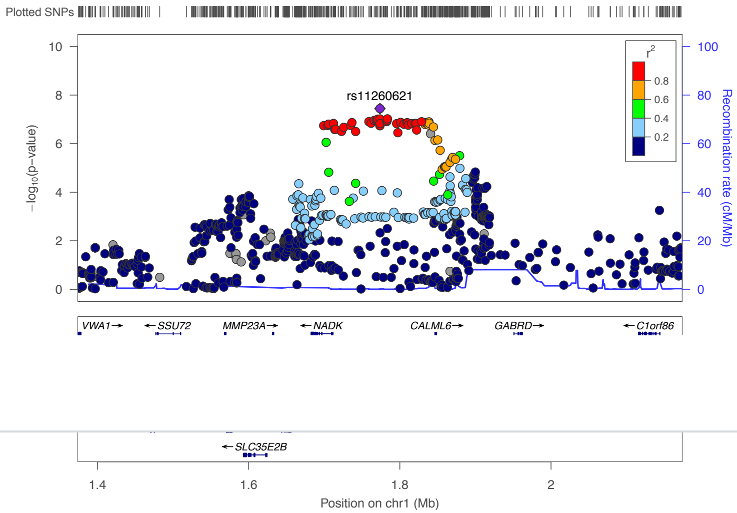


Short people

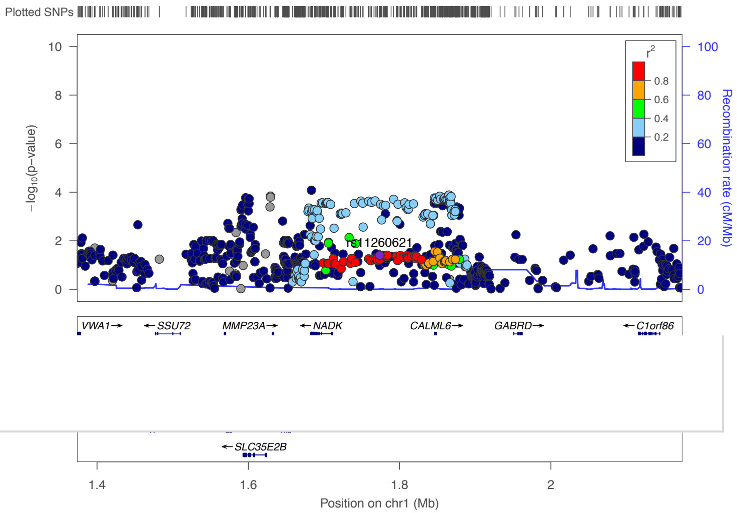


Tall people

### 1p35.1 (top SNP is rs76066944, an intron variant in *YARS1*).

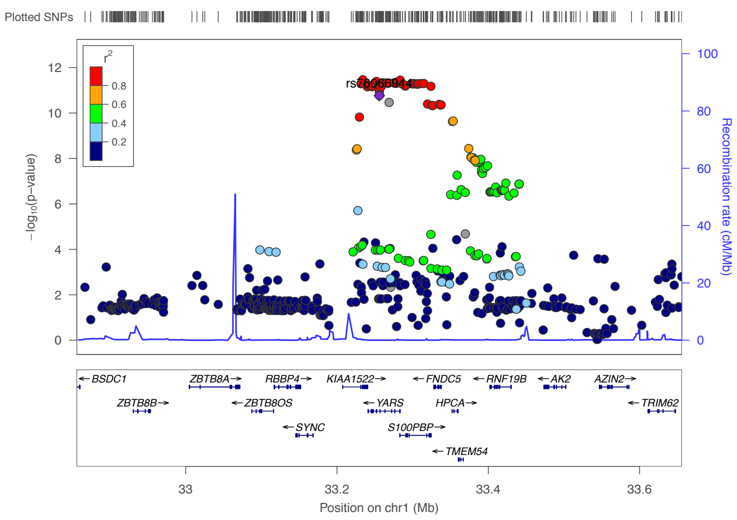


Total population

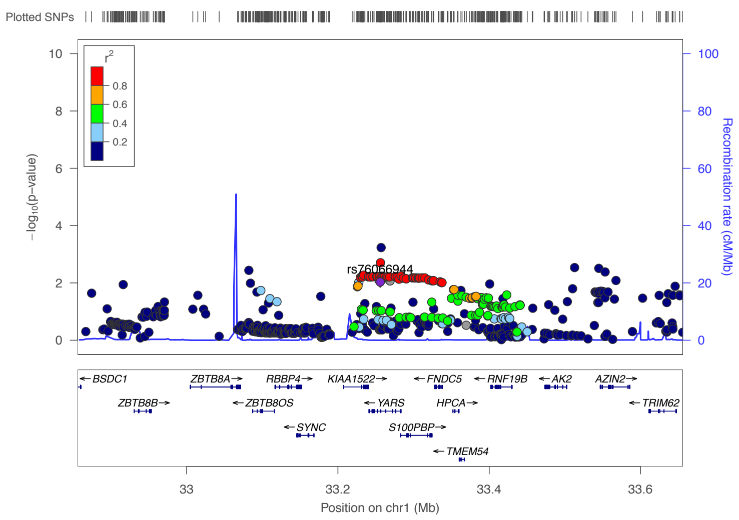


Short people

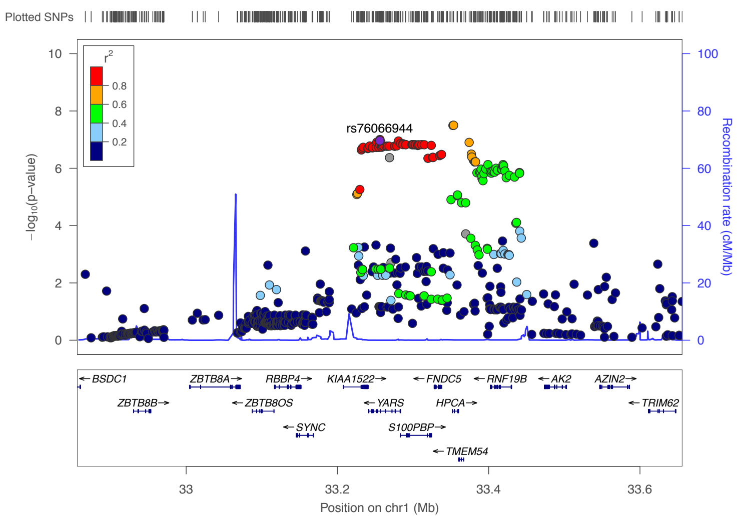


Tall people

### 1p35.1 (top SNP is rs942244, an intron variant in *HPCA*).

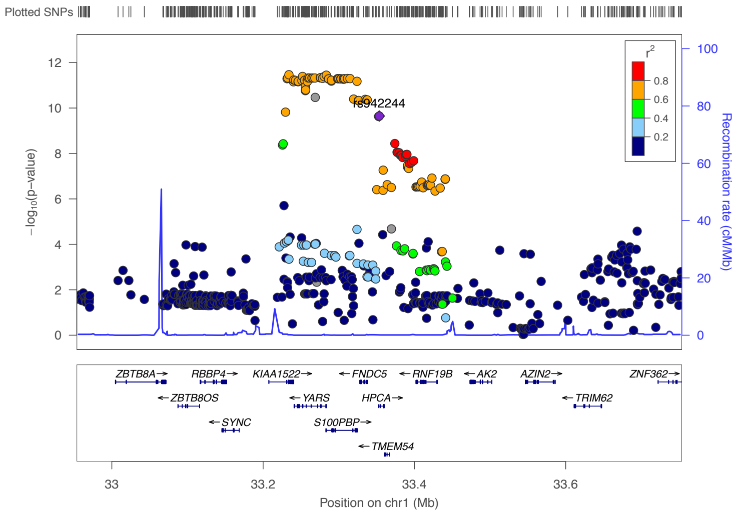


Total population

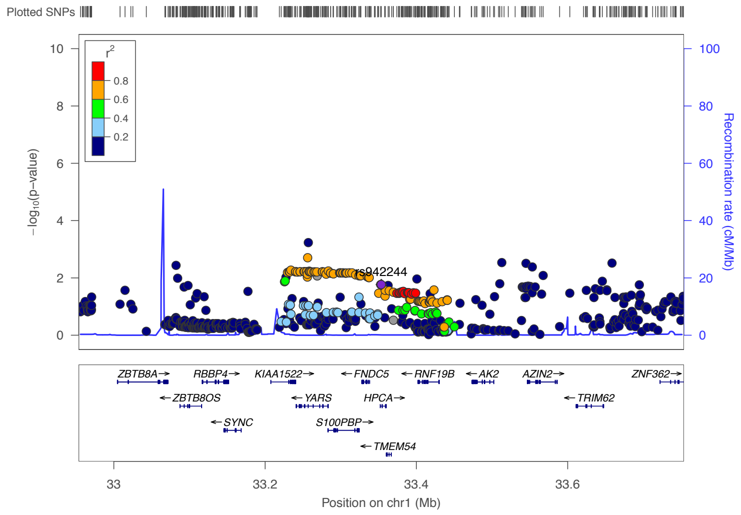


Short people

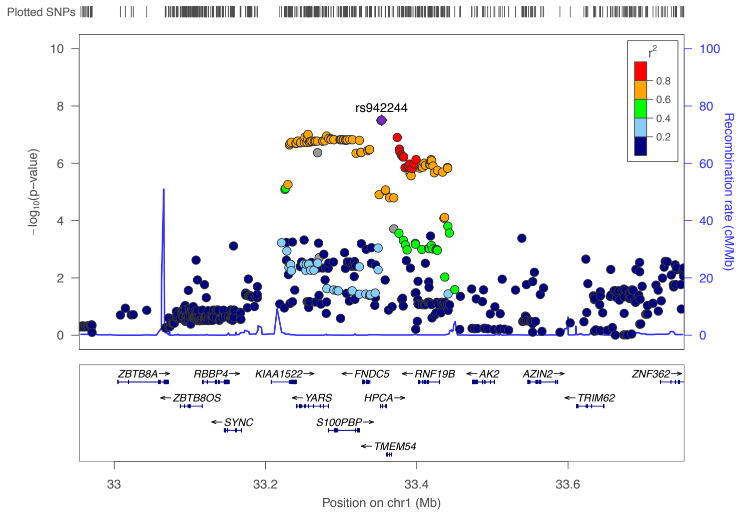


Tall people

### 1p31.1 (top SNP is rs12409142, a down stream gene variant 4.6kb *USP33*).

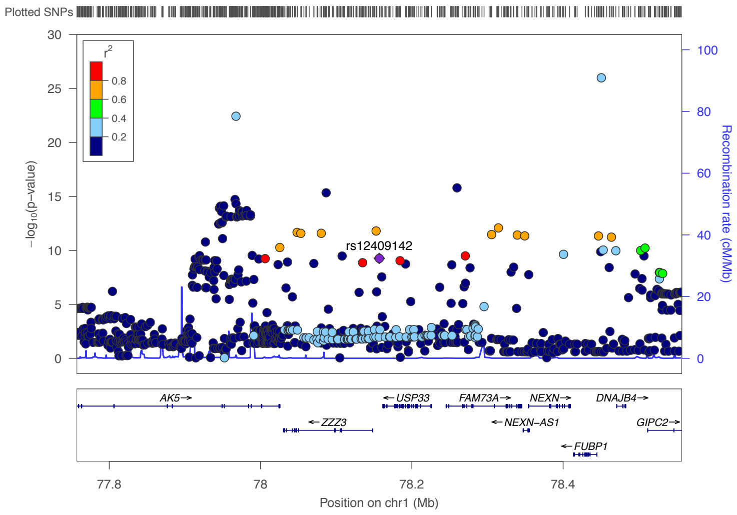


Total population

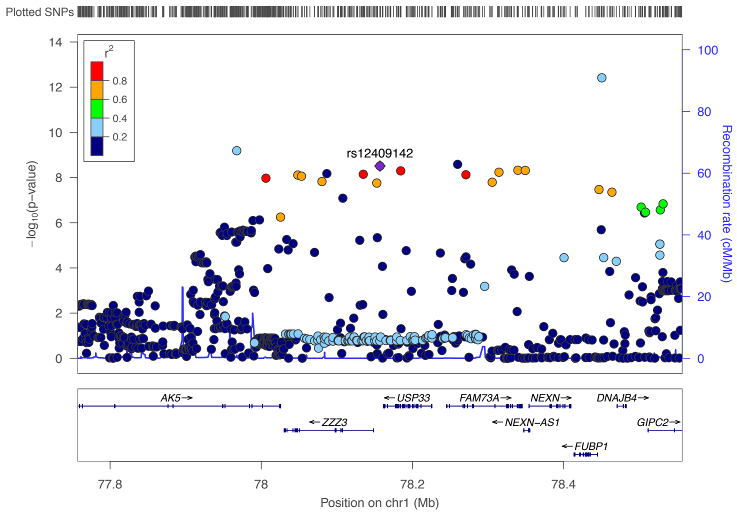


Short people

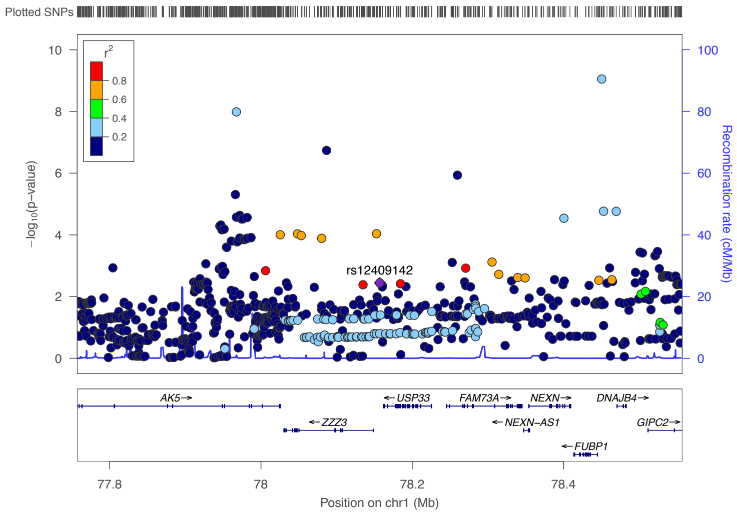


Tall people

### 1q25.2 (top SNP is rs522367 an upstream gene variant for *SEC16B*).

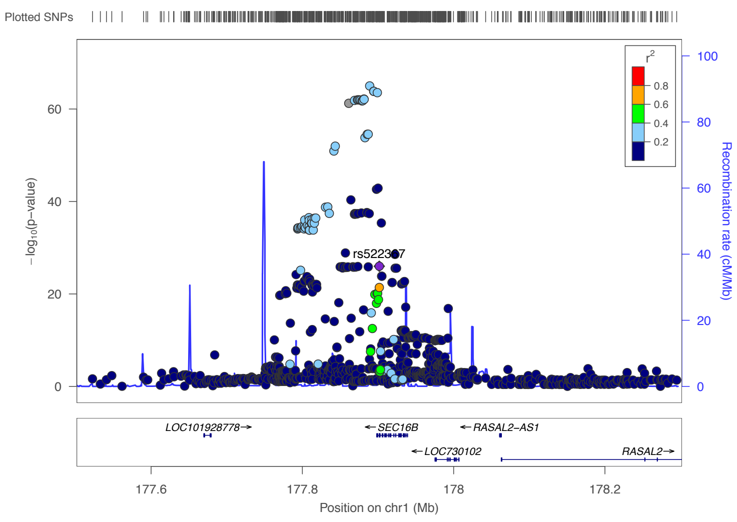


Total population

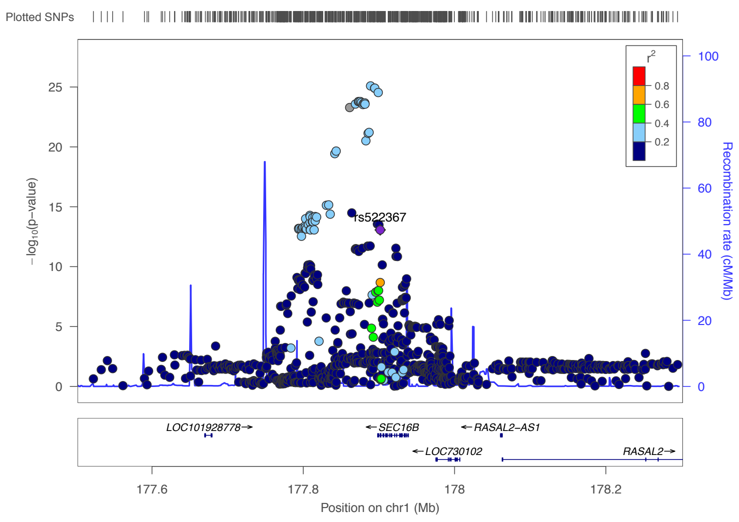


Short people

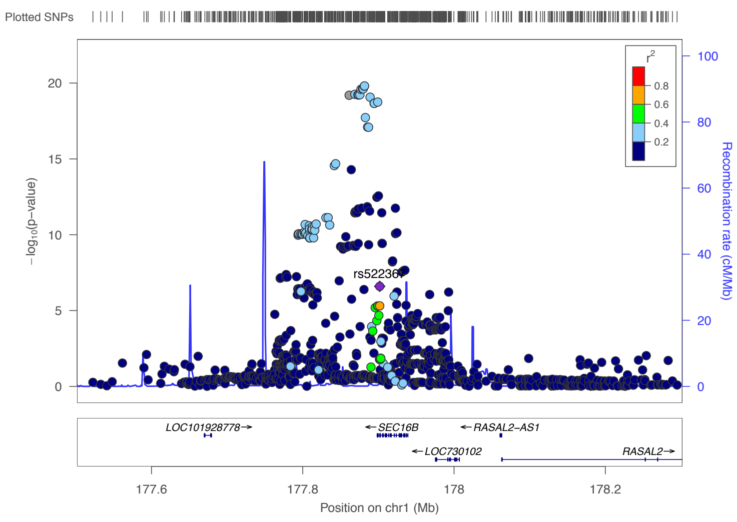


Tall people

### 2p25.3 (top SNP is Rs141224959, an intergenic variant near by *FAM150B*).

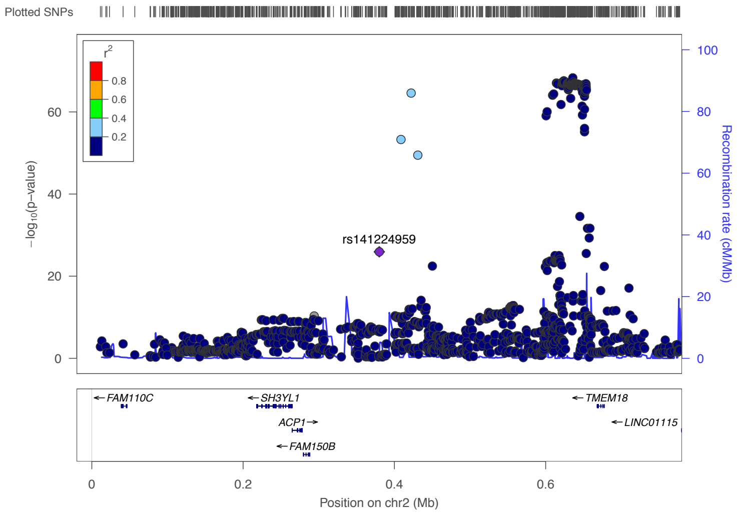


Total population

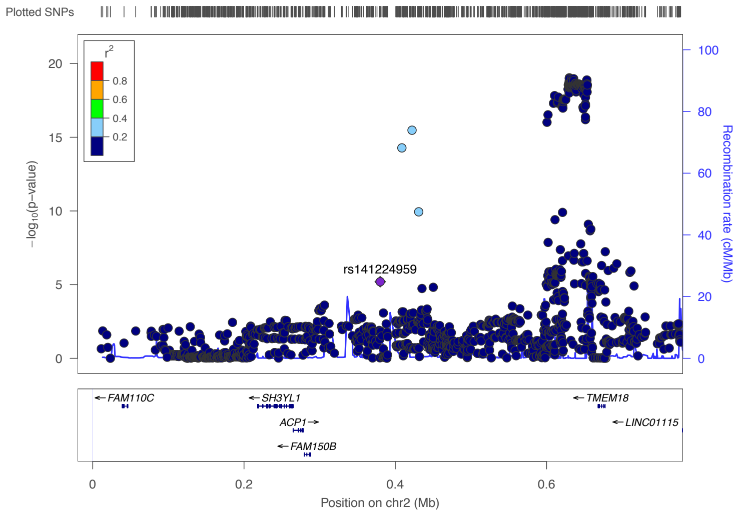


Short people

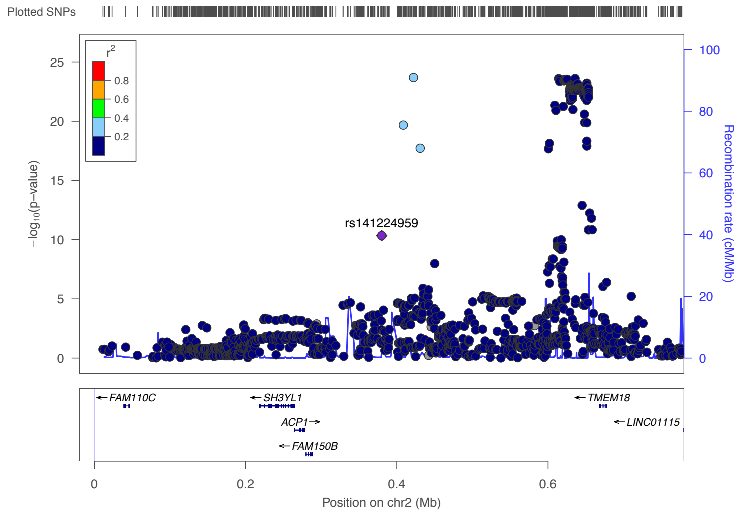


Tall people

### 2p25.3 (top SNP is rs4854330, an intergenic variant).

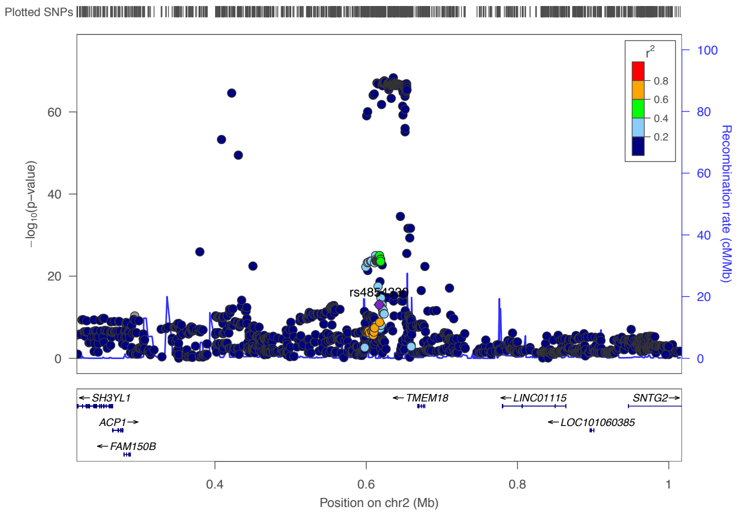


Total population

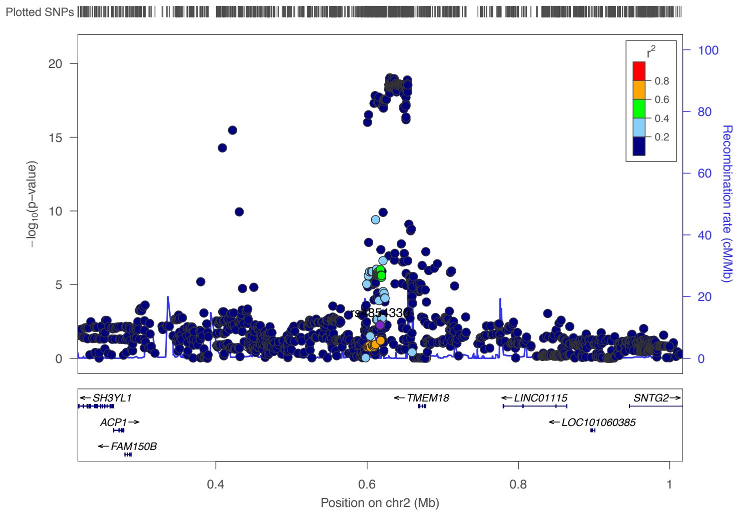


Short people

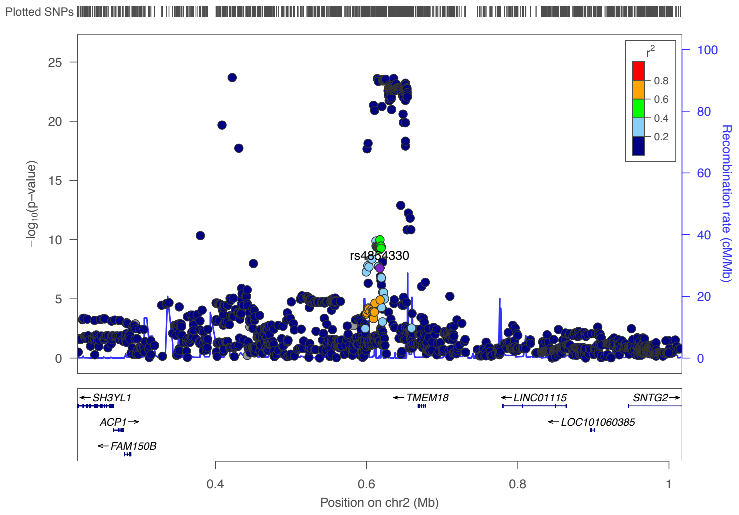


Tall people

### 2p23.3 (top SNP is rs10176214, an upstream gene variant of *ADCY3*).

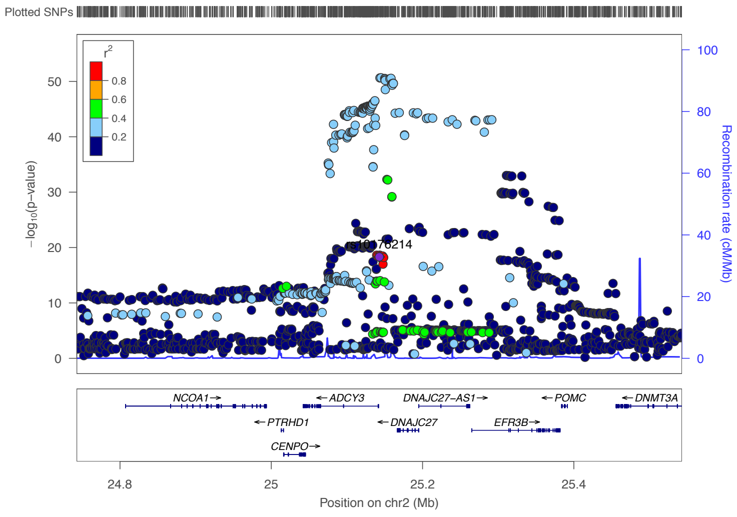


Total population

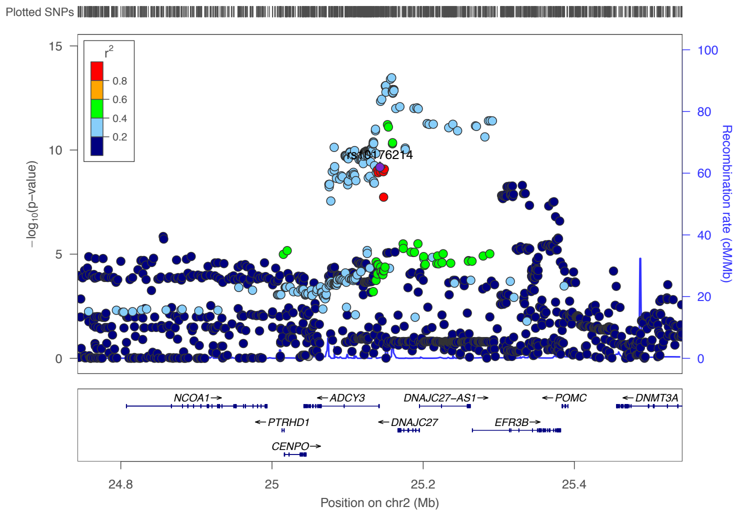


Short people

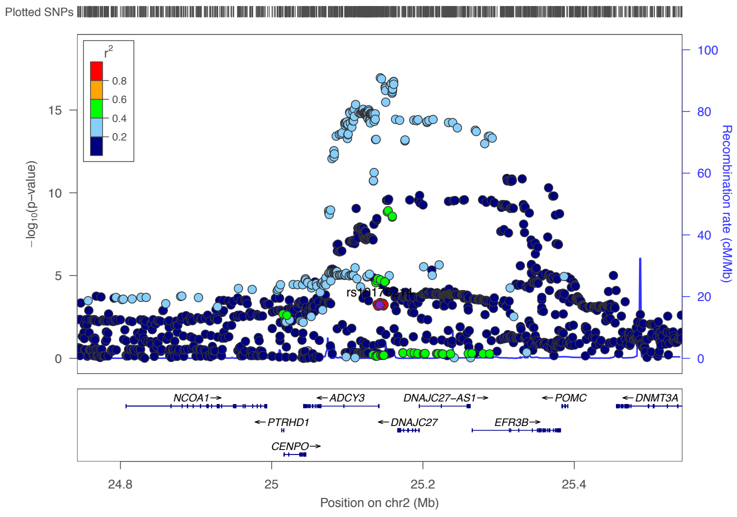


Tall people

### 2p23.3( the top SNP is rs76286777, a non coding transcript exon variant on *DNAJC27*).

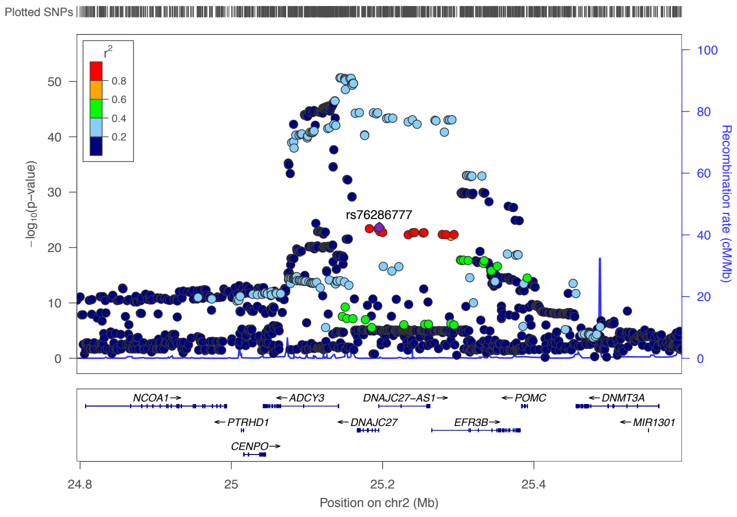


Total population

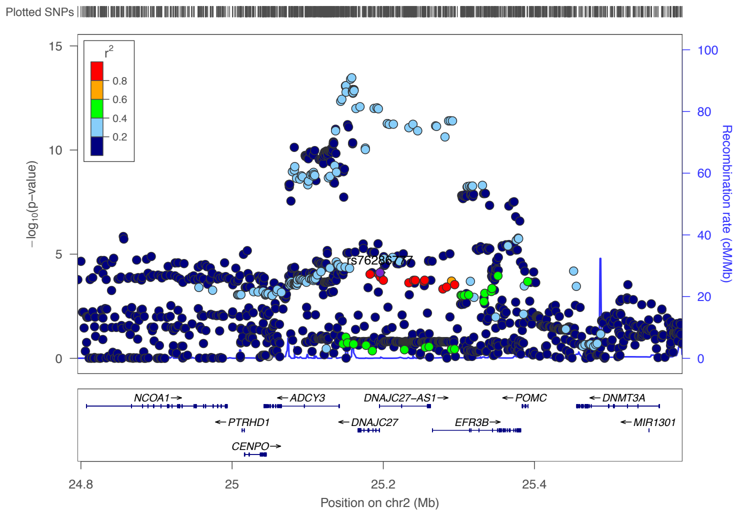


Short people

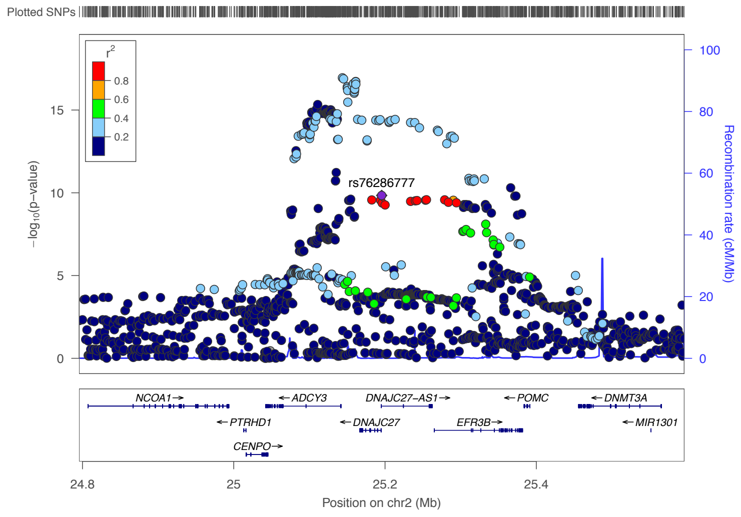


Tall people

### 2q11.2 (top SNP is rs190441457, an intron variant in *AFF3*).

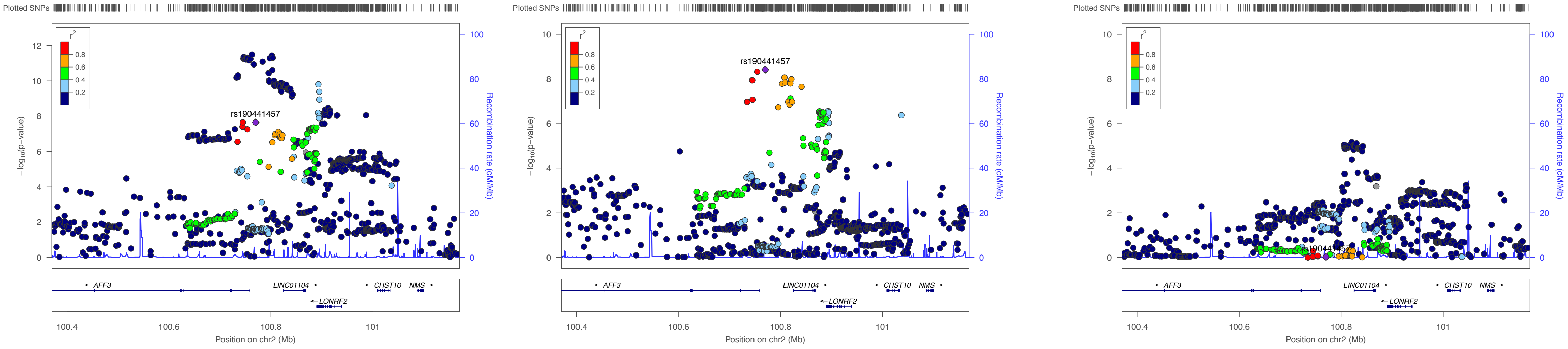


Total population

Short people

Tall people

### 2q24.3 (Top SNP is rs12692738, an intron variant *COBLL1*).

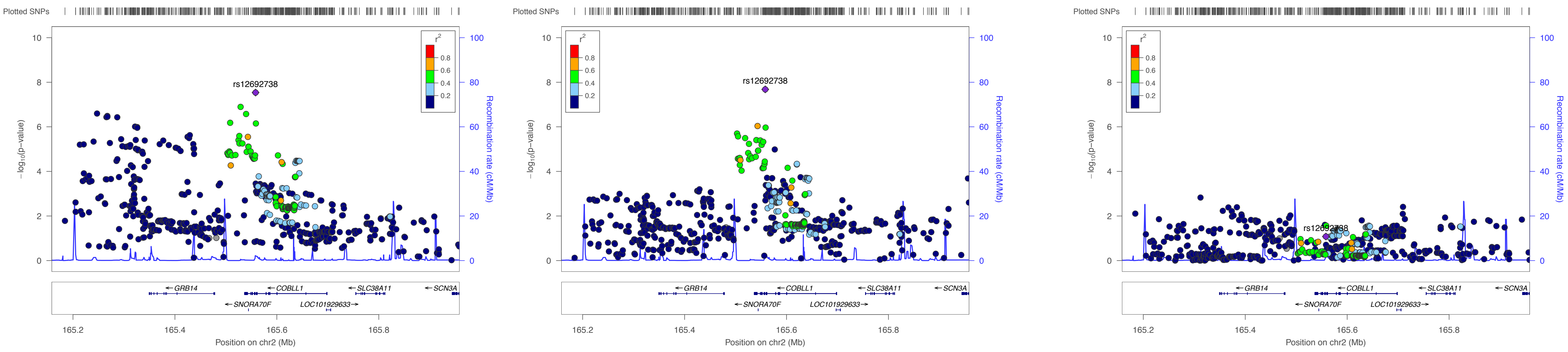


Total population

Short people

Tall people

### 2q31.3 (rs6723591, an intron variant in *SCHLAP1*).

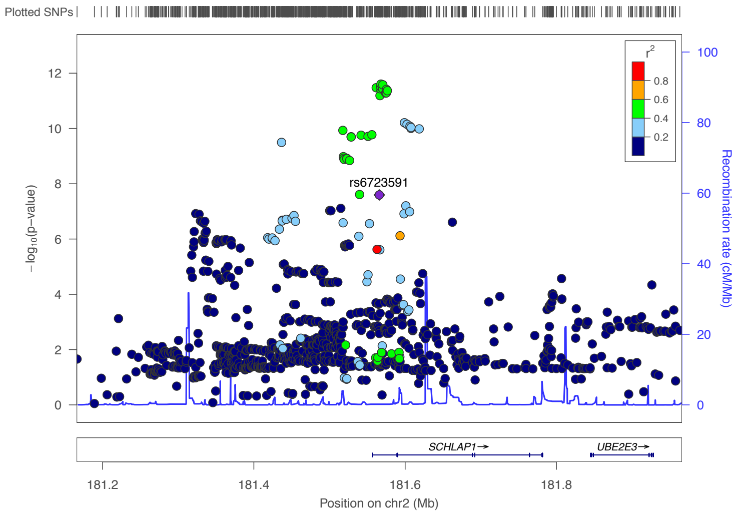


Total population

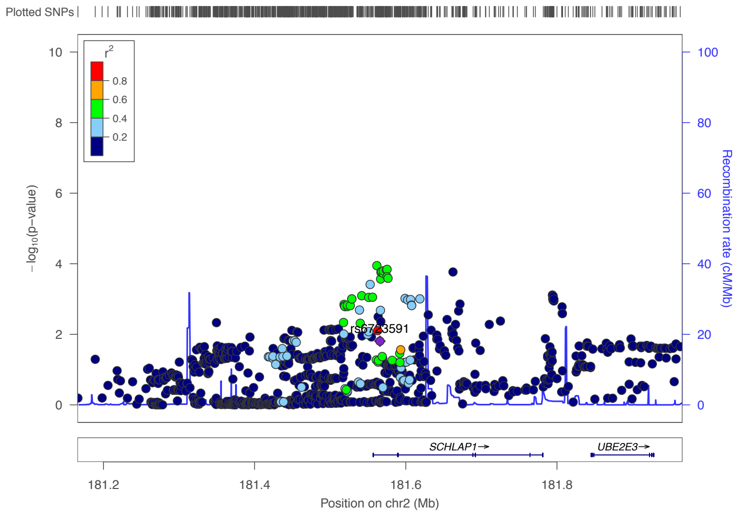


Short people

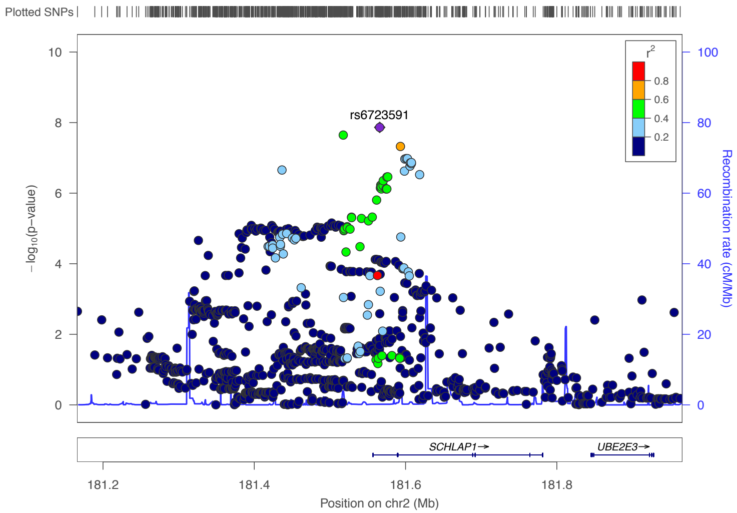


Tall people

### 2q33.3 (top SNP is rs12694002, an intergenic variant nearby *PARD3B*).

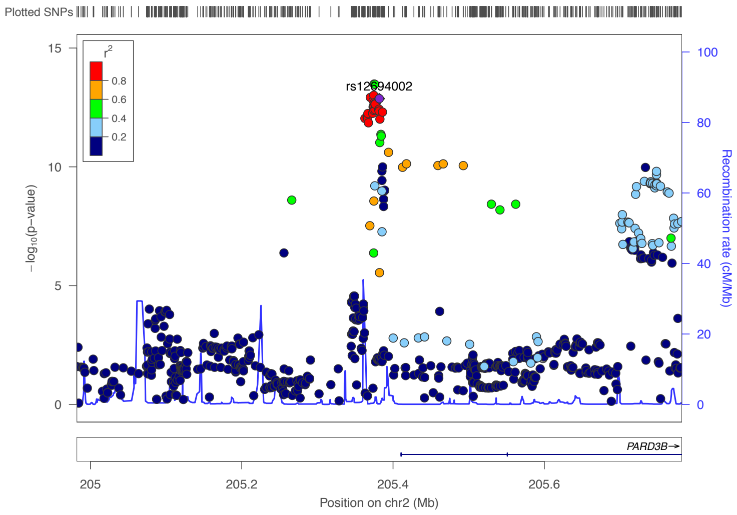


Total population

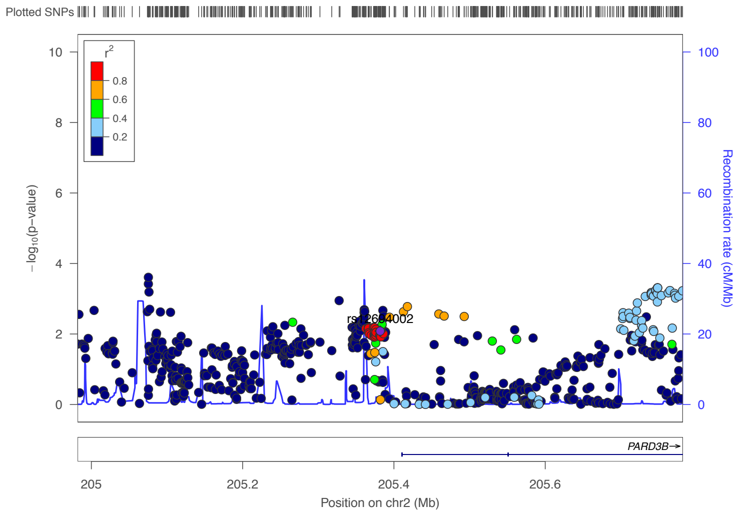


Short people

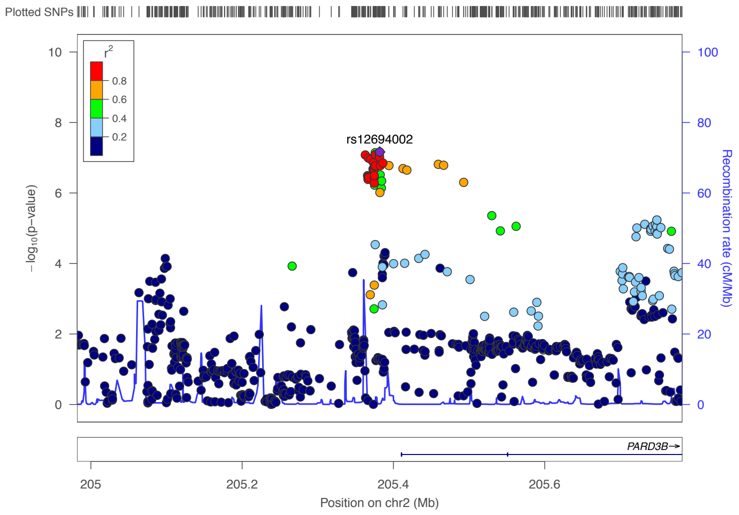


Tall people

### 2q36.3 (top SNP is rs7558924, an intron variant in *DNER*).

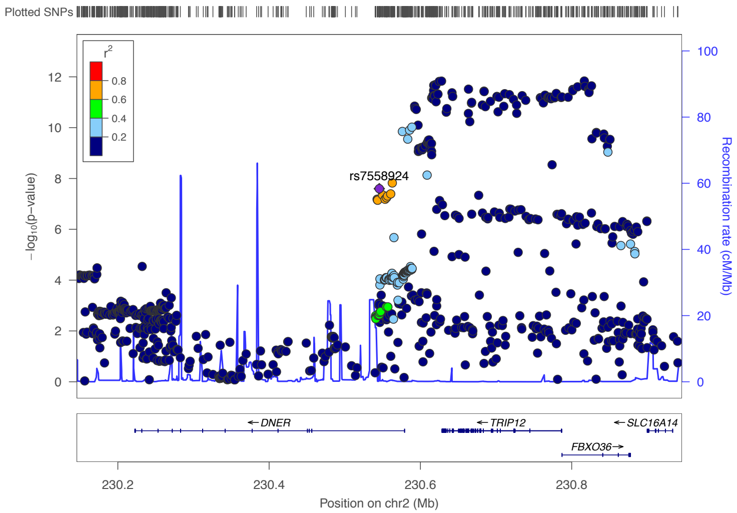


Total population

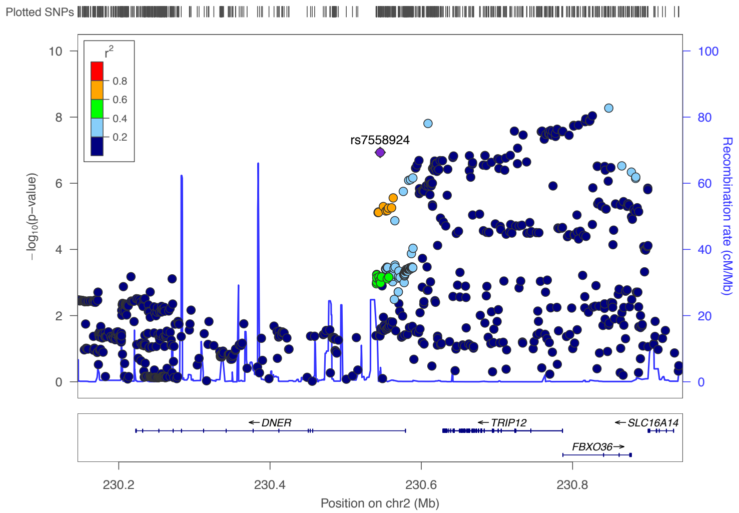


Short people

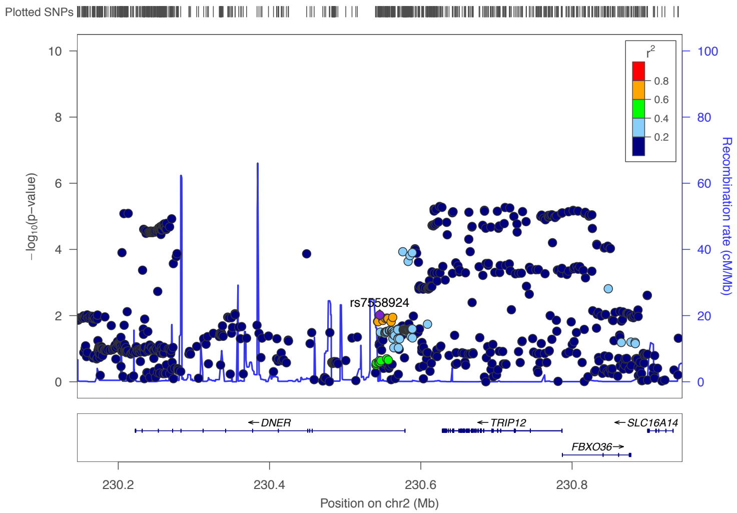


Tall people

### 2q36.3 (top SNP rs7577278 intron variant in *FBXO36*).

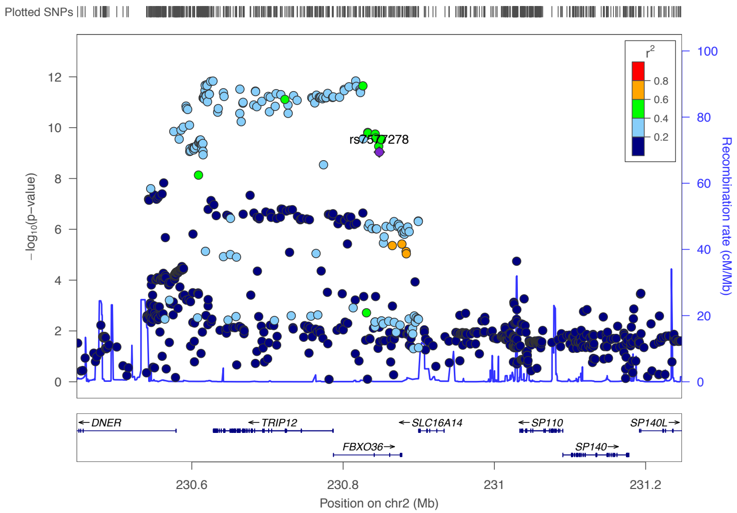


Total population

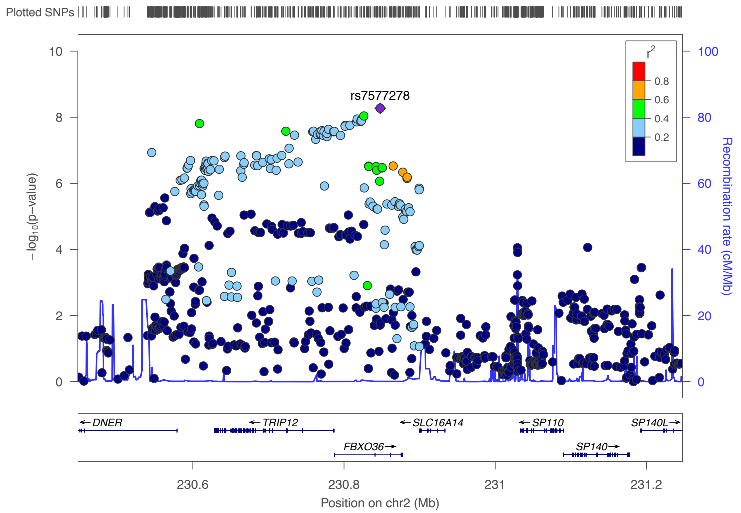


Short people

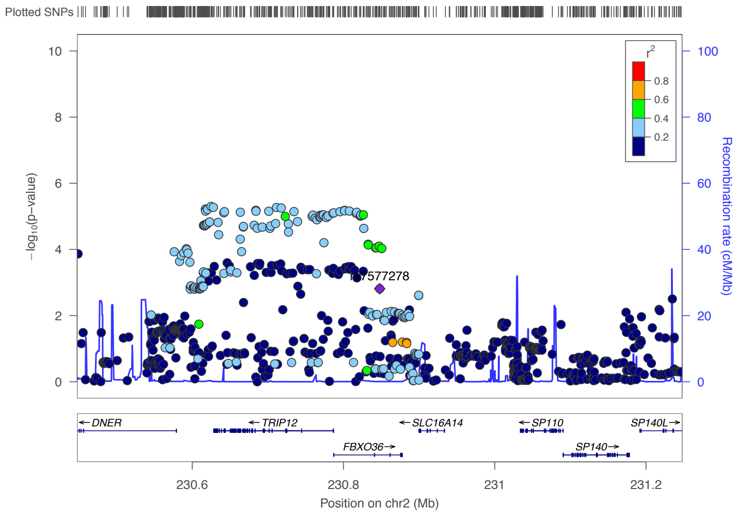


Tall people

### 3p21.31 (rs9843653 a downstream gene variant *MST1R*).

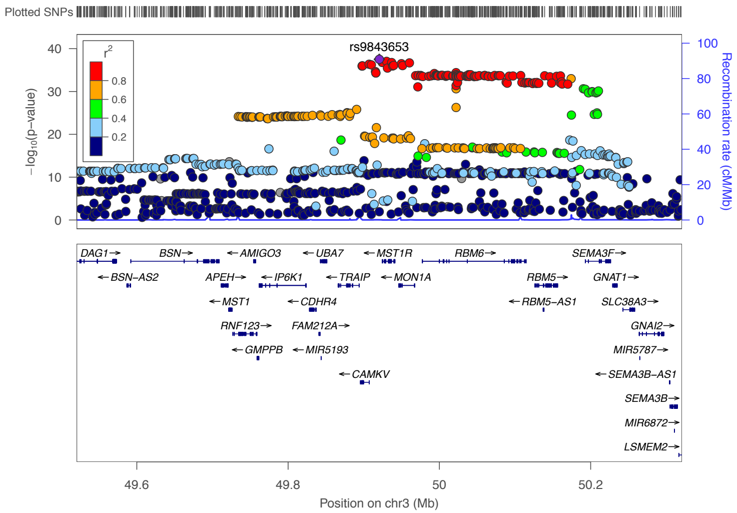


Total population

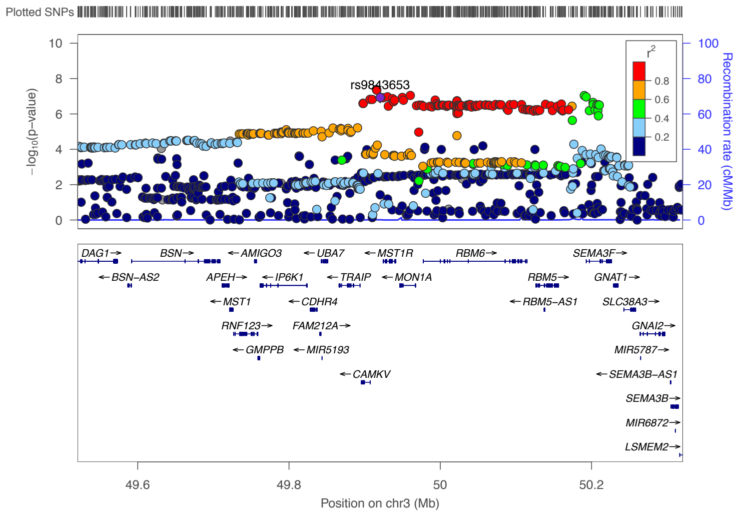


Short people

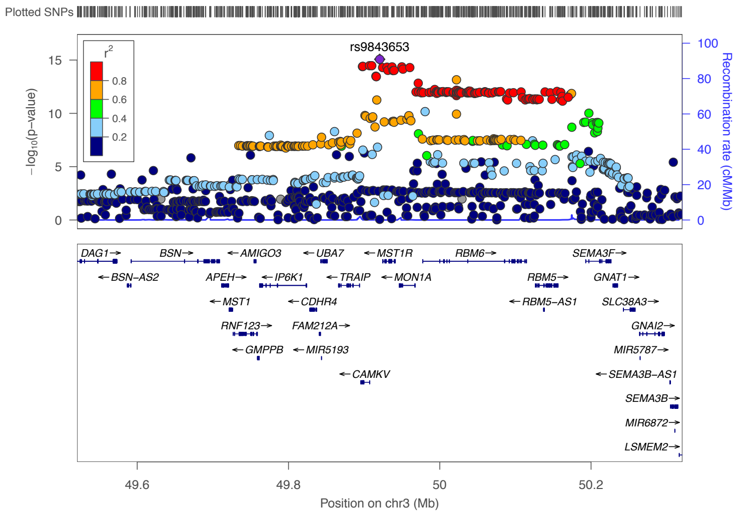


Tall people

### 3p21.31 (top SNP is rs7627910, an upstream gene variant in *MON1A*).

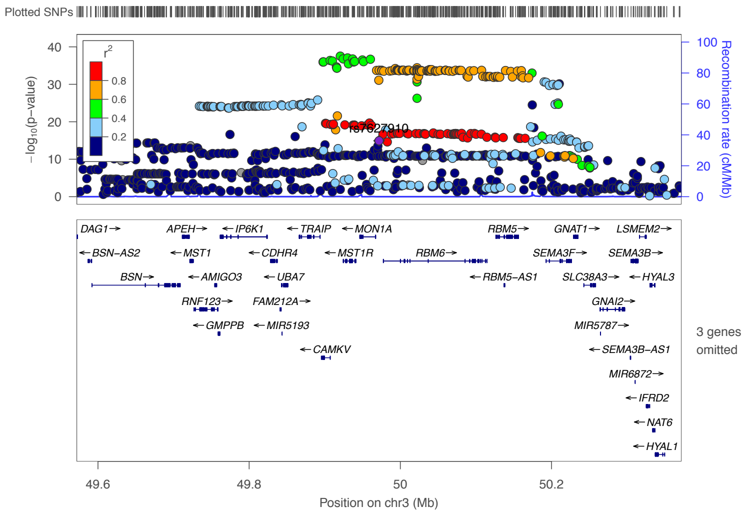


Total population

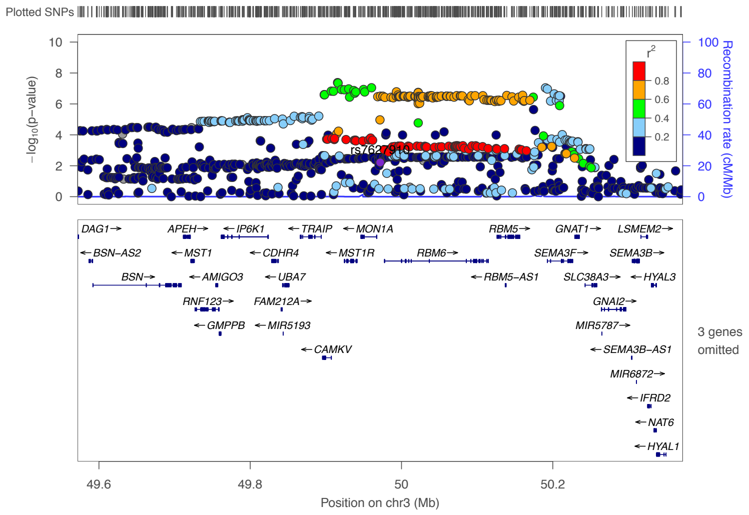


Short people

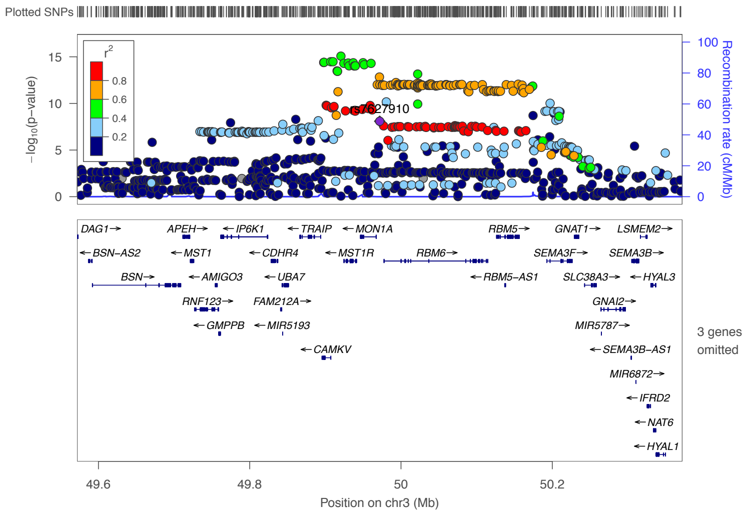


Tall people

### 3q22.2 (top SNP is rs7636391, an intron variant *EPHB1*).

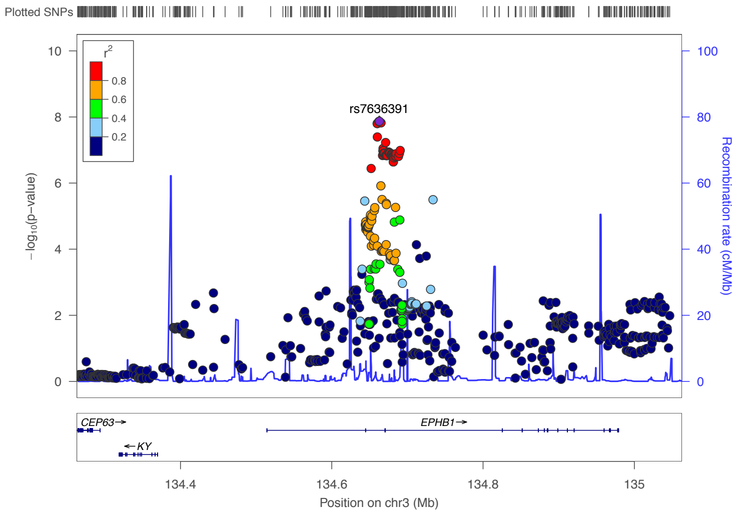


Total population

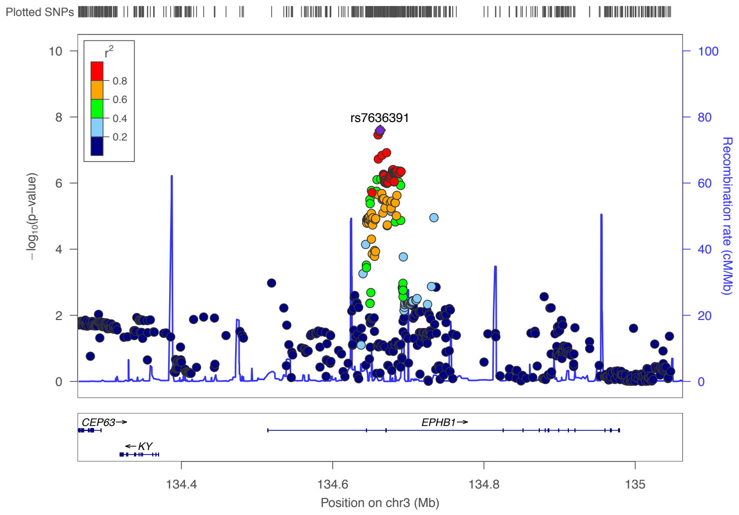


Short people

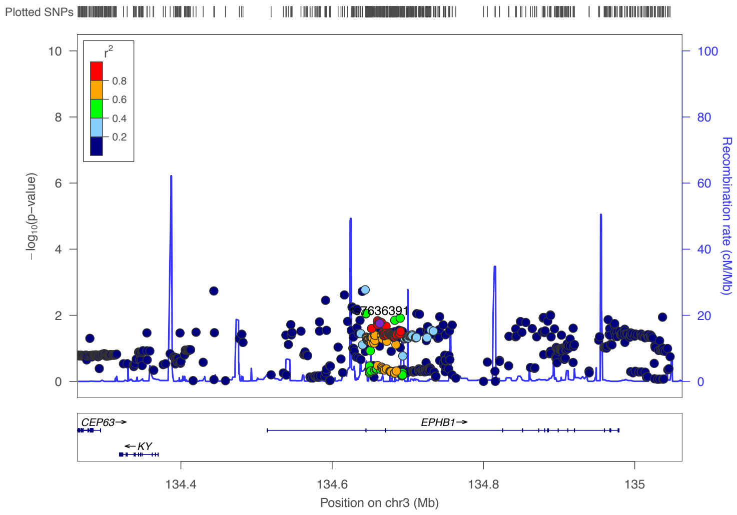


Tall people

### 3q25.2 (rs9851875, an intron variant in *LINC02006*).

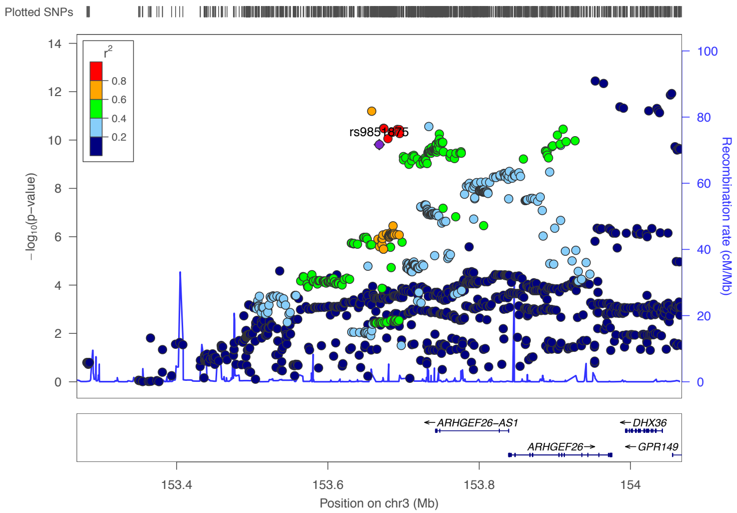


Total population

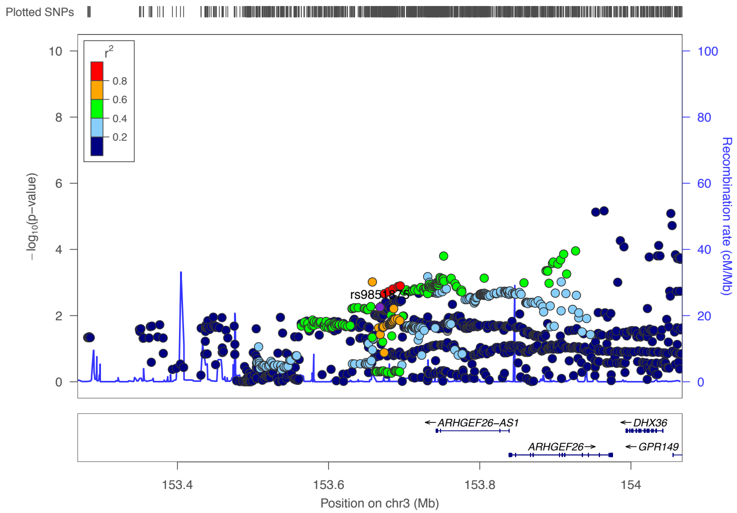


Short people

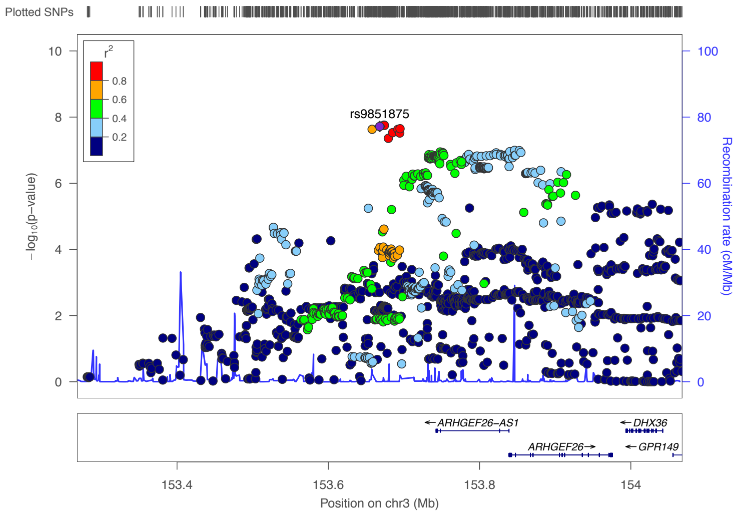


Tall people

### 3q26.1 (top SNP rs4856720, an intron variant *AC112770.1*).

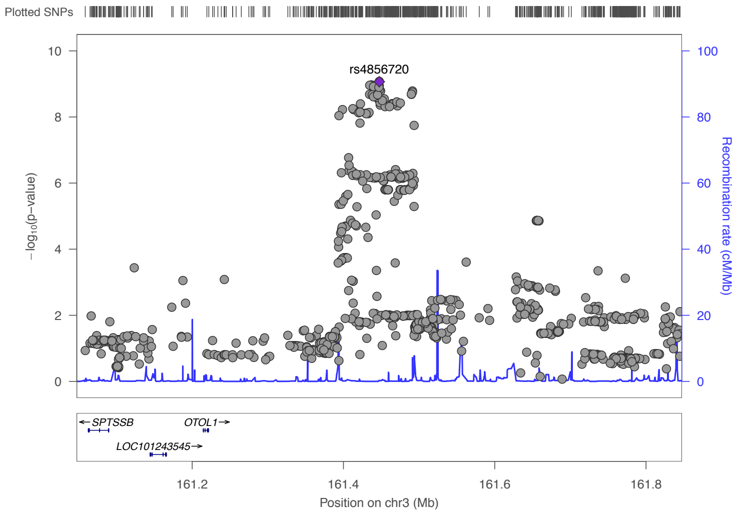


Total population

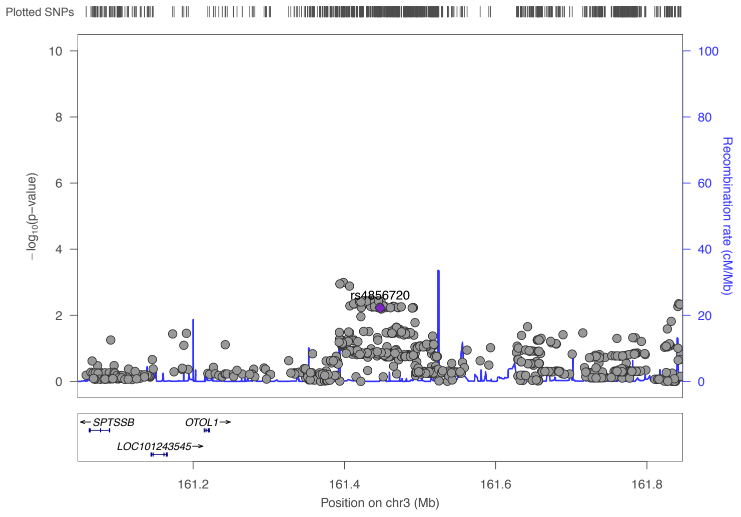


Short people

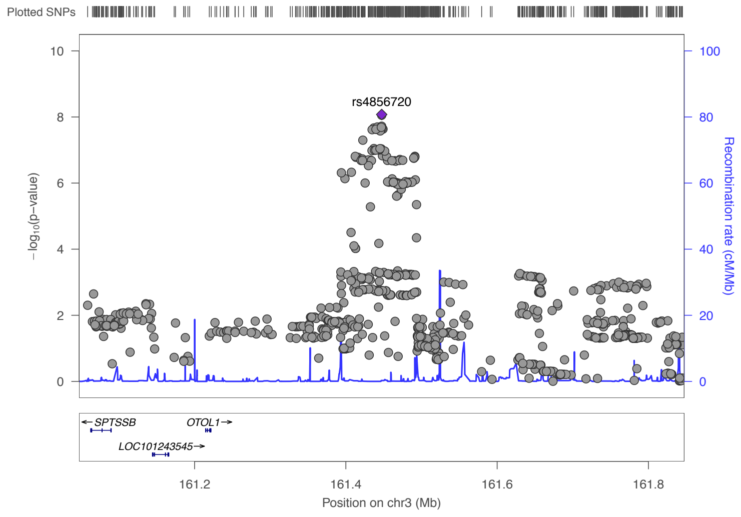


Tall people

### 3q27.2( Top SNP is rs4494964, an intron variant in *ETV5*).

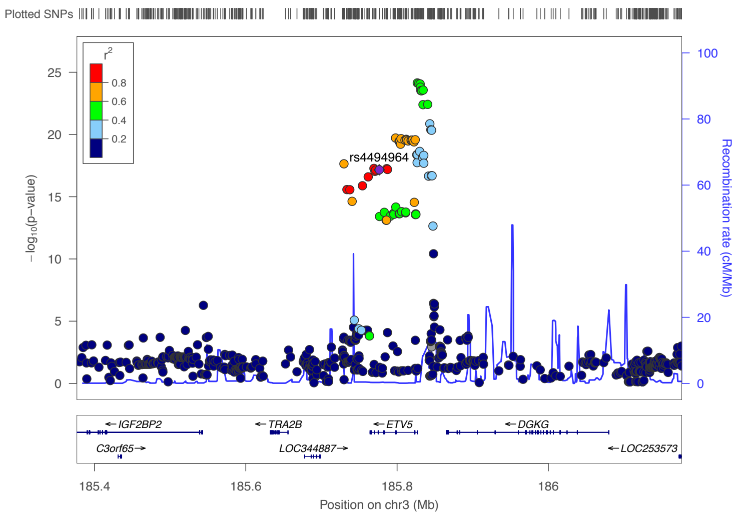


Total population

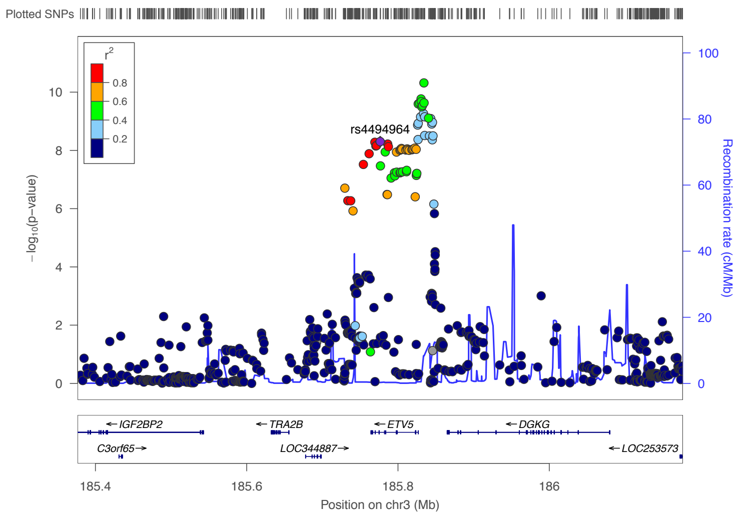


Short people

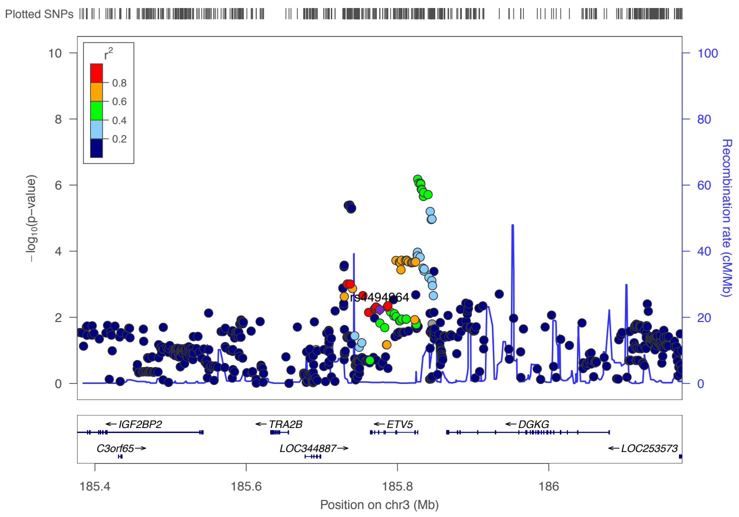


Tall people

### 3q27.2 (top SNP rs7635103, an intron variant in *DGKG*).

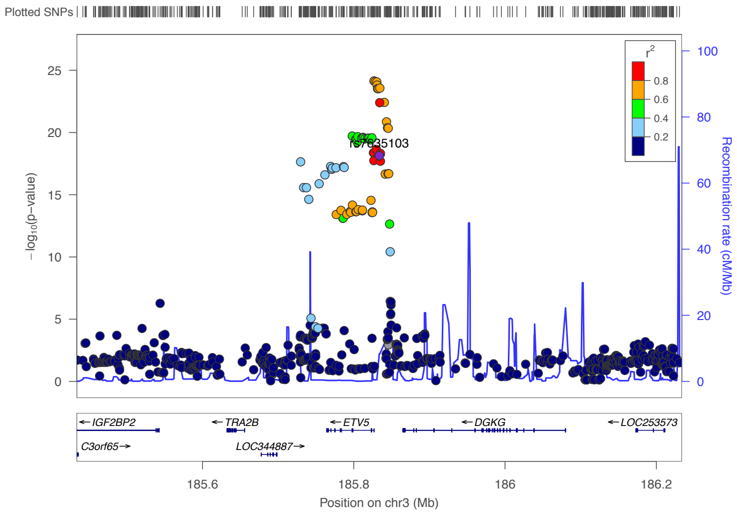


Total population

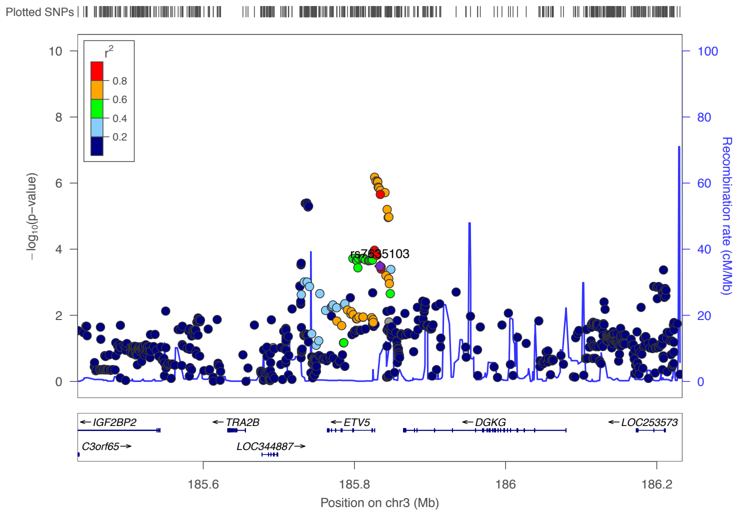


Short people

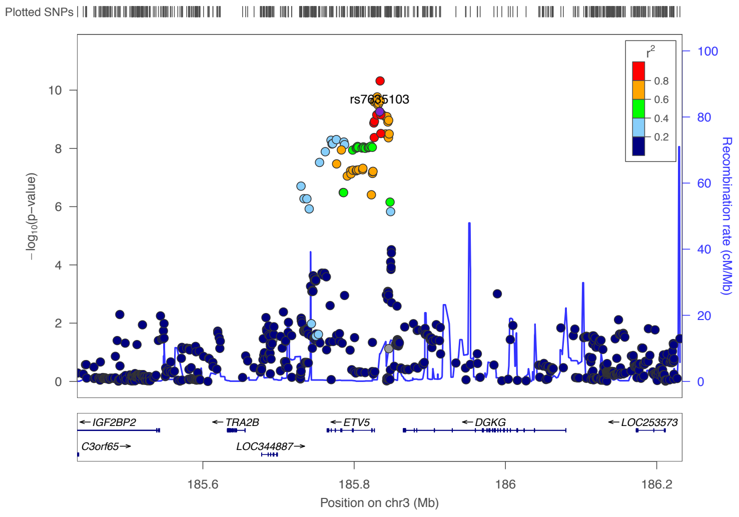


Tall people

### Locus in chromosome 4 (top SNP is rs348492, an intergenic variant).

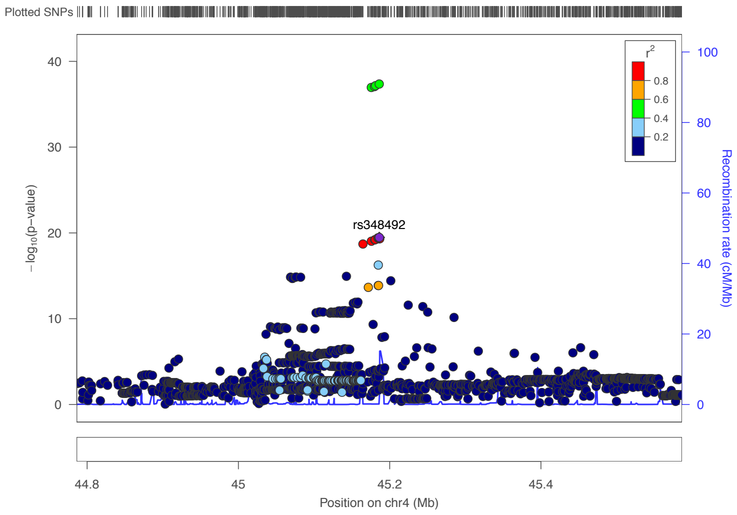


Total population

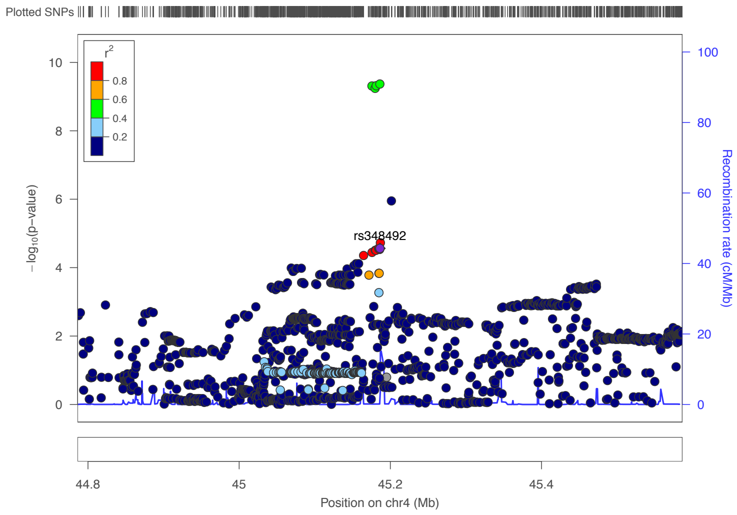


Short people

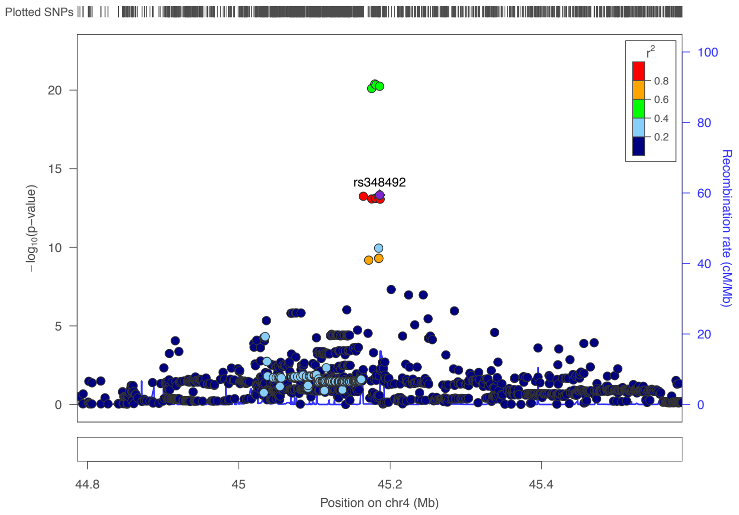


Tall people

### 6p21.3 (top SNP is rs9366858, an intron variant in *PACSIN1*).

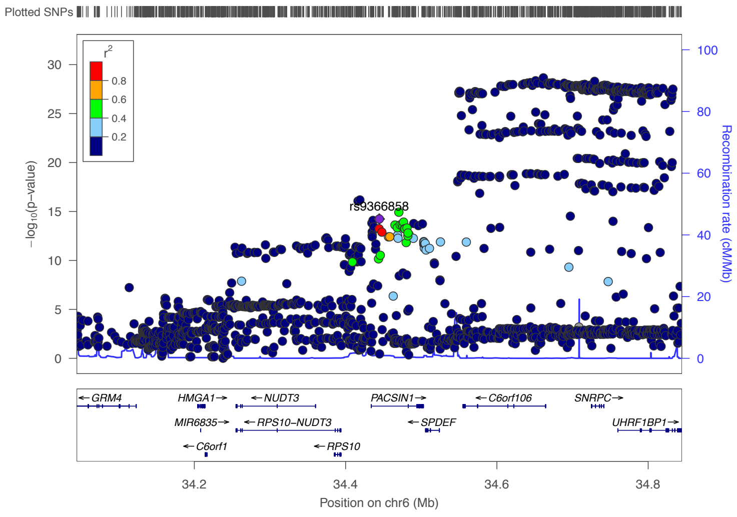


Total population

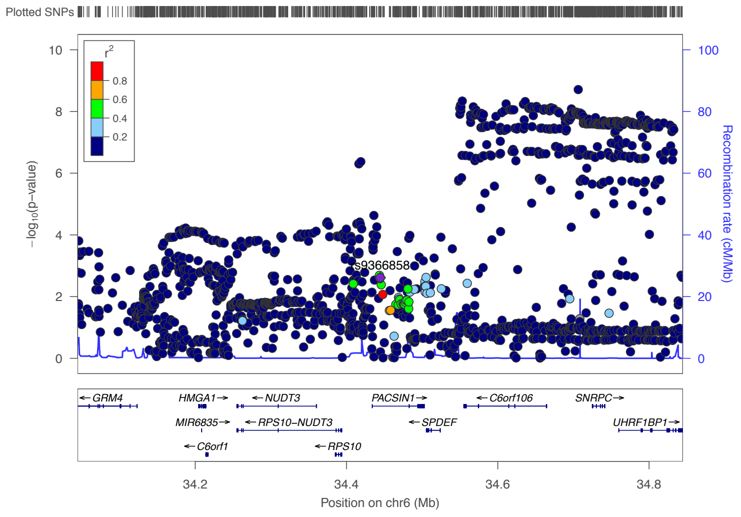


Short people

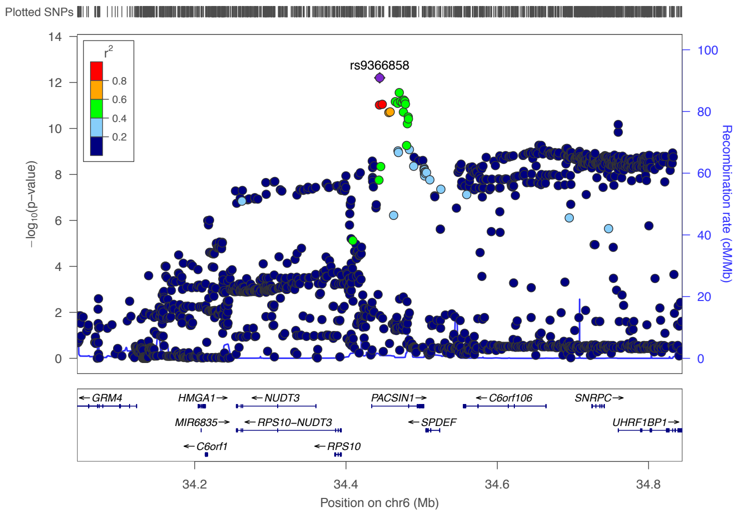


Tall people

### 6p21.3 (top SNP is rs2104332, a regulatory region variant located at *CTCF* binding site and promoter flanking region).

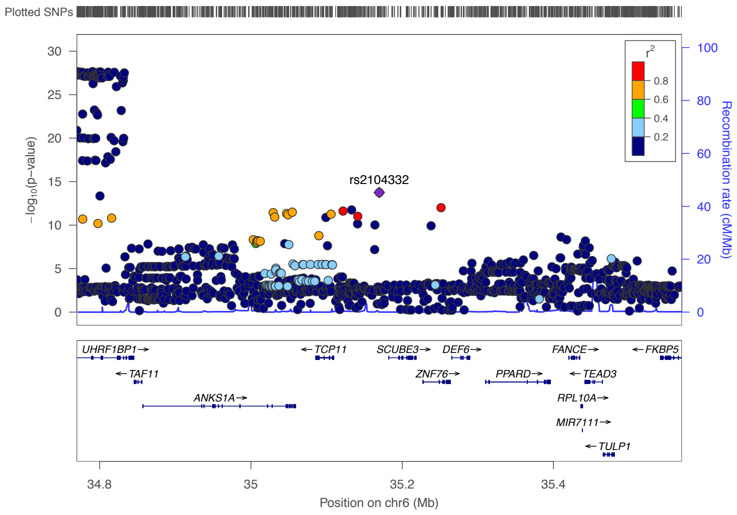


Total population

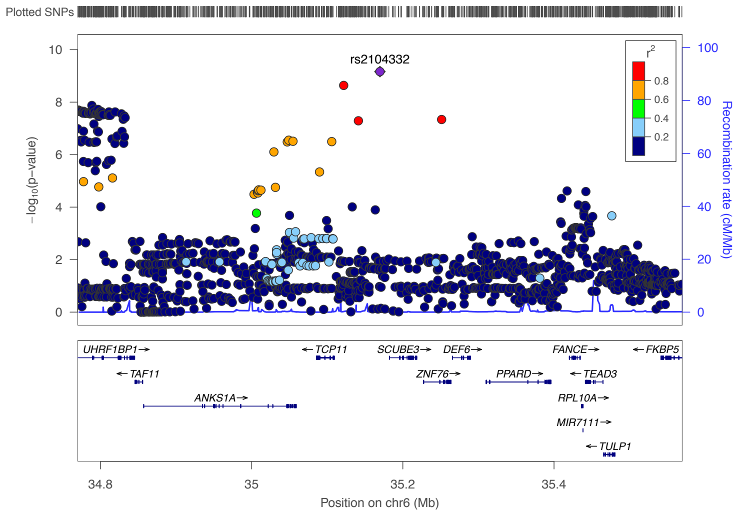


Short people

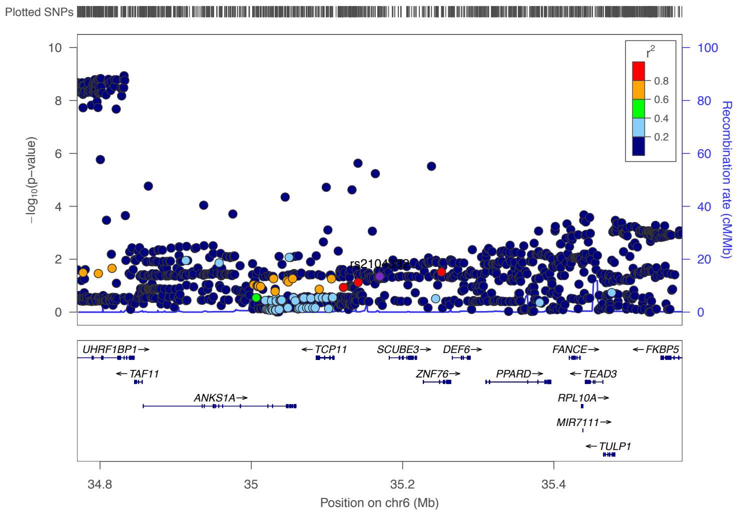


Tall people

### 6p21.2 (top SNP is rs34045288, an intron variant in *LRFN2*).

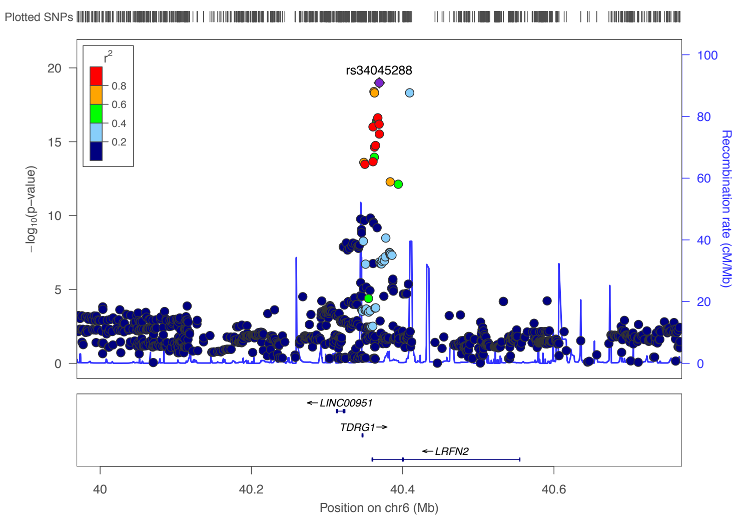


Total population

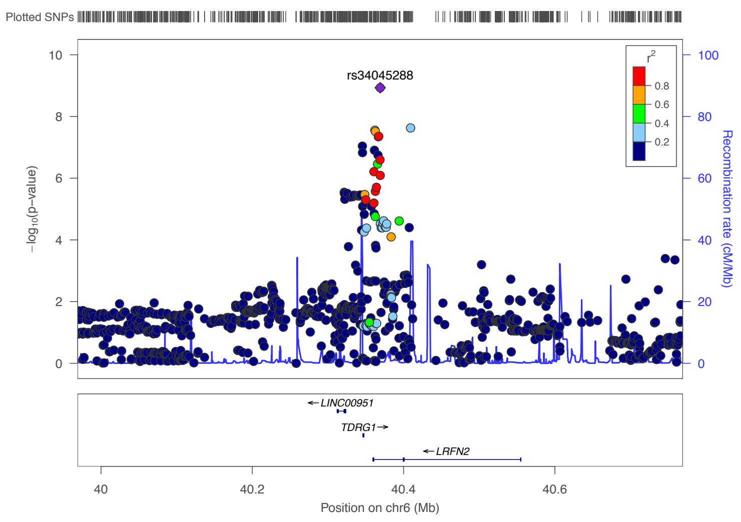


Short people

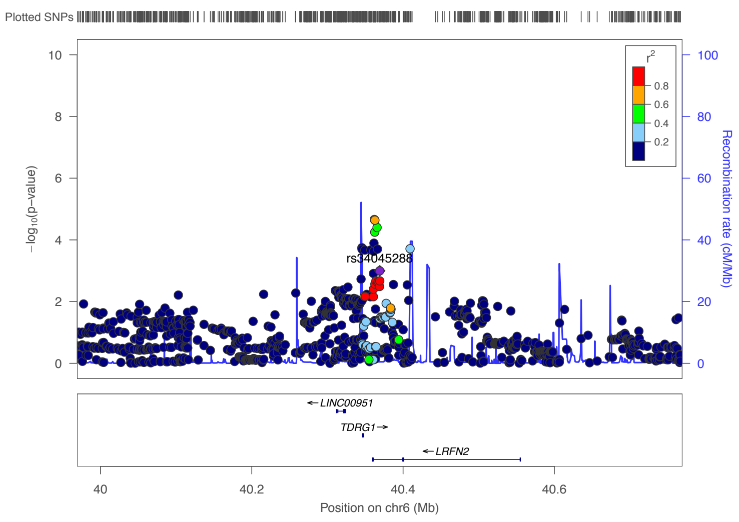


Tall people

### 7q11.23 (top SNP is rs4084934, an regulatory region variant located at *UPK3B* promoter flanking region).

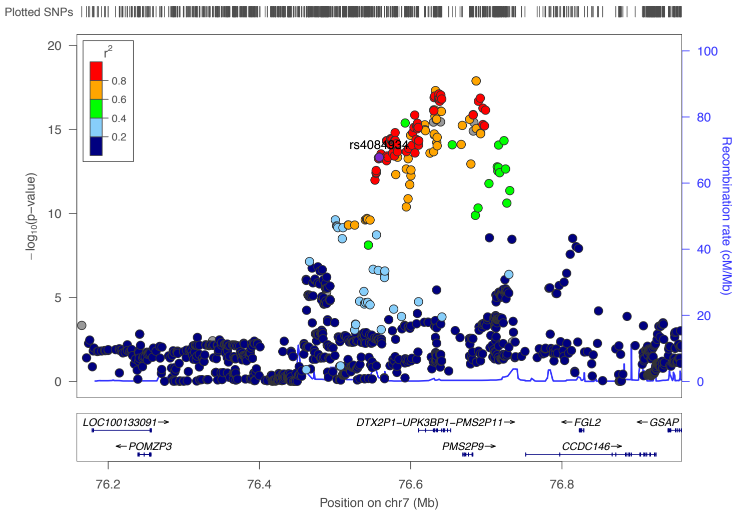


Total population

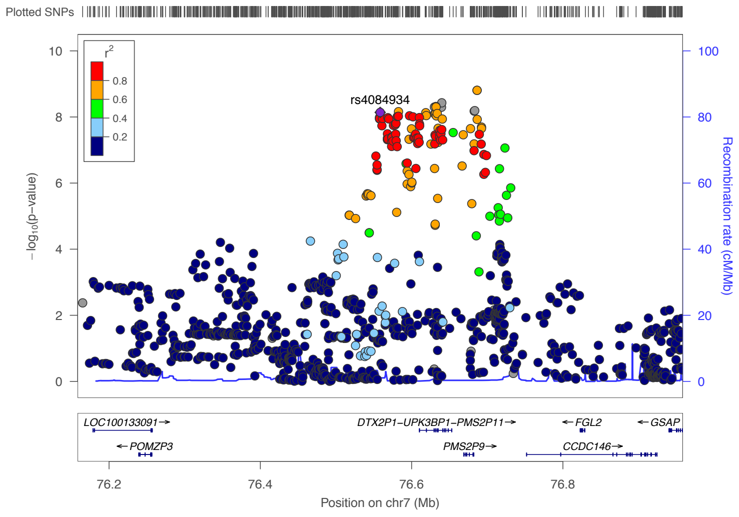


Short people

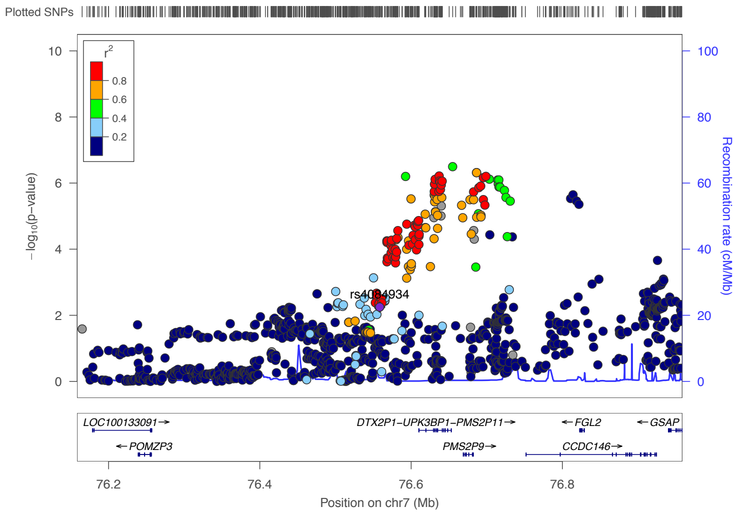


Tall people

### 7q31.1  (top SNP is rs7798292, an intergenic variant  nearby *LINC00998*).

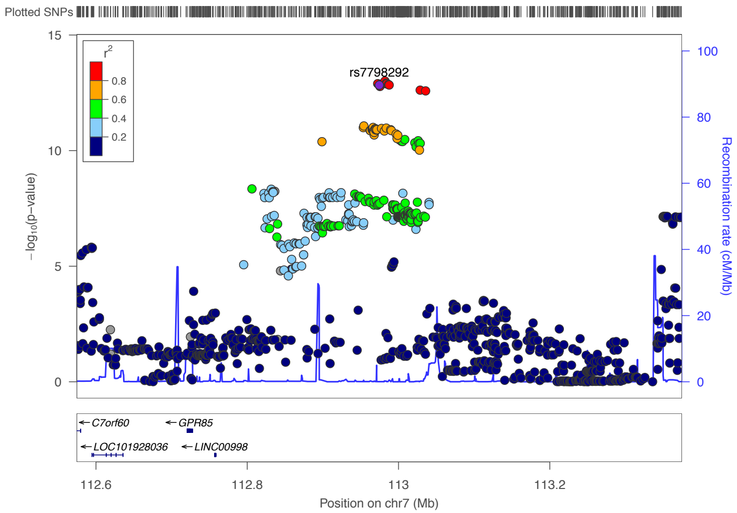


Total population

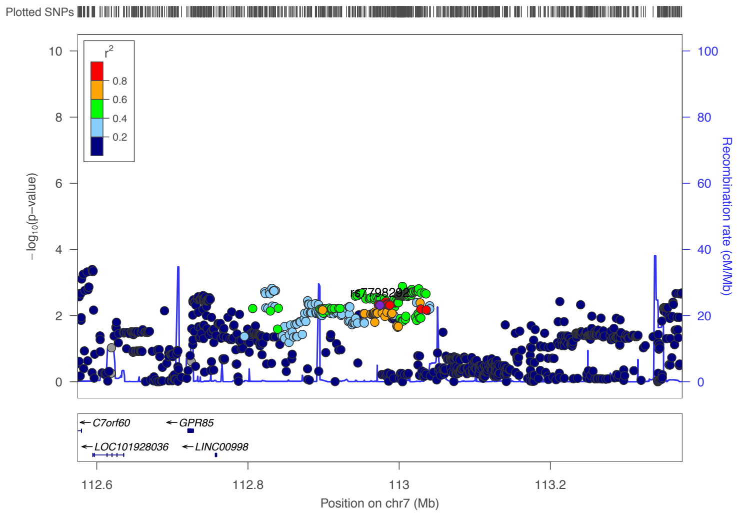


Short people

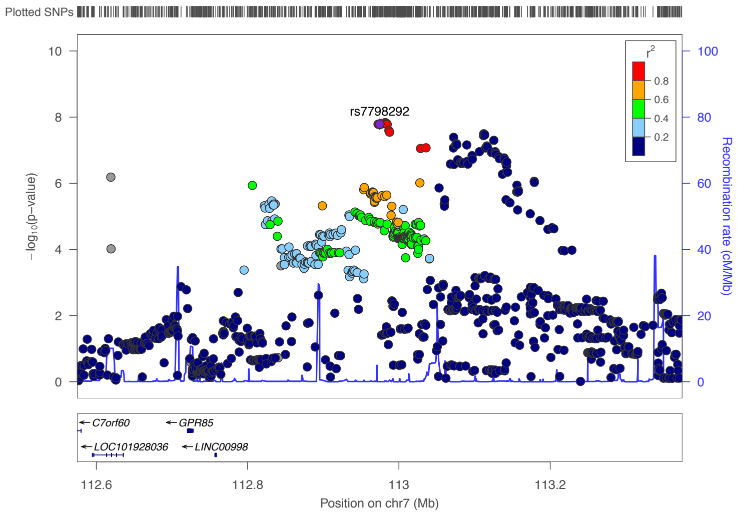


Tall people

### 7q31.1 (top SNP is rs12537134, a regulatory region variant in *LINC00998*).

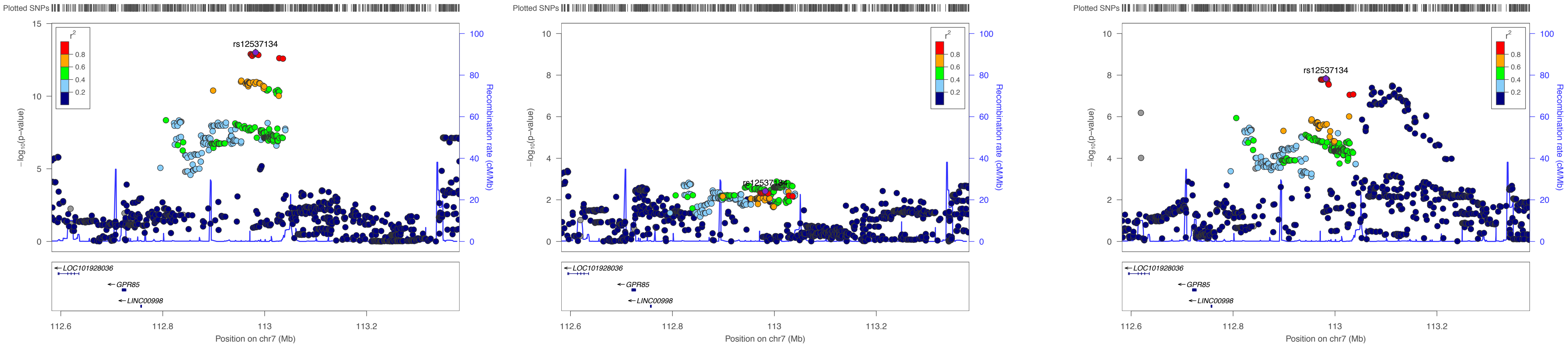

### 8p23.1 (Top SNP is rs3021500, an intron variant in *XKR6* and *MIR598*).

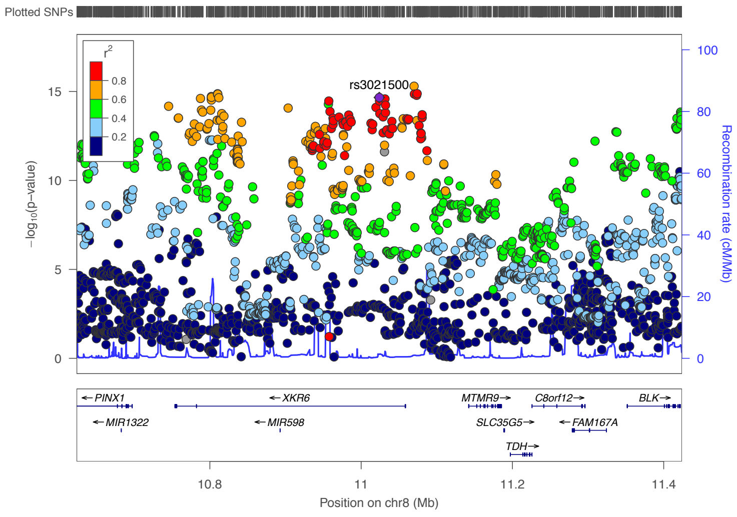


Total population

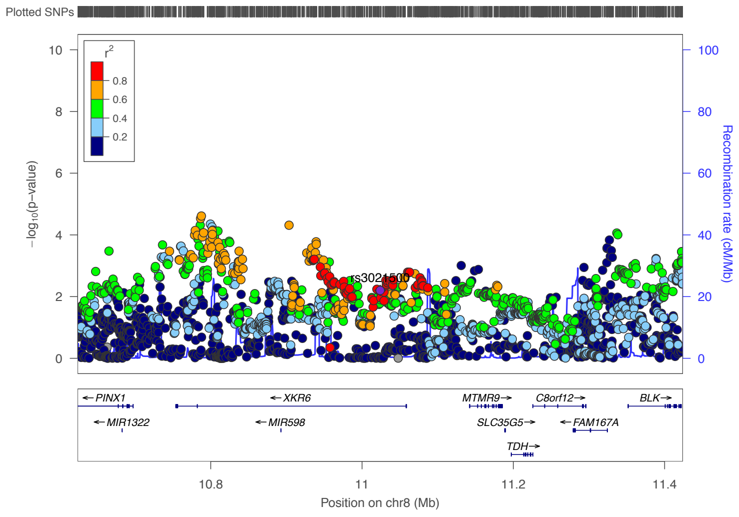


Short people

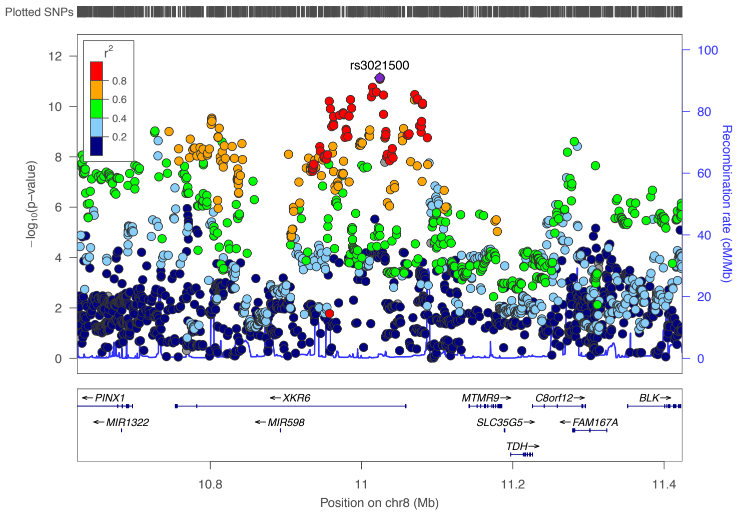


Tall people

### 9p22.3 (top SNP is rs10756686, an intron variant in *CCDC171*).

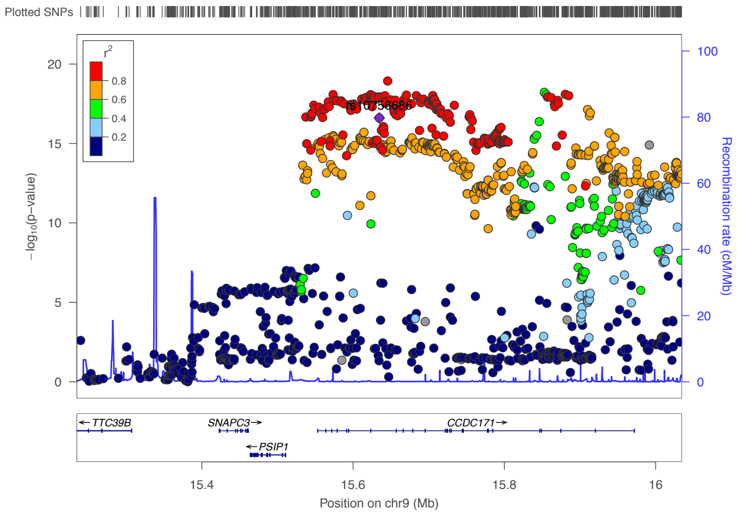


Total population

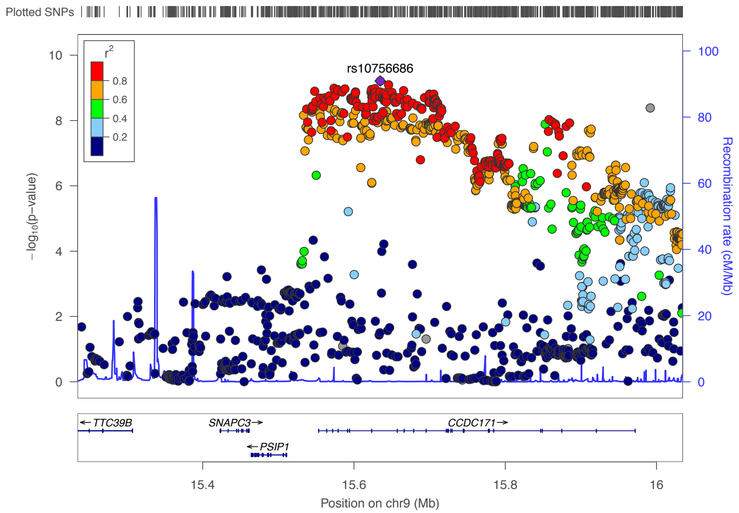


Short people

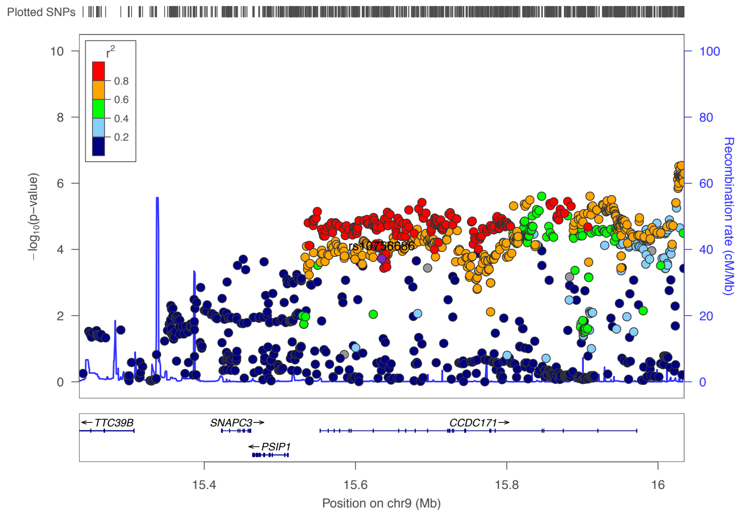


Tall people

### 9q33.2 (top SNP is rs6478538,an intron variant or *a* transcript variant in *TTLL11*).

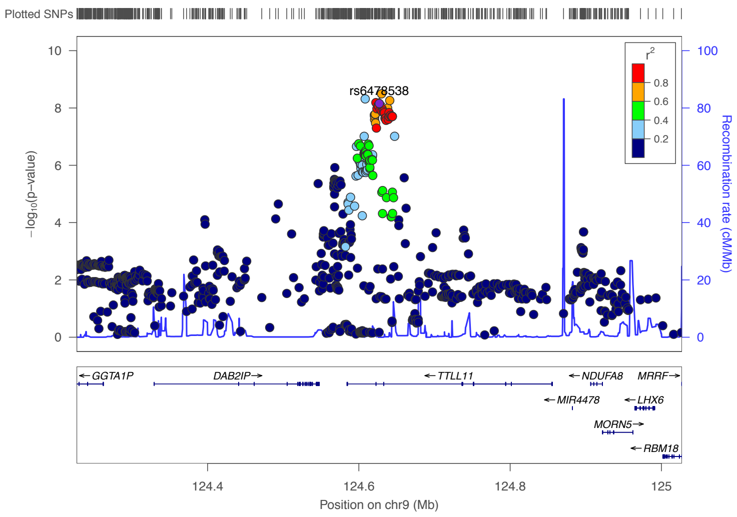


Total population

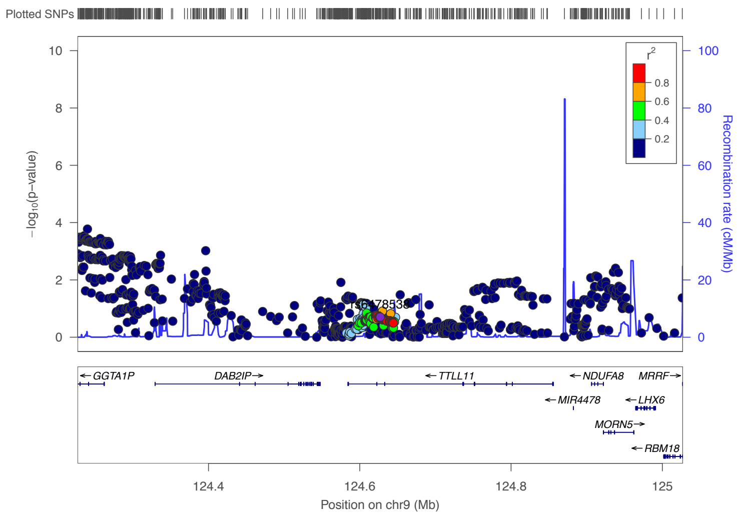


Short people

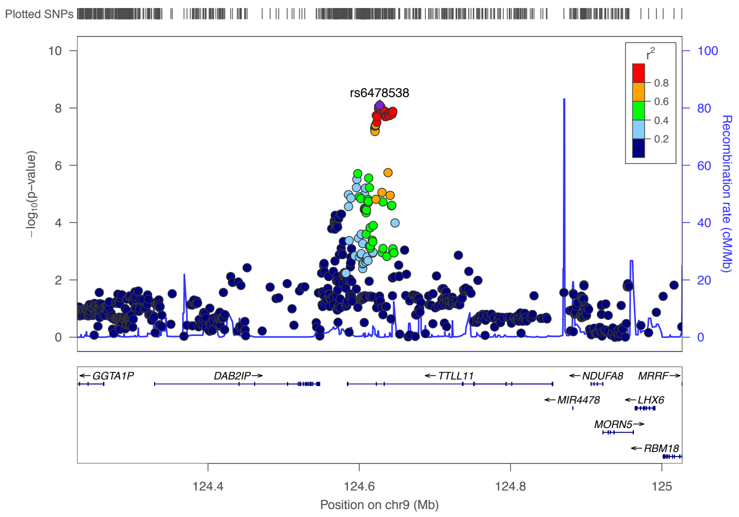


Tall people

### 11p14.1 (top SNP is rs80083564, an intron variant in *BDNF*).

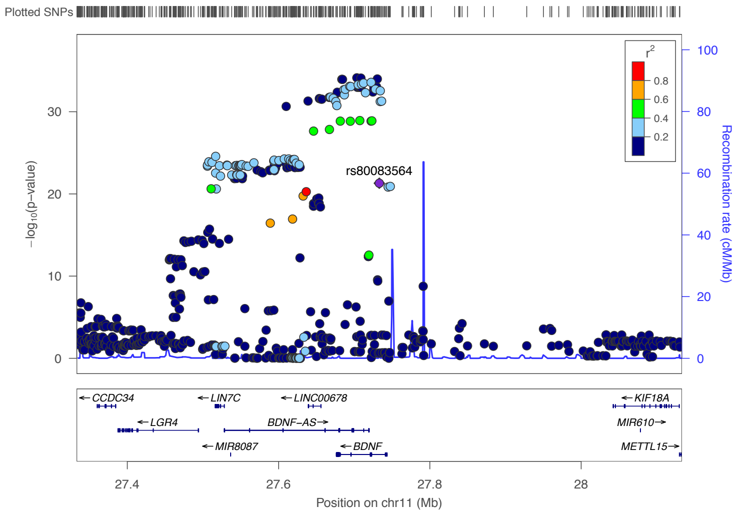


Total population

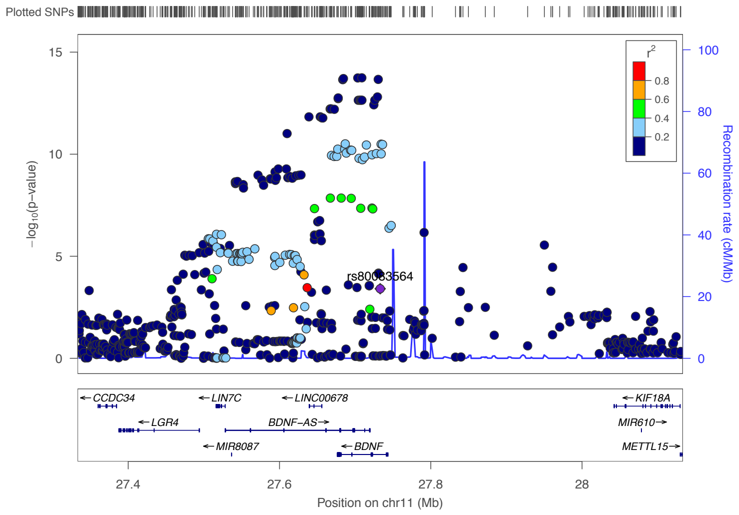


Short people

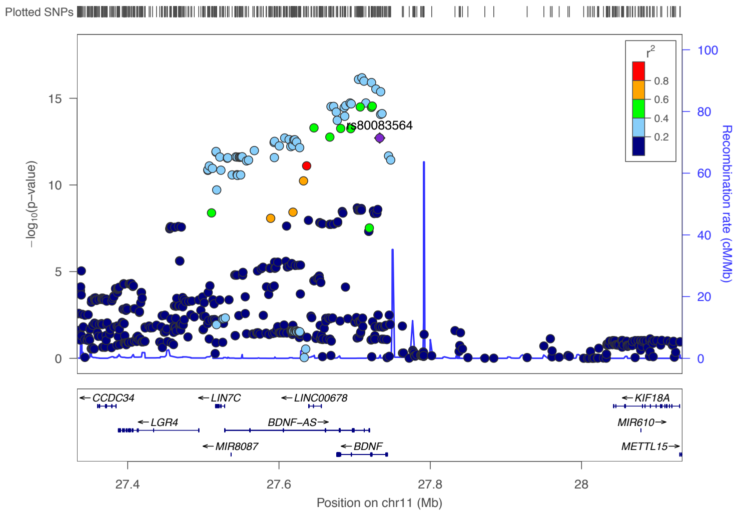


Tall people

### 11p11.2 (top SNP is rs7928842, an intron variant in *CELF1*).

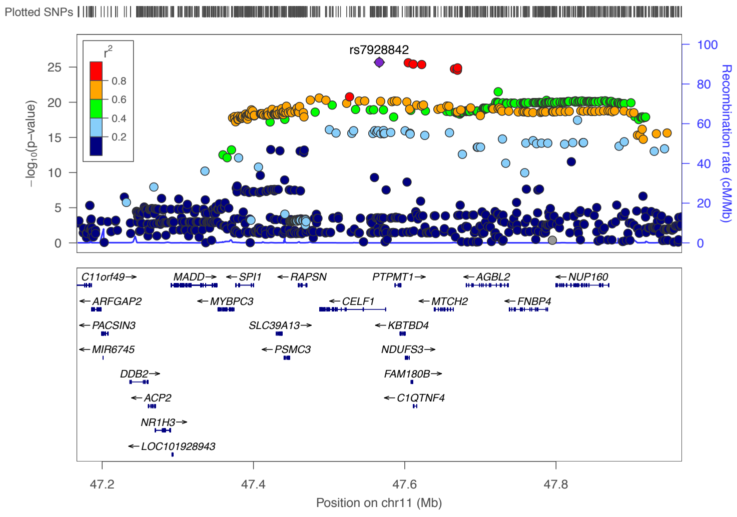


Total population

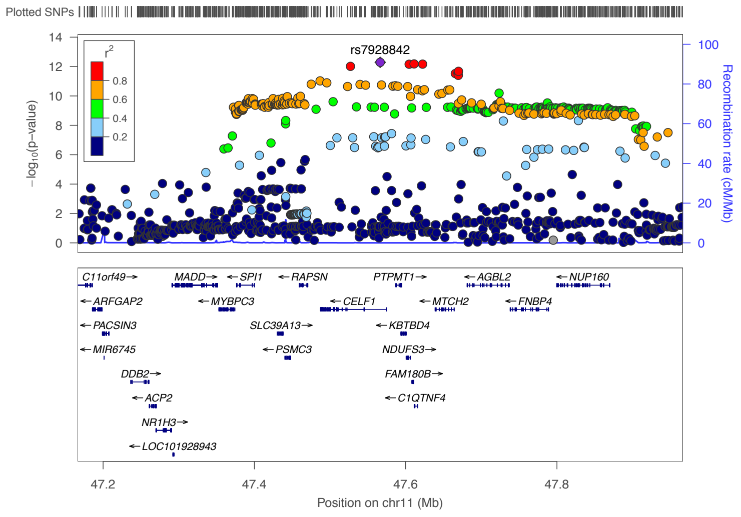


Short people

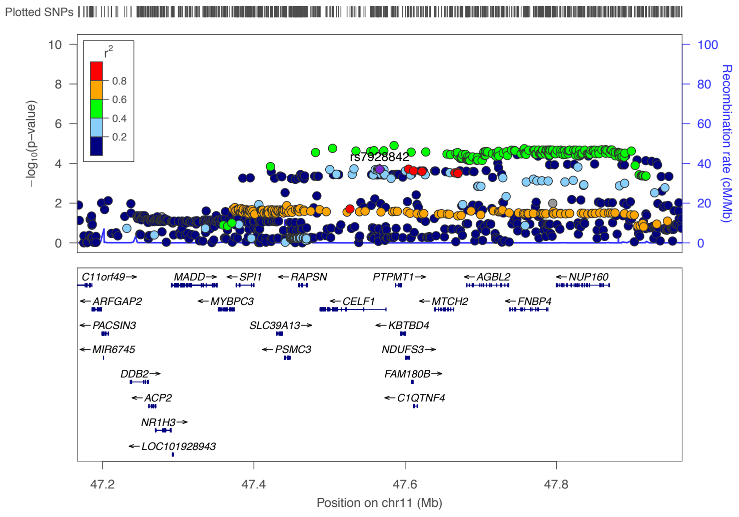


Tall people

### 11q13.1 (top SNP is rs11227317, an intron variant in *CFL1*).

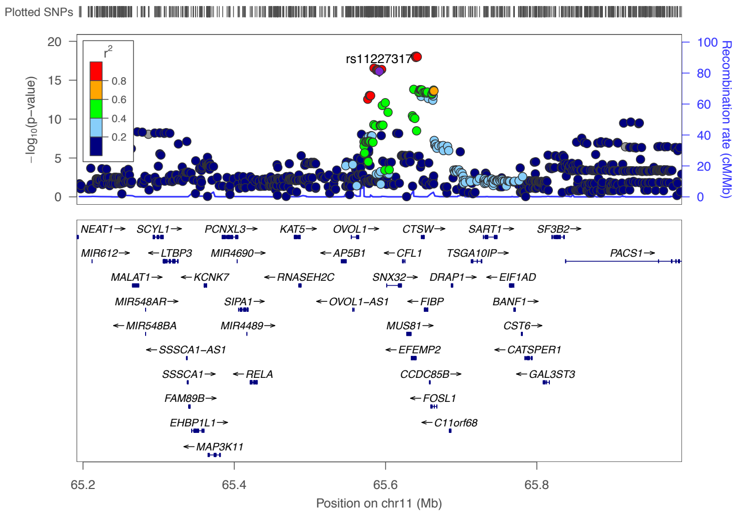


Total population

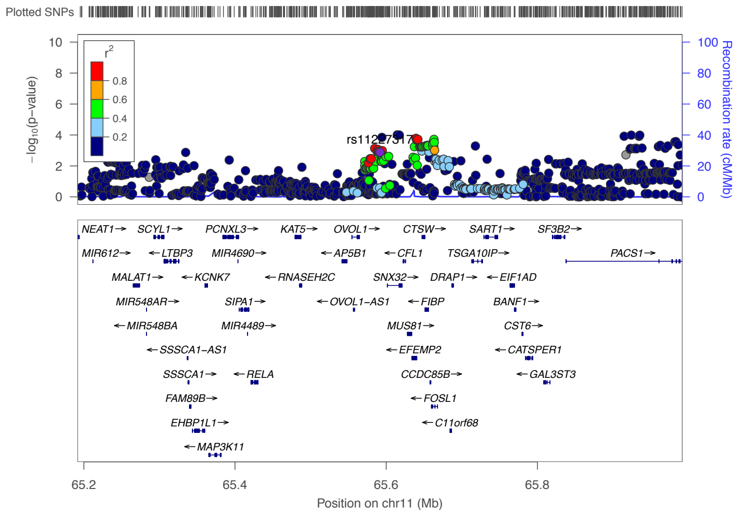


Short people

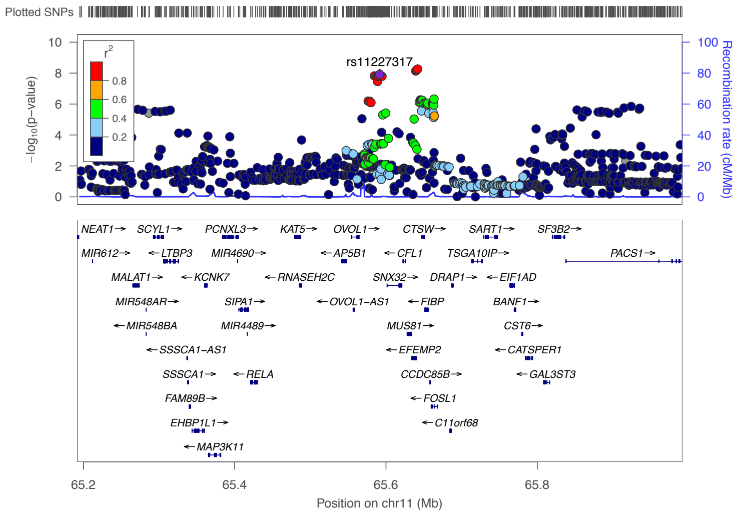


Tall people

### 11q25 (top SNP is rs329648, a down stream gene variant for *IGSF9B*).

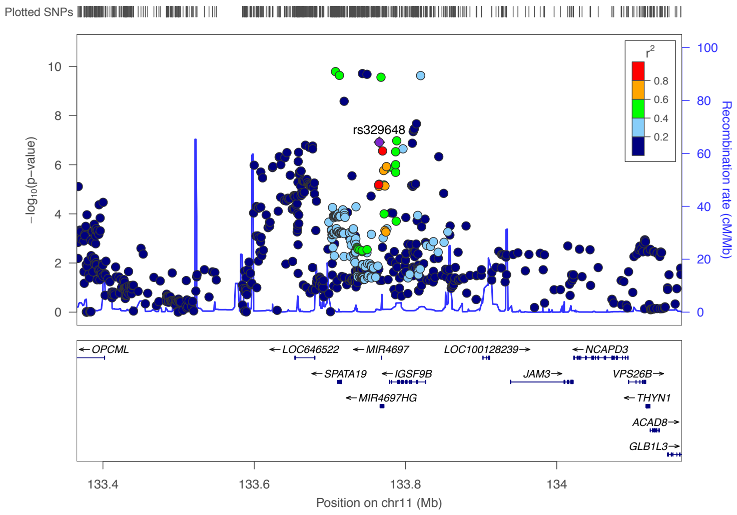


Total population

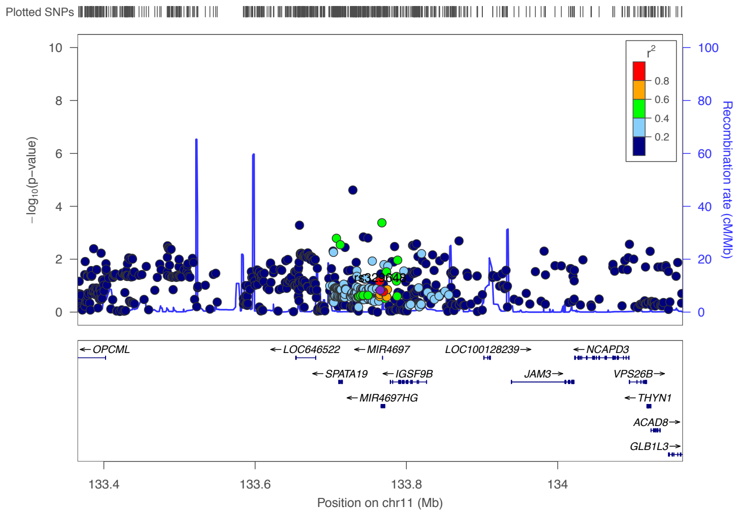


Short people

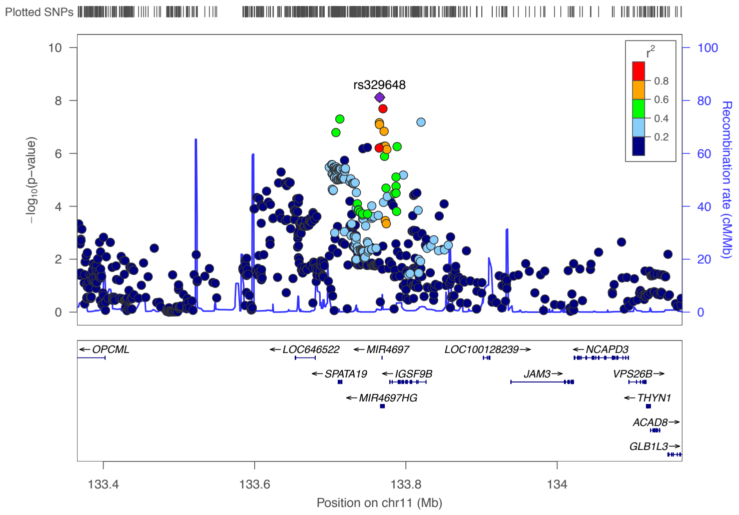


Tall people

### 12q24.31(top SNP is rs147730268, an intron variant in *KNTC1*).

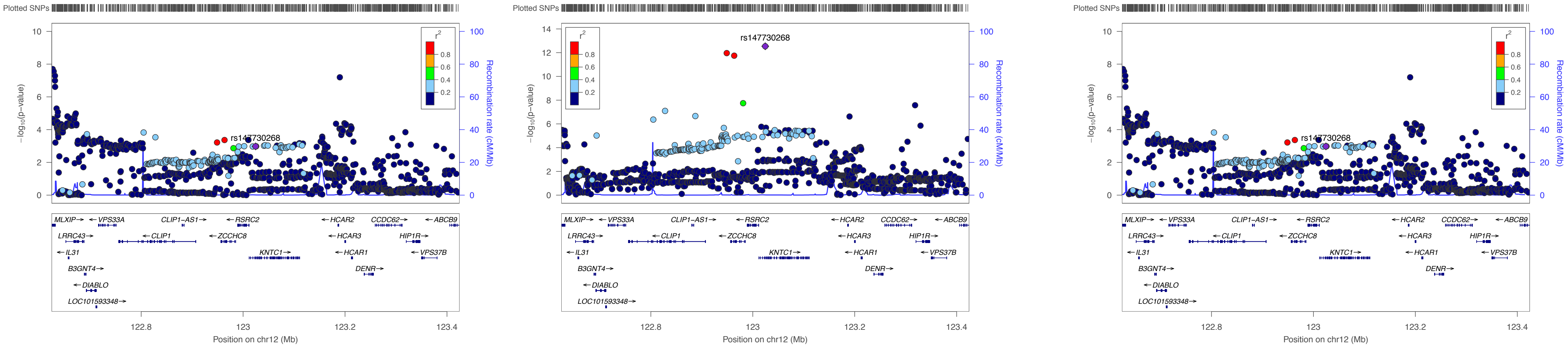


Total population

Short people

Tall people

### 13q14.3 (top SNP is rs1334874, an intergenic variant nearby *LINC00458*).

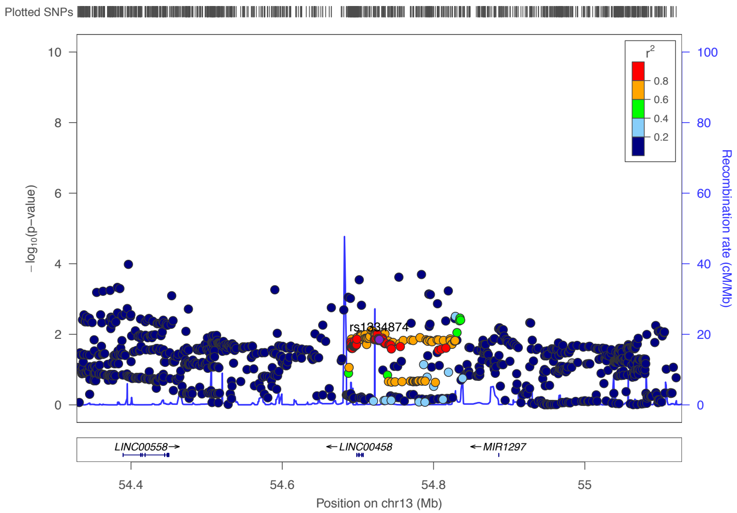


Total population

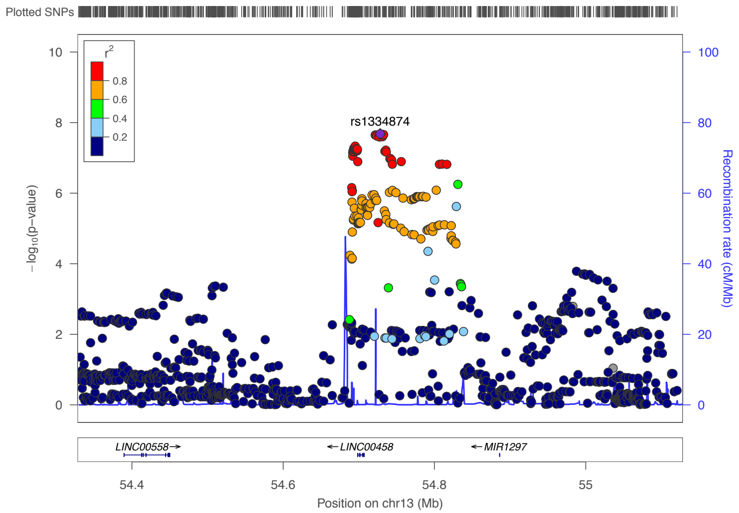


Short people

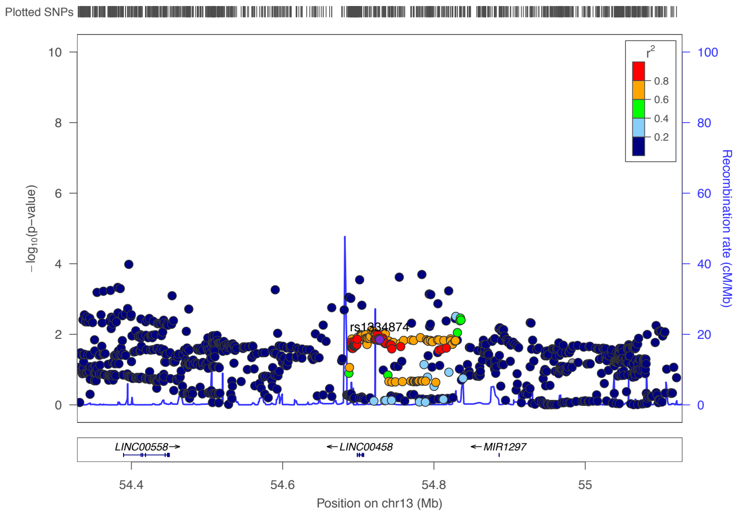


Tall people

### 13q31.1 (top SNP is rs1441262, an intergenic variant in *LINC00331*).

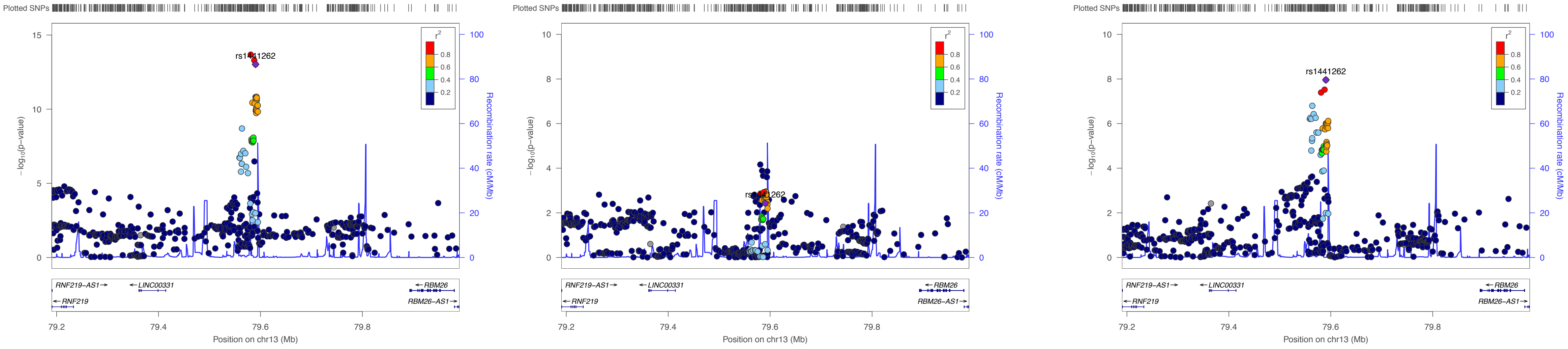


Total population

Short people

Tall people

### 16p13.3 (top SNP is rs2238435, an 3’ UTR variant for *ADCY9*).

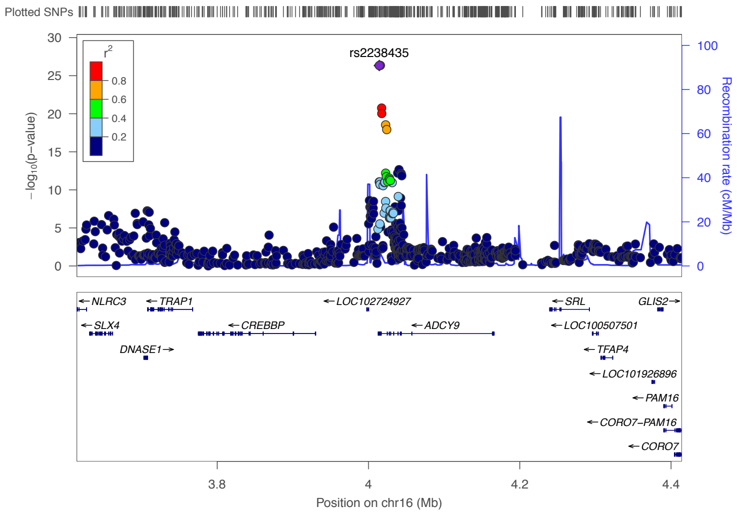


Total population

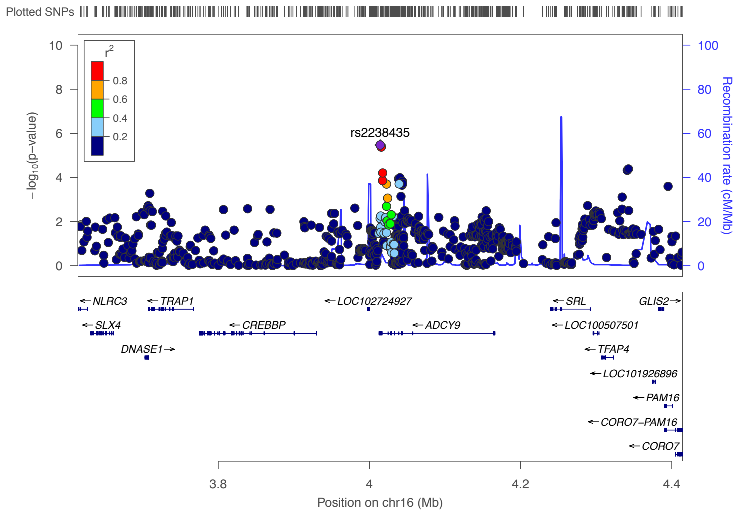


Short people

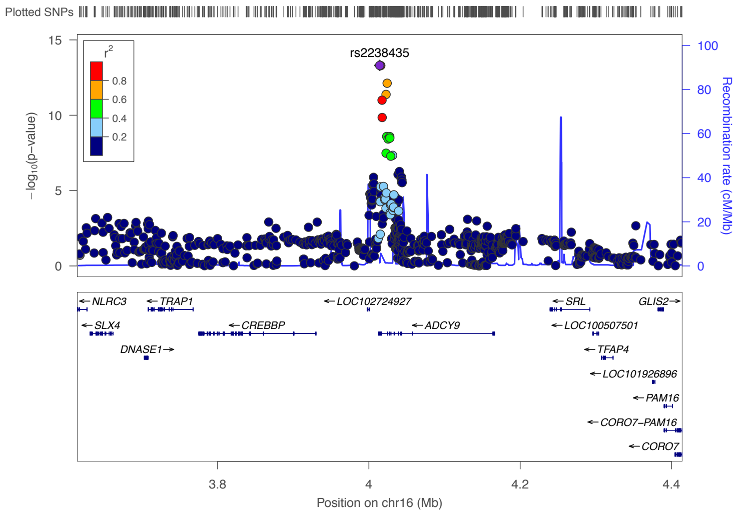


Tall people

### 16p13.3 (top SNP is rs34033929, an intron variant for *ADCY9*).

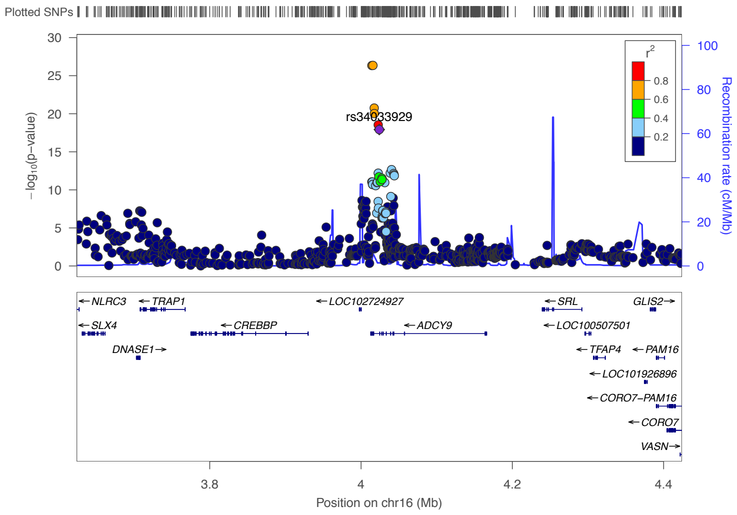


Total population

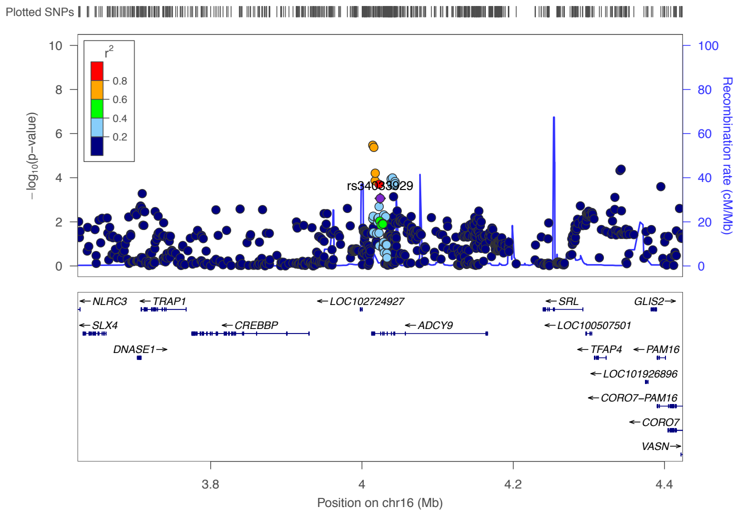


Short people

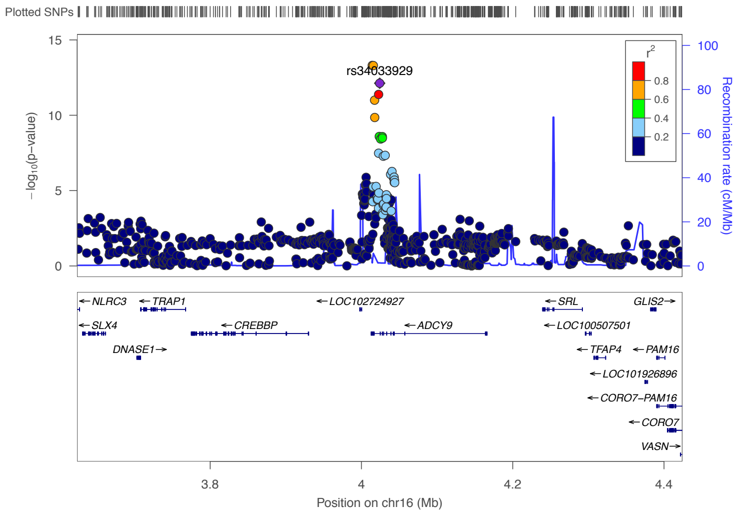


Tall people

### 16p12.3 (top SNP is rs11647854, an intron variant in *PDILT*).

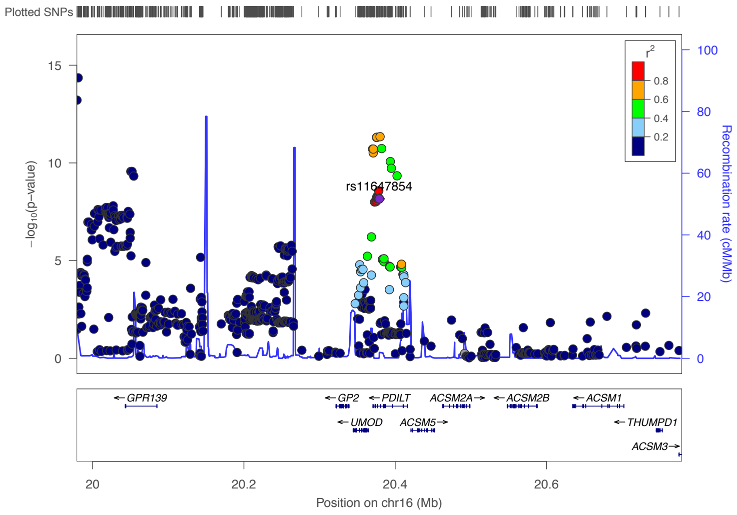


Total population

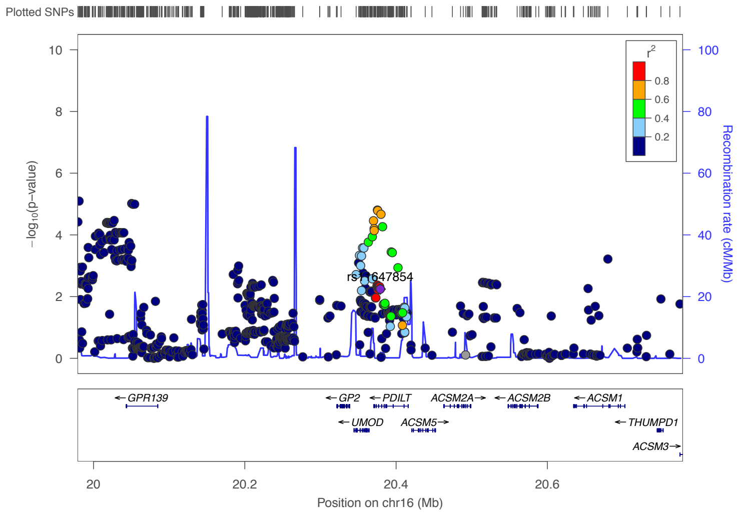


Short people

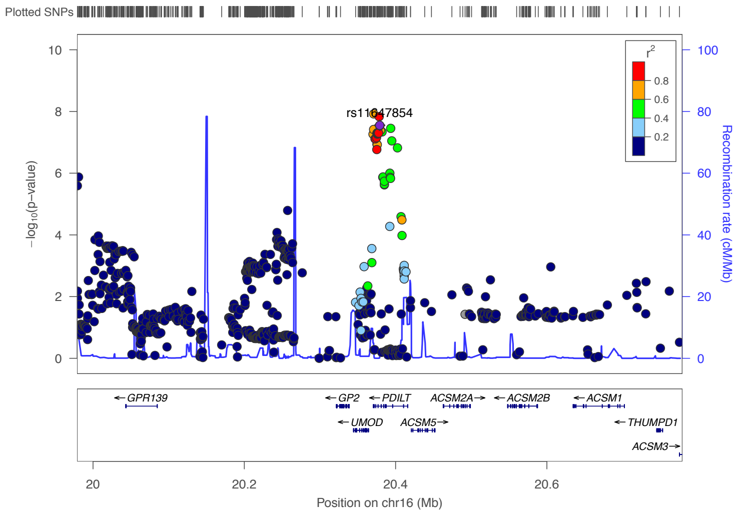


Tall people

### 16p11.2 (top SNP is rs2925624, an non coding transcript exon variant *SULT1A1*).

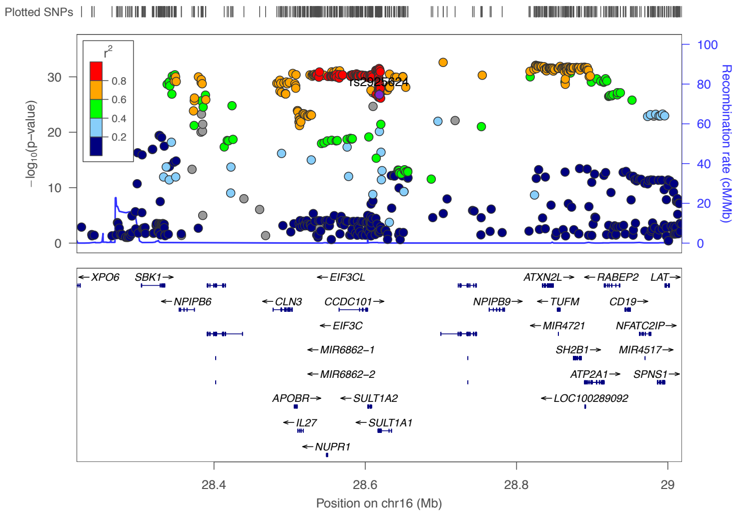


Total population

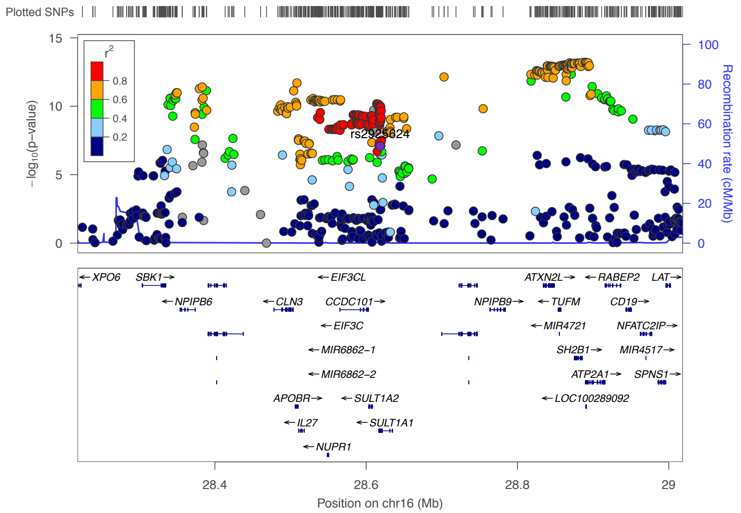


Short people

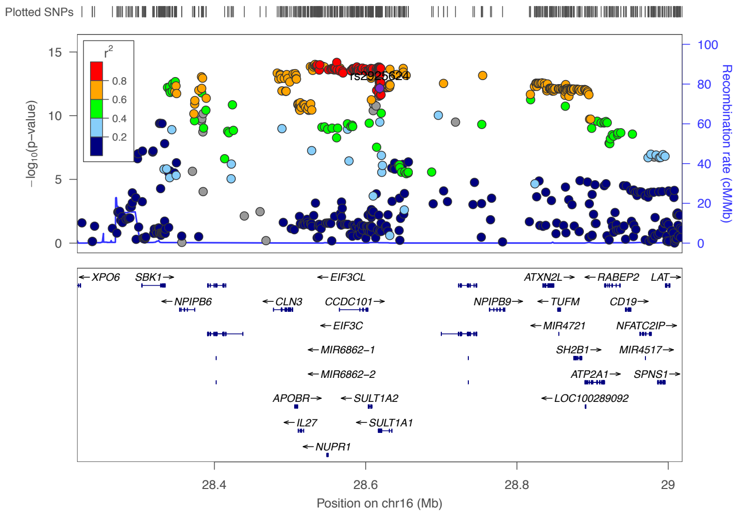


Tall people

### 16p11.2 (top SNP is rs375452507, an upstream gene variant for *BCKDK*).

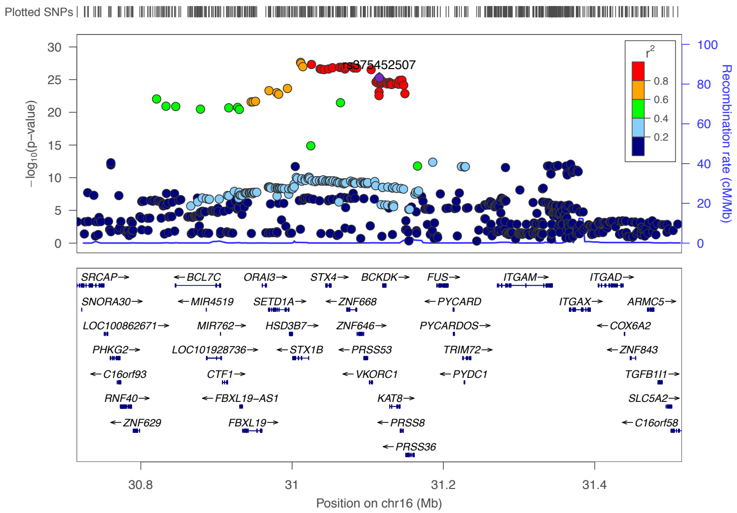


Total population

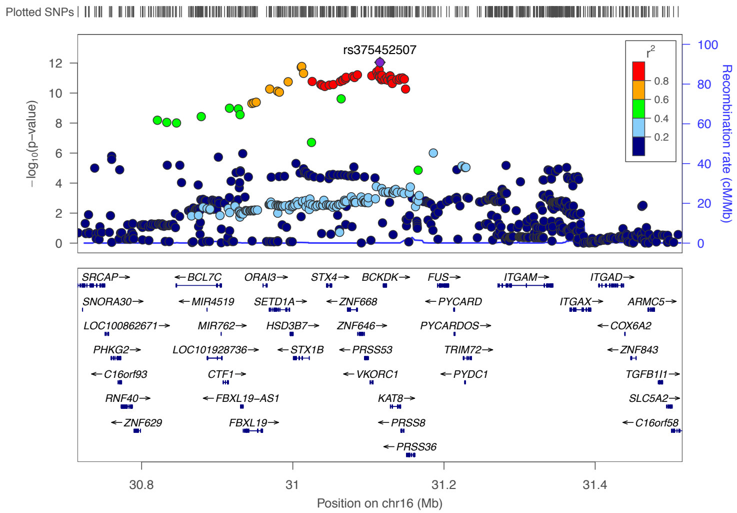


Short people

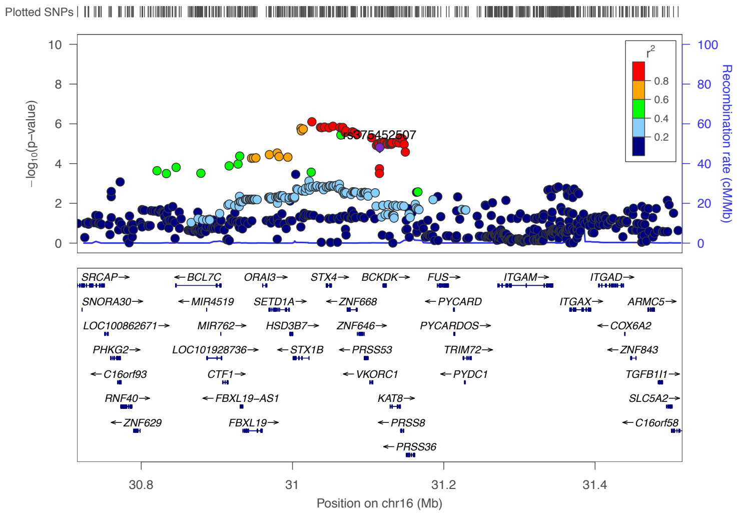


Tall people

### 16q12.2(top SNP is rs8055259, an intron variant  for *FTO*).

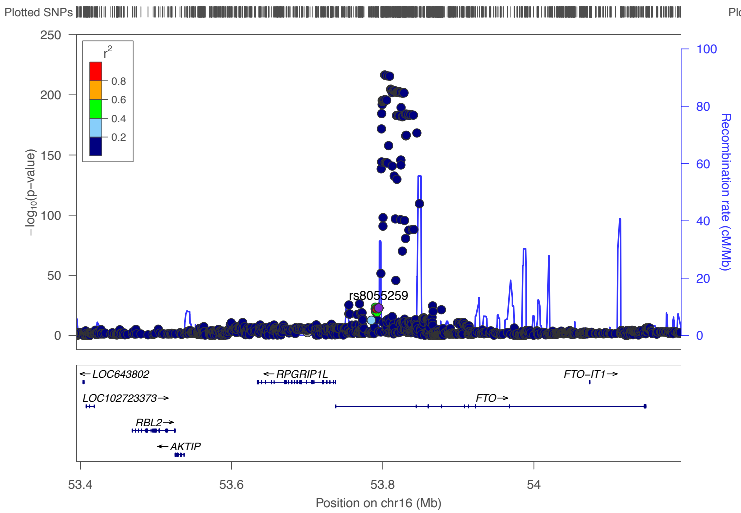


Total population

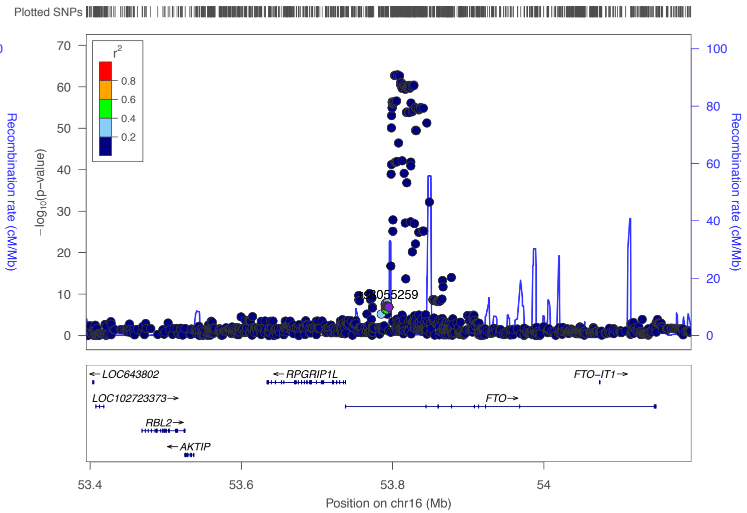


Short people

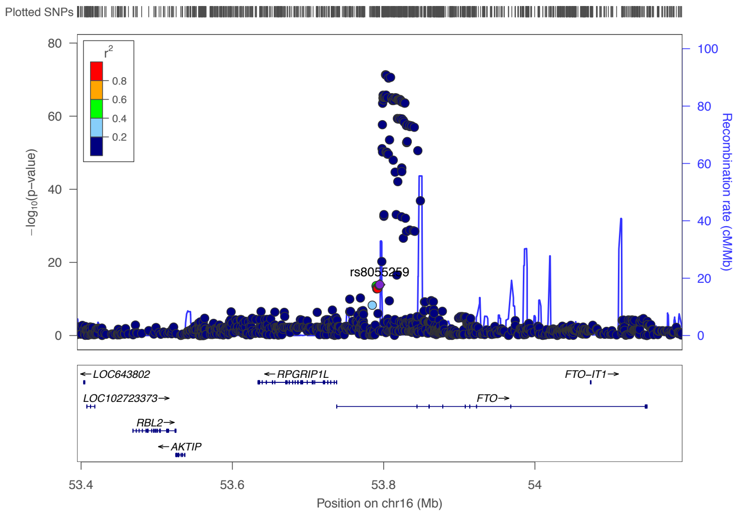


Tall people

### 16q12.2(top SNP is rs8055253, an intron variant for *FTO*).

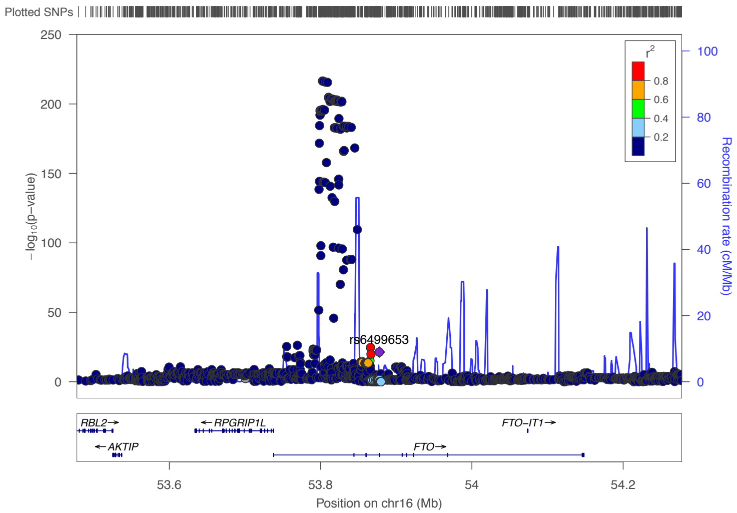


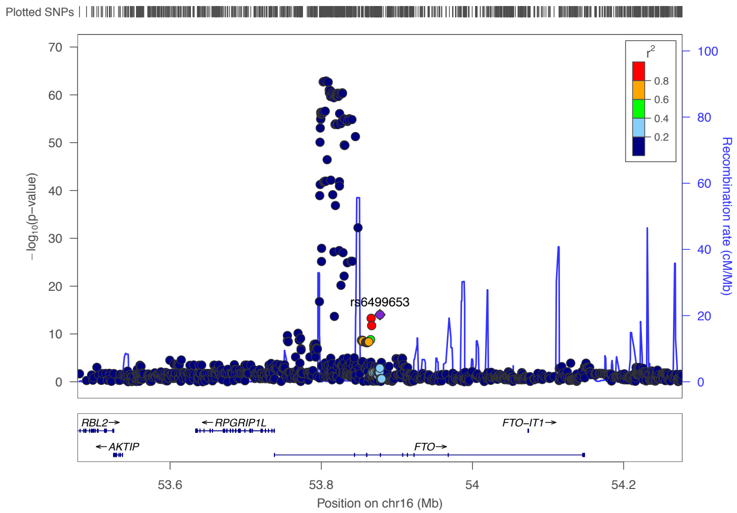


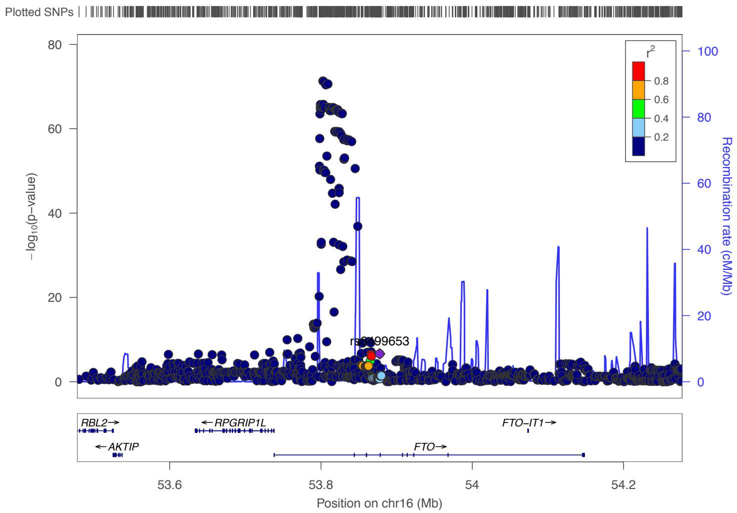


Total population

Short people

Tall people

### 17q21.32 (top SNP is rs68085814, a downstream gene variant *IGF2BP1*).

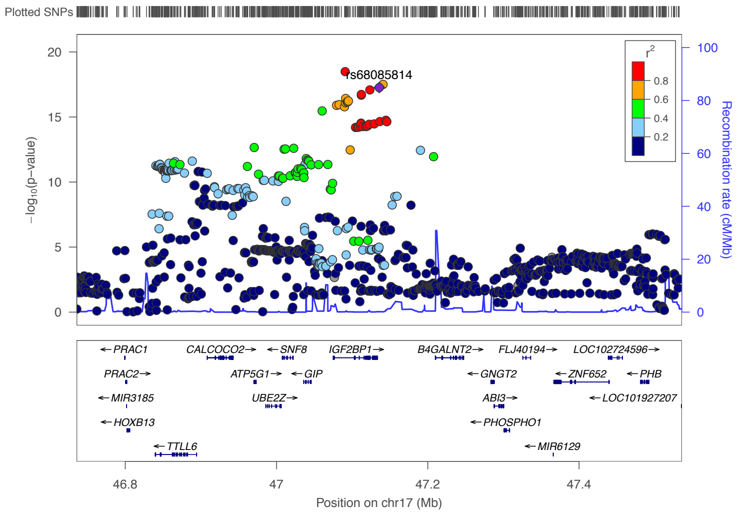


Total population

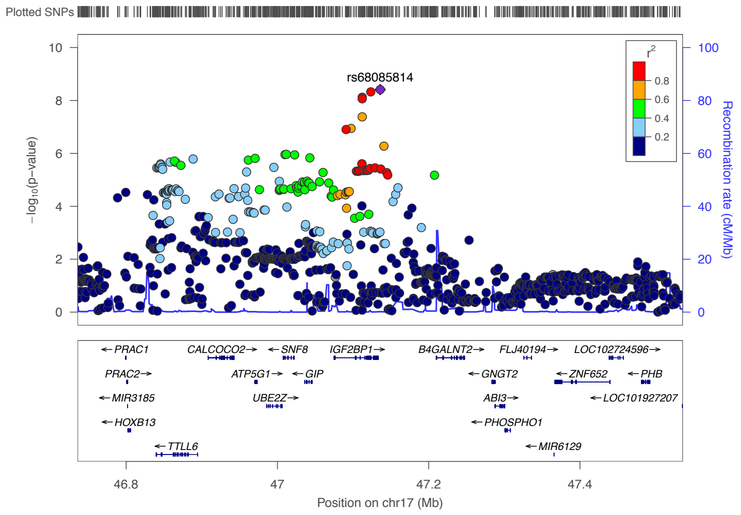


Short people

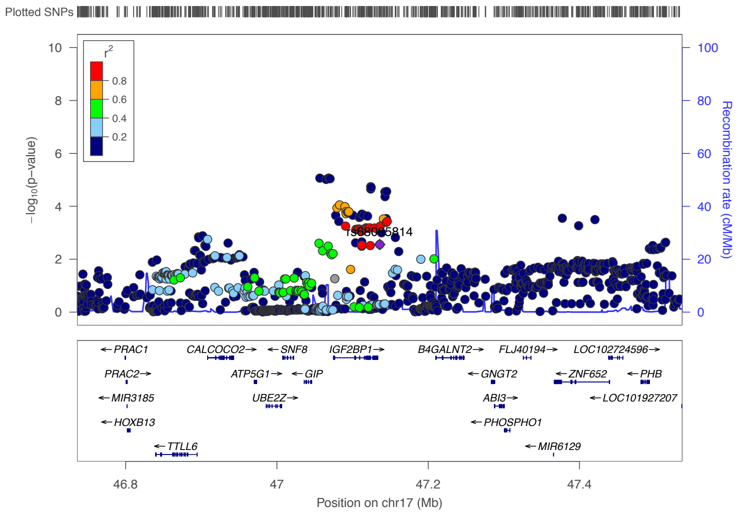


Tall people

### 17q25.3 (top SNP is rs901064, an intron variant in *RPTOR*).

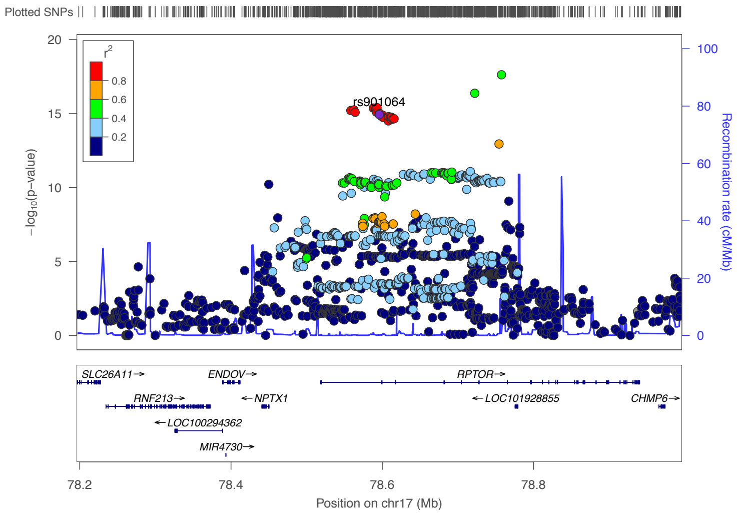


Total population

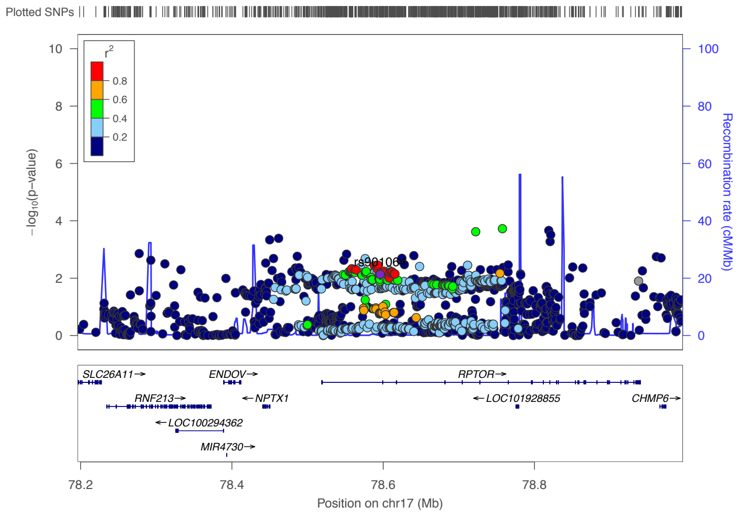


Short people

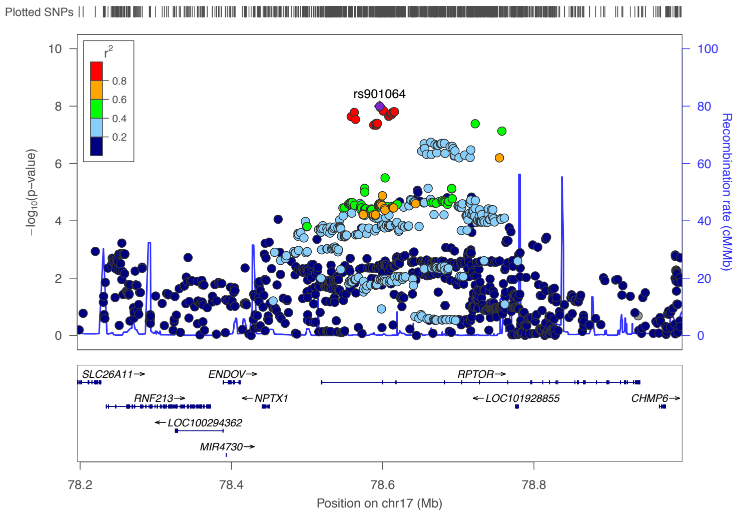


Tall people

### 18p11.31 (top SNP is rs73373997, a downstream gene variant in *EPB41L3*).

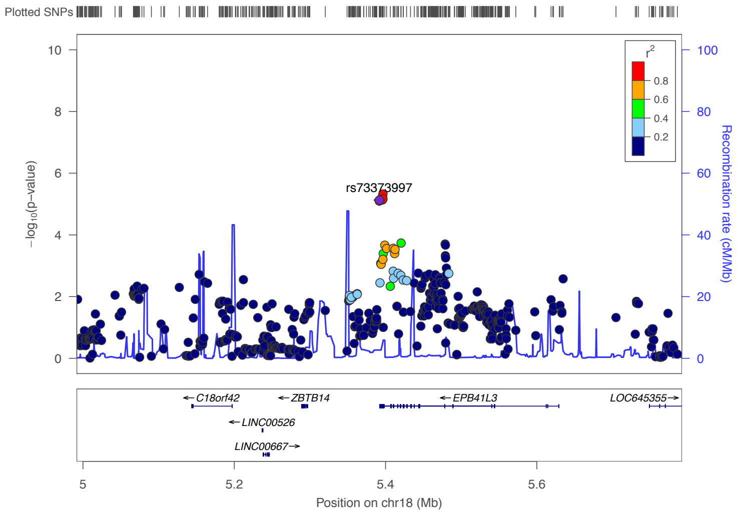


Total population

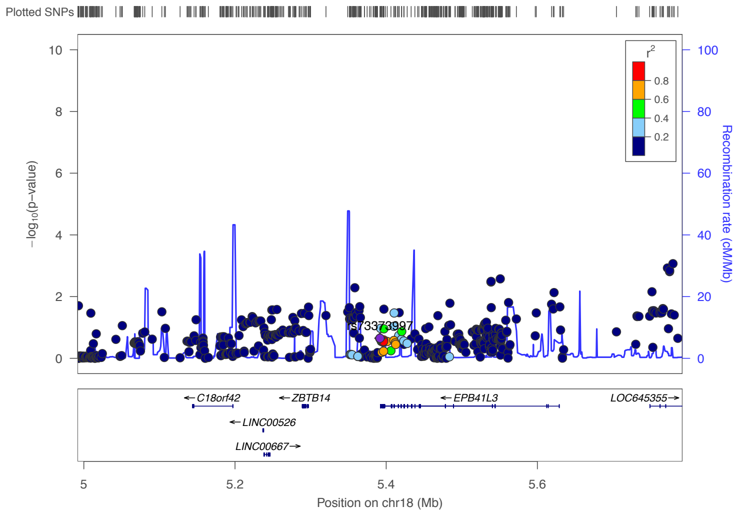


Short people

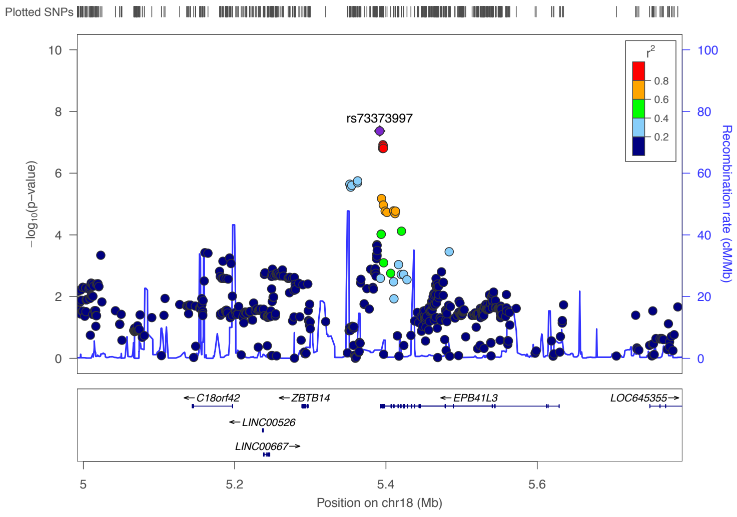


Tall people

### 18q12.3 (top SNP is rs1834145, an intergenic variant in *RIT2* and *SYT4*).

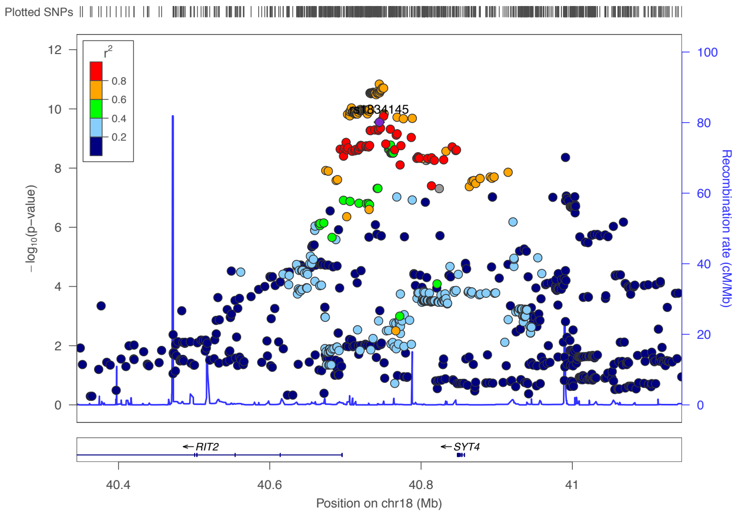


Total population

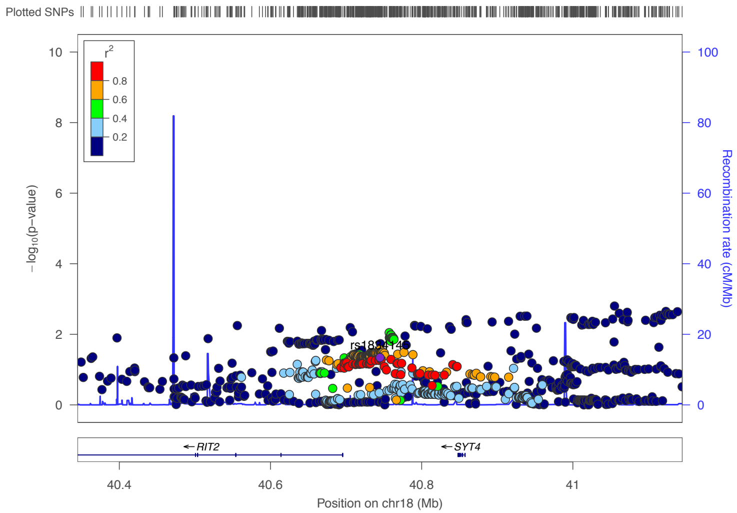


Short people

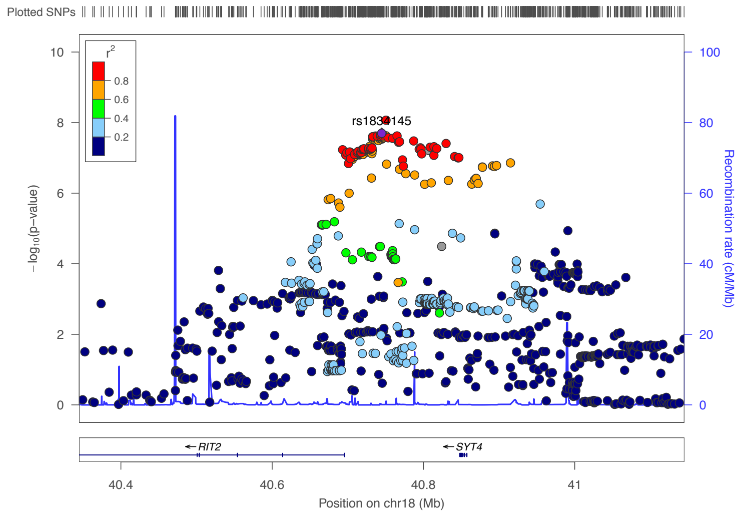


Tall people

### 18q21.32 (top SNP is rs2045442, a regulatory region variant in promoter flanking region of *MC4R*).

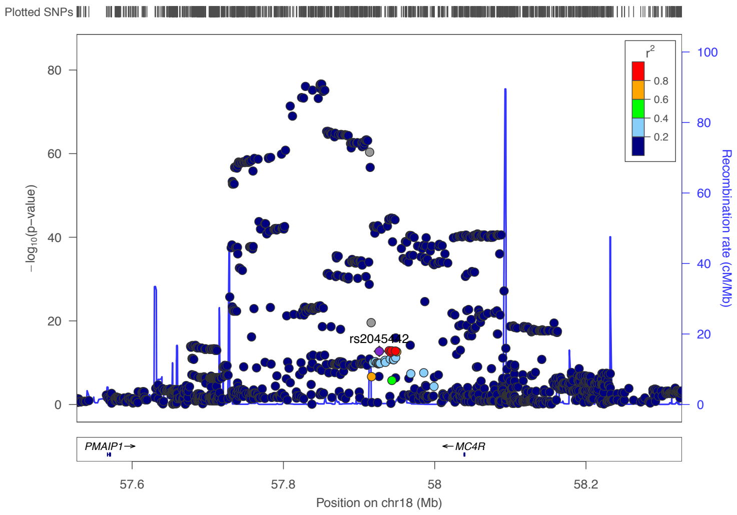


Total population

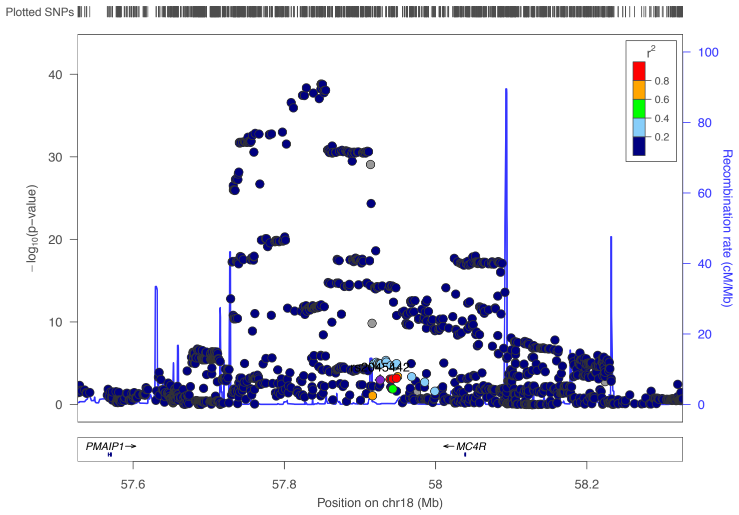


Short people

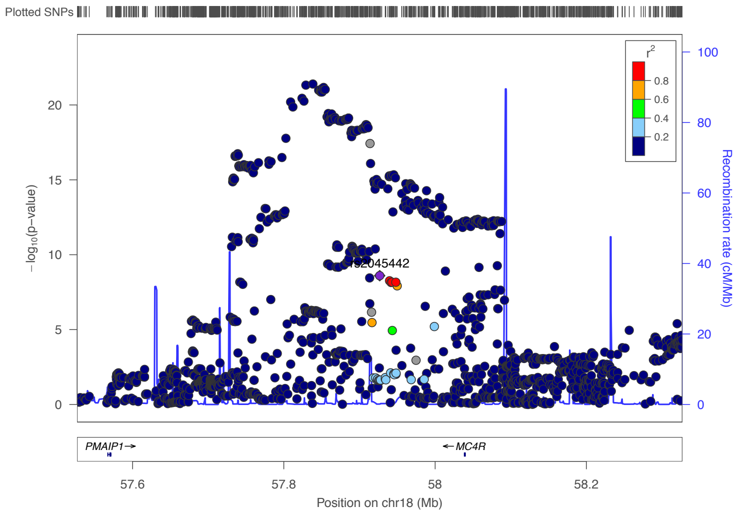


Tall people

### 18q21.32 (top SNP is rs80116344, a intergenic variant nearby *MC4R*).

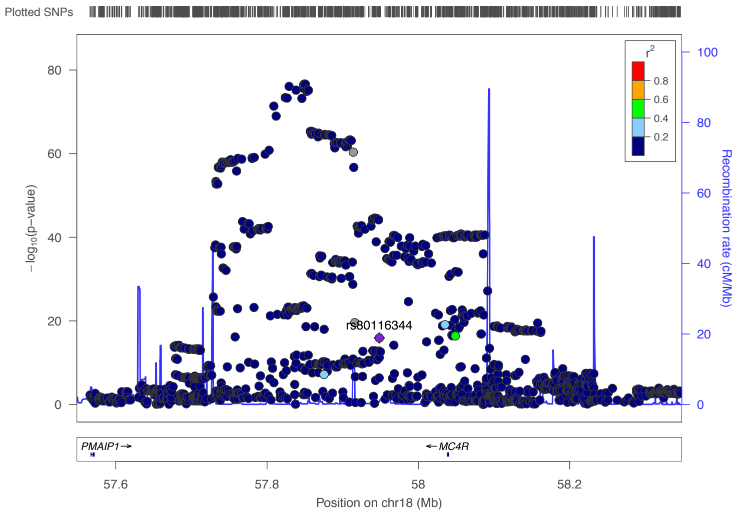


Total population

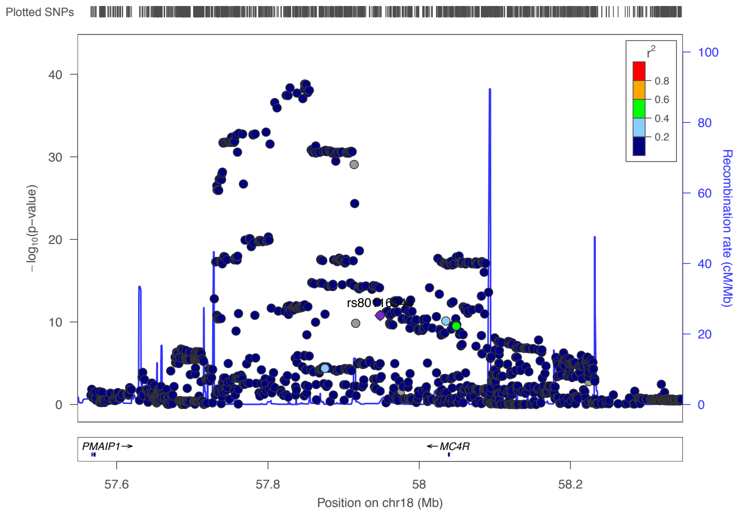


Short people

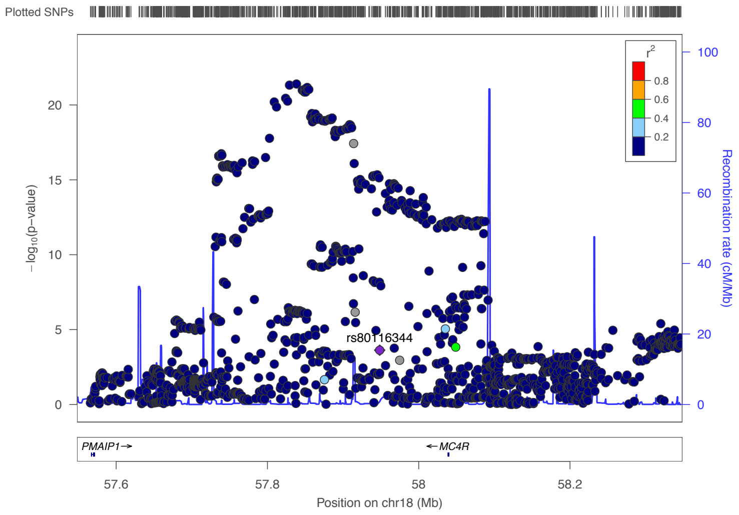


Tall people

### 18q21.32 (top SNP is rs9947450, a intergenic variant nearby *MC4R*).

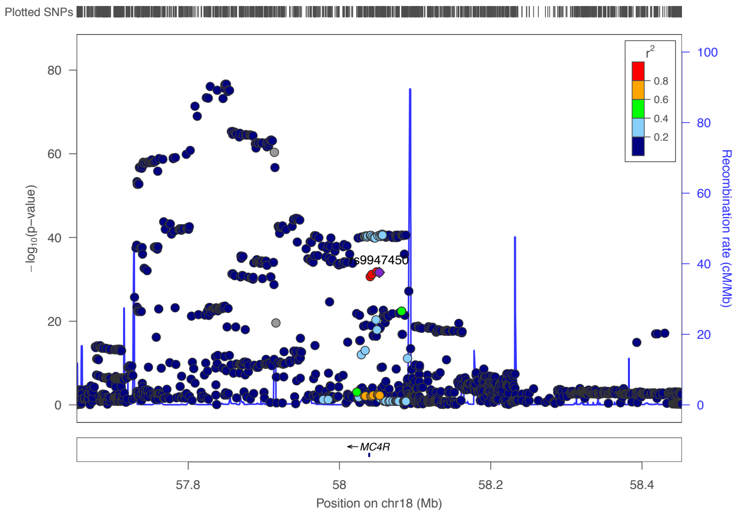


Total population

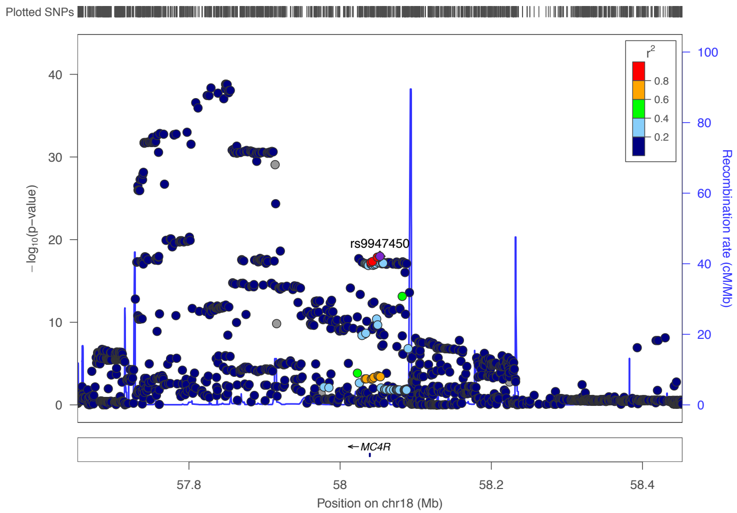


Short people

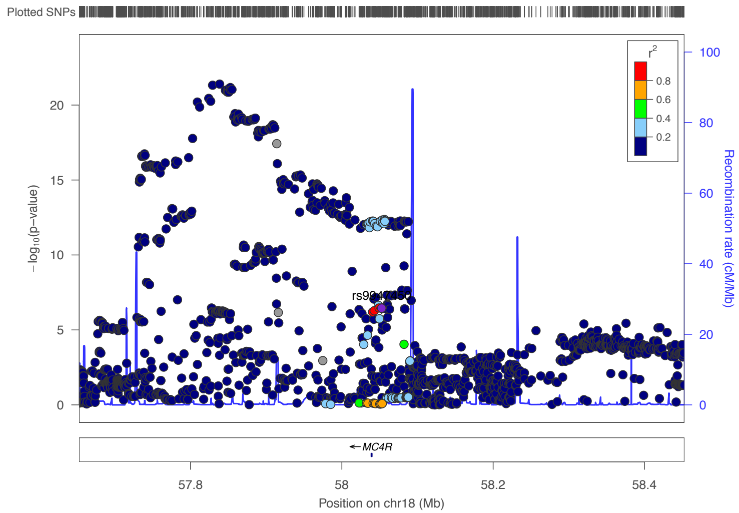


Tall people

### 18q21.32 (top SNP is rs6567168, an intron variant in *AC091576.1*).

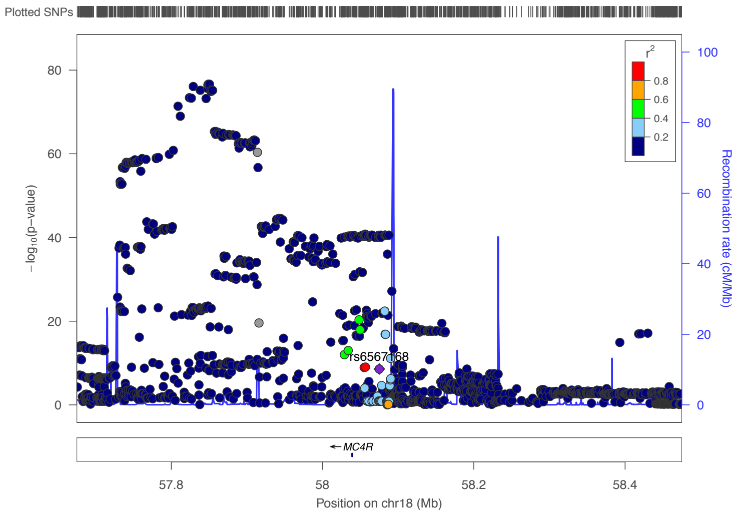


Total population

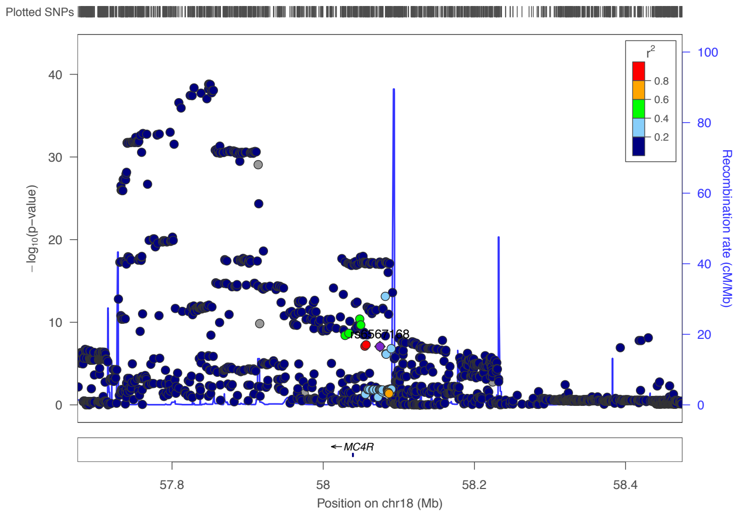


Short people

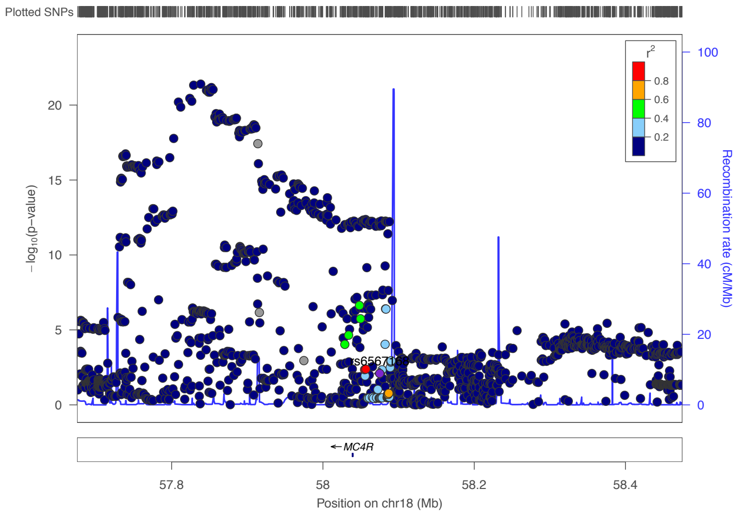


Tall people

### 19q13.32 (top SNP is rs10403089, an intron variant in *SAE1*).

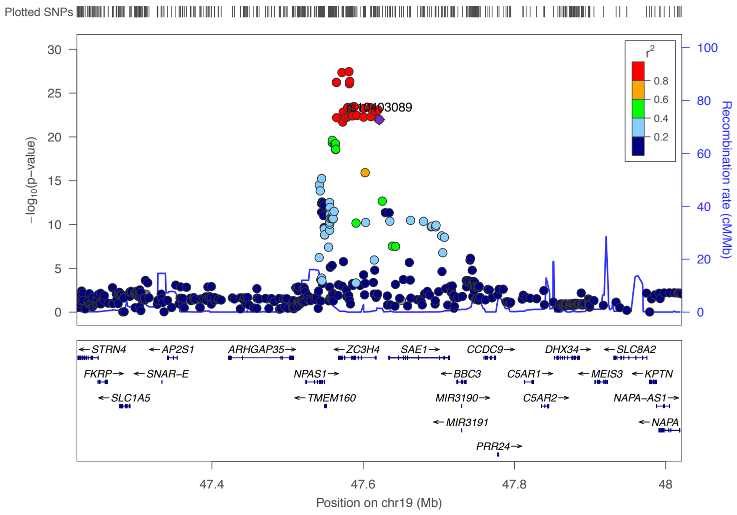


Total population

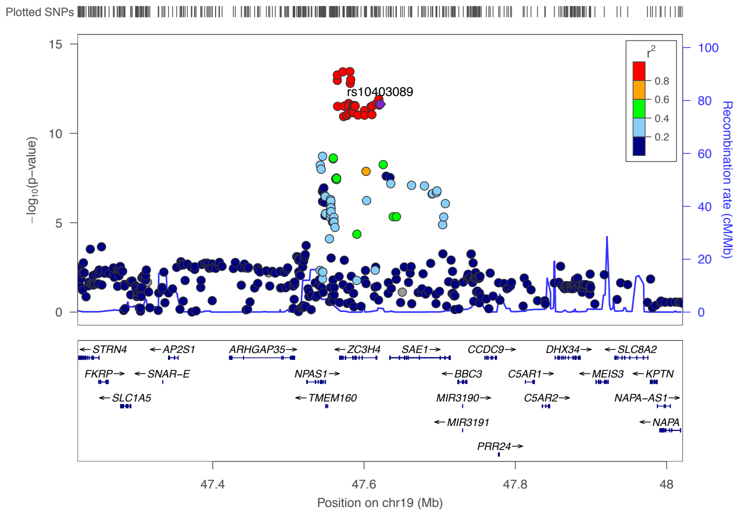


Short people

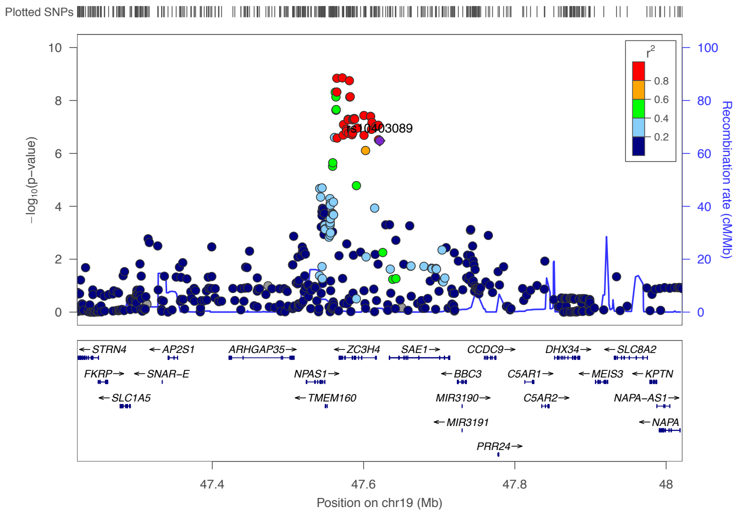


Tall people

### 20p12.2 (top SNP is rs6087022,an intron variant in *PAK5*).

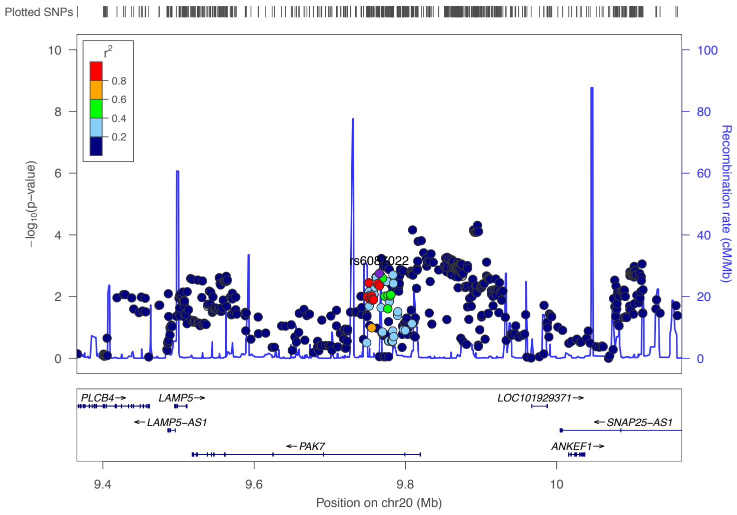


Total population

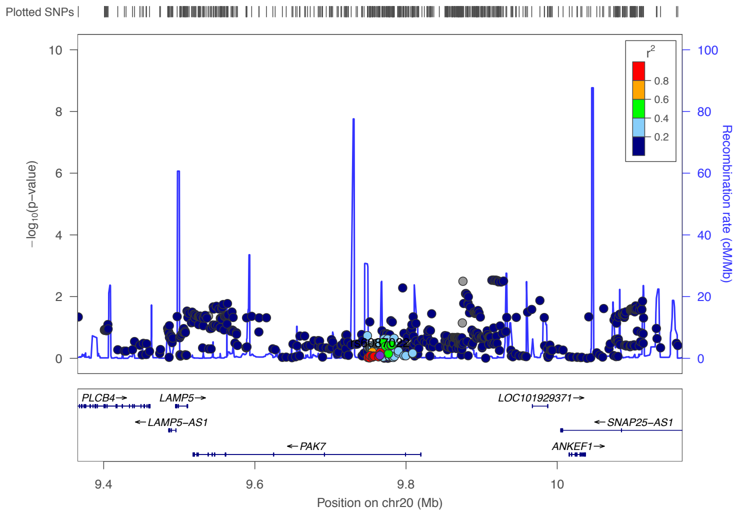


Short people

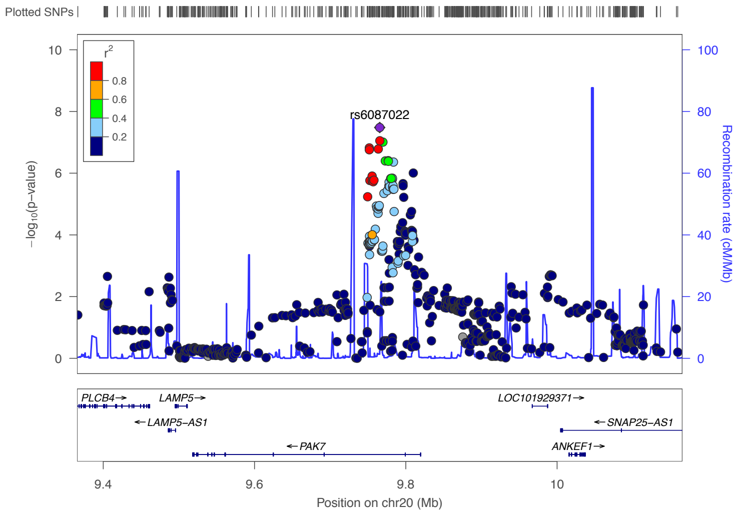


Tall people

### 20q13.2 (top SNP is rs6098902, an intergenic variant nearby *CBLN4*).

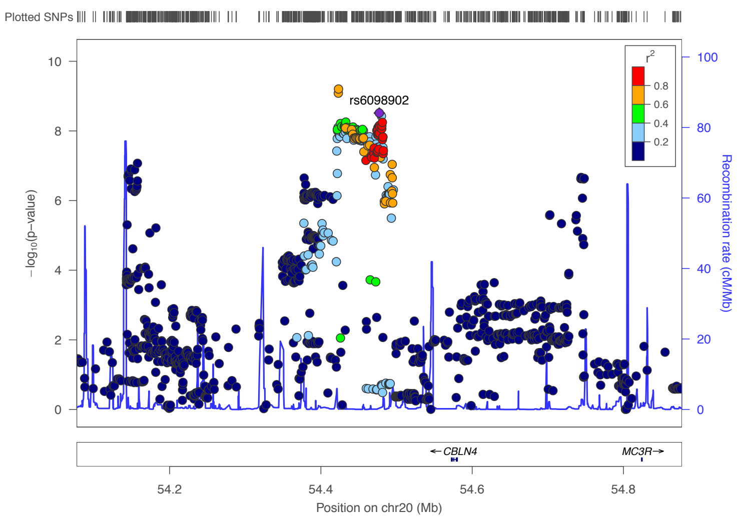


Total population

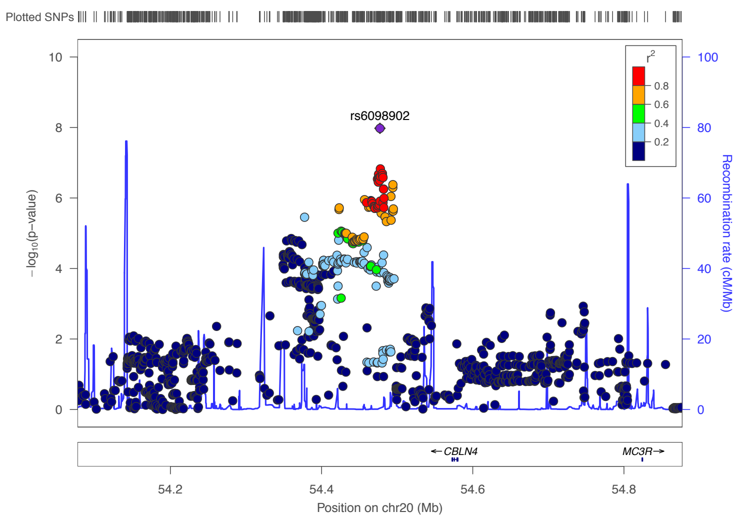


Short people

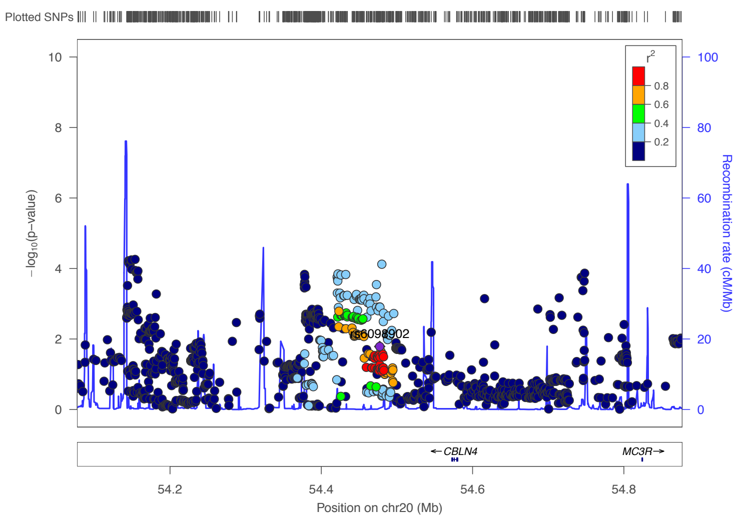


Tall people

### 21q22.3 (top SNP is rs9977825, an intron variant in *ADARB1*).

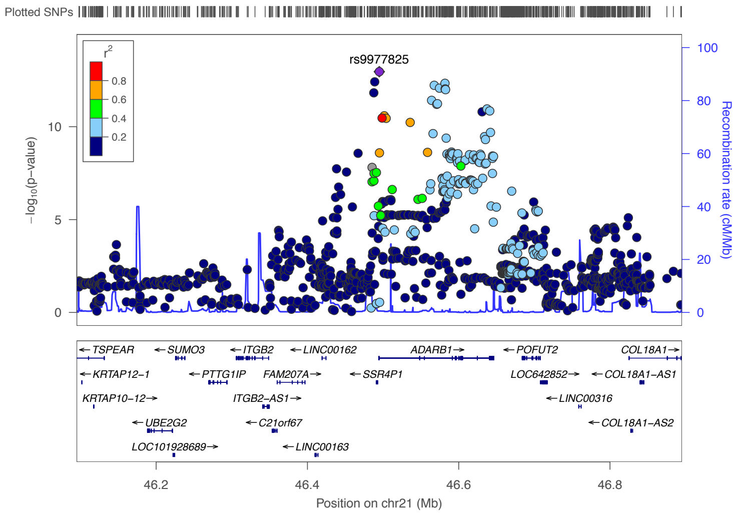


Total population

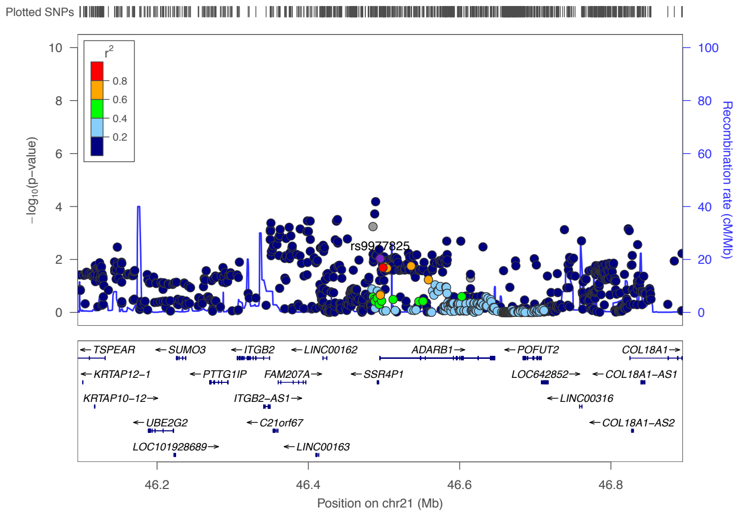


Short people

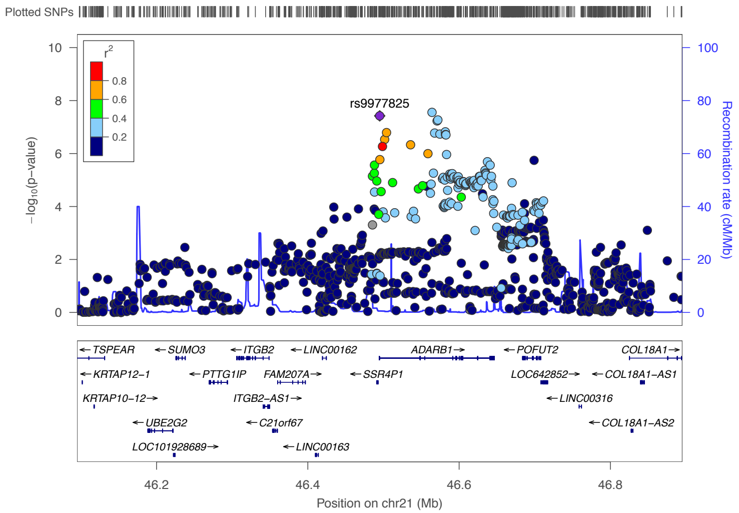


Tall people

### 22q13.1 (top SNP is rs4820409, an intron variant in *TNRC6B*).

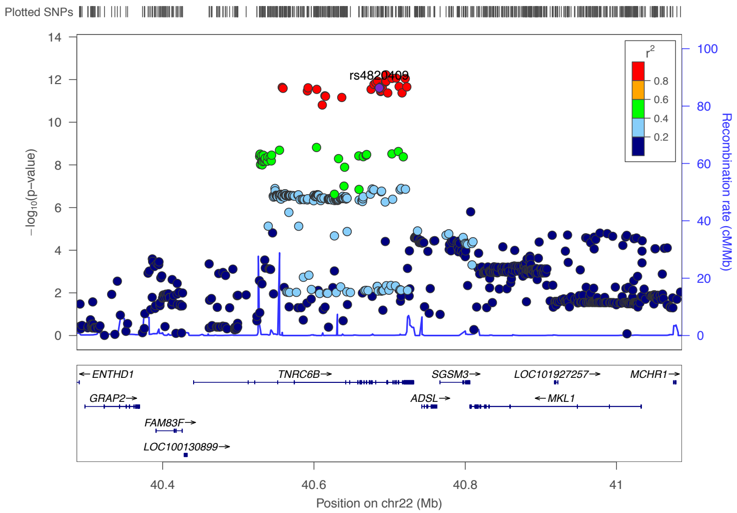


Total population

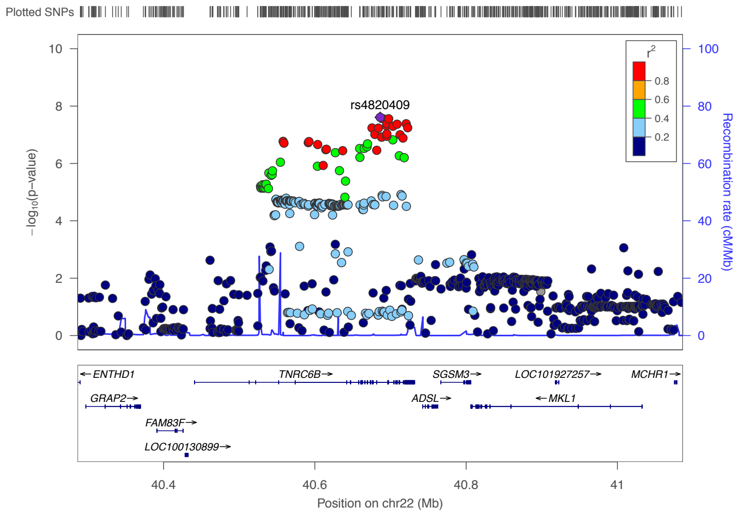


Short people

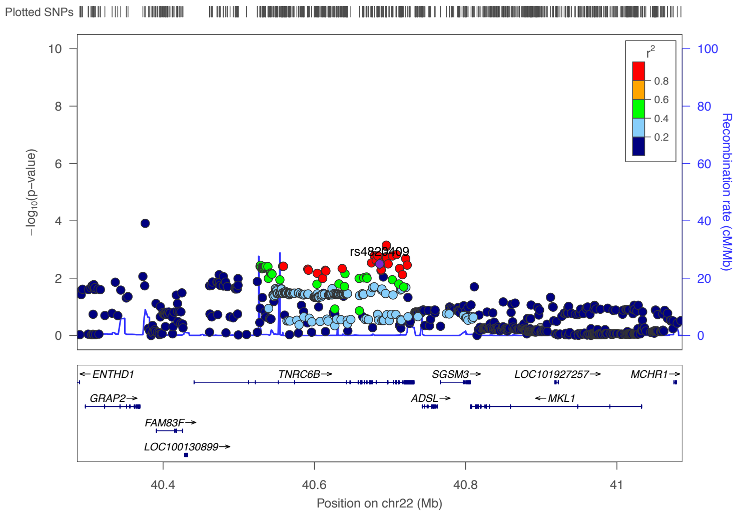


Tall people
